# A high-quality grapevine downy mildew genome assembly reveals rapidly evolving and lineage-specific putative host adaptation genes

**DOI:** 10.1101/350041

**Authors:** Yann Dussert, Isabelle D. Mazet, Carole Couture, Jérôme Gouzy, Marie-Christine Piron, Claire Kuchly, Olivier Bouchez, Claude Rispe, Pere Mestre, François Delmotte

## Abstract

Downy mildews are obligate biotrophic oomycete pathogens that cause devastating plant diseases on economically important crops. *Plasmopara viticola* is the causal agent of grapevine downy mildew, a major disease in vineyards worldwide. We sequenced the genome of *Pl. viticola* with PacBio long reads and obtained a new 92.94 Mb assembly with high continuity (359 scaffolds for a N50 of 706.5 kb) due to a better resolution of repeat regions. This assembly presented a high level of gene completeness, recovering 1,592 genes encoding secreted proteins involved in plant-pathogen interactions. *Pl. viticola* had a two-speed genome architecture, with secreted protein-encoding genes preferentially located in gene-sparse, repeat-rich regions and evolving rapidly, as indicated by pairwise dN/dS values. We also used short reads to assemble the genome of *Plasmopara muralis*, a closely related species infecting grape ivy (*Parthenocissus tricuspidata*). The lineage-specific proteins identified by comparative genomics analysis included a large proportion of RxLR cytoplasmic effectors and, more generally, genes with high dN/dS values. We identified 270 candidate genes under positive selection, including several genes encoding transporters and components of the RNA machinery potentially involved in host specialization. Finally, the *Pl. viticola* genome assembly generated here will allow the development of robust population genomics approaches for investigating the mechanisms involved in adaptation to biotic and abiotic selective pressures in this species.

**DATA AVAILABILITY:** Raw reads and genome assemblies have been deposited in GenBank (BioProjects PRJNA329579 for *Pl. viticola* and PRJNA448661 for *Pl. muralis*). Genome assemblies, gene annotations and analysis files (e.g. orthology relationships, full tables for GO enrichment analyses, pairwise dN/dS values and branch-site tests) have been deposited in Dataverse (*Pl. viticola* assembly and annotation: doi.org/10.15454/4NYHD6, *Pl. muralis* assembly and annotation: doi.org/10.15454/Q1QJYK, analysis files: doi.org/10.15454/8NZ8X9). Links to the data and information about the grapevine downy mildew genome project can be found at http://grapevine-downy-mildew-genome.com/.

## INTRODUCTION

Oomycetes are eukaryotic organisms with fungus-like characteristics. They belong to the Stramenopiles, which also include diatoms and brown algae. Oomycetes are distributed worldwide, and comprise marine, freshwater and terrestrial organisms, the most frequently studied of which are plant and animal pathogens (Thines & Kamoun 2010). These pathogens cause many diseases of economic and ecological importance. The most notorious examples include saprolegniosis in freshwater fishes, potato late blight, sudden oak death, white rust and downy mildew in a large number of plant species. Oomycetes have a large range of lifestyles, including obligate biotrophy (Thines & Kamoun 2010), a parasitic relationship in which the pathogen extracts nutrients from the living tissues of its host. An understanding of the virulence mechanisms of oomycete pathogens and of the ways in which they adapt to their hosts is the key to developing sustainable control strategies. Several oomycete genome sequences have already been published, for species from the genus *Saprolegnia* (Jiang et al. 2013), white rusts (McMullan et al. 2015; Kemen et al. 2011), the genus *Pythium* (Adhikari et al. 2013; Lévesque et al. 2010), the genus *Phytophthora* (Haas et al. 2009; Tyler et al. 2006; Lamour et al. 2012; Liu et al. 2016) and downy mildews (Baxter et al. 2010; Derevnina et al. 2015; Sharma et al. 2015). This abundant genomic data for oomycetes has provided opportunities to improve our understanding of the evolutionary processes linked to pathogenic lifestyles. For example, it has made it possible to identify genes lost following the adaptation to obligate biotrophy (Kemen et al. 2011; Baxter et al. 2010) and novel effector families involved in the manipulation of plant defenses (Bozkurt et al. 2012), and has given rise to the “two-speed genome” concept for filamentous plant pathogens (Dong et al. 2015).

*Plasmopara viticola*, an obligate biotrophic and heterothallic (i.e. sexual reproduction requires individuals with different mating types) oomycete, is the causal agent of downy mildew on grapevine (species of the Vitaceae family). It is endemic to North America and was introduced into Europe in the late 1870s (Fontaine et al. 2013). This pathogen is now found on cultivated grapevine (*Vitis vinifera*) worldwide, and downy mildew is considered to be one of the most important diseases of grapevine. *Pl. viticola* has recently been shown to form a complex of host-specialized species on wild *Vitis* in North America (Rouxel et al. 2013, 2014). The development of microsatellite markers for *Pl. viticola* over the last decade has led to improvements in our understanding of the genetic diversity, epidemiology (Gobbin et al. 2005, 2006) and invasion history of this species in Europe (Fontaine et al. 2013). *Pl. viticola* has a strong adaptive potential, as demonstrated by the rapid emergence of fungicide resistances in this species (reviewed in Delmas et al. 2017) and of the recent erosion of plant partial resistances (Delmas et al. 2016). Resources for population genetics studies based on genome-wide diversity are required to decipher the processes involved in this rapid adaptation, but such tools are currently lacking.

Upon infection, oomycetes secrete effector proteins, which modify host metabolism and defenses for the benefit of the pathogen. Effectors are described as apoplastic when they act in the extracellular space of the plant, and as cytoplasmic when translocated into the plant cell. RxLR effectors are the most widely studied class of cytoplasmic effectors (Bozkurt et al. 2012). Candidate apoplastic and cytoplasmic effectors have been described for *Pl. viticola* (Mestre et al. 2016, 2012; Xiang et al. 2017; Yin et al. 2015) and several genes encoding RxLR effectors have recently been functionally characterized (Xiang et al. 2016, 2017; Liu et al. 2018). However, the role of these proteins in infection and adaptation to the host remains unclear. Genomic resources for *Pl. viticola* are required to improve our understanding of the genetic mechanisms underlying plant-pathogen interactions in this pathosystem and their coevolution. Recent years have seen the publication of various transcriptomes (Mestre et al. 2016; Yin et al. 2015) and draft genome sequences based on short-read sequencing (Dussert et al. 2016; Brilli et al. 2018; Yin et al. 2017). Unfortunately, these assemblies remain incomplete and/or fragmented.

We produced a high-quality assembly of the *Pl. viticola* genome with PacBio long-read technology, which we used to characterize more extensively the repertoire of pathogenicity-related genes in this species. We also used short reads to assemble the first genome for *Plasmopara muralis*, a phylogenetically close species that infects members of the Vitaceae family such as *Parthenocissus quinquefolia* and *Parthenocissus tricuspidata*. We used these sequences and available oomycete genome sequences for a comparative genomics analysis, which identified gene families absent from all biotrophic oomycete species. Finally, we performed several tests to identify genes potentially under positive selection and involved in the adaptation of *Pl. viticola* and *Pl. muralis* to their hosts.

## MATERIALS AND METHODS

### Preparation of the material and DNA extraction

Sporangia from the sequenced *Pl. viticola* isolate (INRA-PV221) were previously collected for the assembly of a draft genome sequence (Dussert et al. 2016). This isolate belongs to clade B (*Pl. viticola* f. sp. *aestivalis*), as defined by Rouxel et al. (2013). Detached grapevine leaves from *Vitis vinifera* cv. Cabernet-Sauvignon plants were disinfected with 65% calcium hypochlorite (50 g/L), washed with sterilized water and infected with fresh sporangia (the inoculum also contained 5 g/L ampicillin, 1 g/L ticarcillin-clavulanate and 1 g/L boscalid). The plants were placed in a growth chamber for five to seven days at 22°C (12 h light/12 h dark photoperiod). We then collected sporangiophores and sporangia in TE buffer (10 mM Tris-HCl, 1 mM EDTA). The suspension was centrifuged (20,000 g, 4°C), the supernatant was removed and the pellet was stored at -80°C. Genomic DNA was extracted with a CTAB procedure, according to the protocol of Cheeseman et al. (2014), but with an additional ethanol precipitation step. The extracted DNA was quantified with a Qubit fluorometer (Invitrogen), and sequenced with a PacBio RS II (Pacific Biosciences) sequencer using P6-C4 chemistry (GeT-PlaGe facility, Toulouse, France).

*Plasmopara muralis* infects *Parthenocissus quinquefolia* in North America (Rouxel et al. 2014) and is also found on grape ivy (*Parthenocissus tricuspidata*) in Europe (Thines 2011). Sporangiophores and sporangia were collected from *Pl. muralis* isolate INRA-PM001 growing on the leaves of a *Parthenocissus tricuspidata* plant in Villenave d’Ornon, France. Genomic DNA was extracted with a standard CTAB protocol, as described by Delmotte et al. (2006), and sent to Beckman Coulter Genomics (Grenoble, France) for sequencing on an Illumina HiSeq 2000 sequencer (2x100 bp paired-end reads).

### RNA extraction & sequencing

Leaves of *Vitis vinifera* cv. Muscat Ottonel were inoculated with *Pl. viticola* isolate INRA-PV221, as described by Mestre et al. (2012). Infection was checked 24, 48 and 72 hours post-inoculation (hpi), by observation under a microscope, and leaves for these three time points were snap-frozen in liquid nitrogen and stored at -80°C. For each time point, three biological repetitions, with different batches of plants and inoculum, were performed. Total RNA was extracted by CTAB/phenol–chloroform extraction followed by LiCl precipitation (Zeng & Yang 2002). Residual genomic DNA was removed by DNase treatment with the Ambion Turbo DNA-free kit. RNA quality was checked with an Agilent BioAnalyzer. RNA-seq libraries were constructed from 4 μg of total RNA with the Illumina TruSeq kit. Sequencing was performed at the GeT-PlaGe facility (Toulouse, France) on an Illumina HiSeq 2000 sequencer (2x100 bp paired-end reads) for samples at 24 and 48 hpi and on an Illumina HiSeq 3000 sequencer (2x150 bp) for samples at 72 hpi, producing a mean of 38 million paired-end reads per sample.

### Genome assemblies

The PBcR wgs8.3rc2 assembly pipeline (Berlin et al. 2015) was used for read correction and assembly of the corrected reads. Corrected reads were also assembled with FALCON (Chin et al. 2016). Despite parameter tuning, the preliminary version of FALCON-Unzip (downloaded in April 2016) was unable to generate haplotypes and the FALCON assemblies were less comprehensive than our best PBcR assembly. However, the PBcR assembly was about twice the expected size of the genome due to the high degree of heterozygosity. We eliminated redundancy in the assembly with an ad-hoc procedure, and used the FALCON assembly to scaffold the PBcR contigs. The assembly was then polished with Pilon 1.20 (Walker et al. 2014), for which outputs were modified using custom scripts. Details of the assembly and polishing procedures are available in the Supplementary note.

The *Pl. muralis* genome was assembled with SPAdes 3.9.0 (Bankevich et al. 2012), using a custom procedure to discard contaminants (Supplementary note). Only scaffolds of more than 1 kb in length were retained. The median read coverage for the final assembly was 120x.

### Genome annotation

#### Repeat annotation

Repeated sequences in the two genomes were annotated with REPET 2.2 (Quesneville et al. 2005; Flutre et al. 2011): *de novo* libraries of repeated sequences were built with the TEdenovo pipeline (default parameters except minNbSeqPerGroup: 5 and no structural search), and genomes were annotated with the TEannot pipeline (default parameters, with the procedure recommended on the software website). Repeated elements from the Repbase20.05 database were also aligned to genome sequences during this step, but were used only to softmask sequences for gene prediction and not for other analyses.

#### Gene prediction

For *Pl. viticola*, RNA-seq reads were aligned against the genome with STAR 2.4 (Dobin et al. 2013) then assembled *de novo* into transcripts with Trinity release 20140717 (Grabherr et al. 2011), using a genome-guided approach. Genes were predicted with BRAKER 1.8 (Hoff et al. 2016), with the aligned RNA-seq reads, and MAKER 2.31 (Holt & Yandell 2011), with the assembled transcripts and proteomes of nine other oomycete species as evidence. For *Pl. muralis*, for which no RNA data were available, genes were predicted with MAKER, using gene predictor software trained with the *Pl. viticola* proteome. Details are available in the Supplementary note.

For the two predicted proteomes, we filtered out proteins with functional domains of transposable elements detected with InterProScan 5 (Jones et al. 2014) or for which hits were obtained with TransposonPSI (http://transposonpsi.sourceforge.net; 30% coverage, e-value < 1e^-10^).

### Quality control

The quality of genome assemblies and gene annotations was assessed with BUSCO 2.0 (Simão et al. 2015), using the Alveolata-Stramenopiles dataset. The *Pl. viticola* long-read assembly was aligned against the previously published draft assembly for the same isolate (Dussert et al. 2016) with the SynMap tool (Lyons et al. 2008) from CoGe (Lyons & Freeling 2008) with default parameters, and the results were visualized with Circos (Krzywinski et al. 2009). We also used QUAST 2.3 (Gurevich et al. 2013) to compare the annotated PacBio assembly (used as a reference) with the previous draft.

### Protein functional annotation and secretome characterization

Functional annotation of the proteome and Gene Ontology (GO) term mapping were performed with Blast2Go PRO 4.1.5 (Gotz et al. 2008), using results from BLASTP+ (-evalue 1e^-5^ -max_hsp 20 -num_alignment 20) with the NCBI nr database and from InterProScan.

For the identification of putatively secreted proteins, we first discarded proteins predicted by TargetP 1.1 (Emanuelsson et al. 2000) to have a mitochondrial targeting peptide (RC value between 1 and 3). The remaining proteins were analyzed with TMHMM 2.0 (Krogh et al. 2001), retaining only those with no transmembrane domain or a single transmembrane domain in the first 70 amino acids. Finally, SignalP 4 was used to detect signal peptides in the remaining proteins (Petersen et al. 2011).

One of the most widely families of secreted effectors in *Phytophthora* species and downy mildews has a conserved N-terminal RxLR (Arg-X-Leu-Arg) domain, often followed by a dEER (Asp-Glu-Glu-Arg) motif. We searched for candidate effector RxLR proteins in each secretome, using the Galaxy workflow of Cock et al. (2013): genes identified as putative effectors by the method described by Win et al. (2007) and either of the two methods described by Whisson et al. (2007) were retained. These candidates were used as queries in high-stringency (-evalue 1e^-30^) BLASTP searches against the respective *Plasmopara* proteomes. The sequences of the proteins identified as BLAST hits were aligned, and those with clear RxLR and/or dEER motifs at the expected positions were retained. In parallel, secretomes were used as queries in a low-stringency (-evalue 1e^-5^) BLASTP search against a database of oomycete RxLR effectors, and BLAST hits were considered as candidates. Finally, candidate effector genes were subjected to a structural homology analysis with Phyre2 (Kelley et al. 2015). Proteins with high-confidence (100%) structural homology to proteins with known functions but lacking RxLR and/or dEER motifs at the expected positions were considered to be false-positives and were discarded. The WY-motif, a conserved structural fold, is often found in RxLR proteins (Boutemy et al. 2011). We searched for this motif in secreted proteins with HMMER 3.1b2 (available at hmmer.org), using the HMM profile from Boutemy et al. (2011), with the same domain score cut-off of 0. For proteins with WY motifs not detected as putative RxLR effectors, amino-acid sequences were aligned with MAFFT 7 (Katoh & Standley 2013), and sequence logos from these alignments were generated using WebLogo 2.8.2 (Crooks et al. 2004).

The secretomes of the two species were tested for enrichment in GO terms for the “biological process” and “molecular function” ontologies with the topGO package (Alexa & Rahnenfuhrer 2016) in R (R Core Team 2017). We used Fisher’s exact test with the “elim” algorithm, which takes into account local dependencies between GO terms (Alexa et al. 2006). With this algorithm, tests for different GO terms are not independent, and a classical false discovery rate (FDR) (Storey & Tibshirani 2003) approach is not applicable. We followed the procedure of Daub et al. (2013), using modified R code from this paper, to calculate the empirical FDR with 200 permutations of the gene label (secreted / not secreted), and we reported GO terms with a FDR < 10%.

### Genome characteristics and architecture

The size of coding sequences, coding exons and introns, and the number of exons were determined with GenomeTools (Gremme et al. 2013). The RNA-seq read count for *Pl. viticola* genes in each library was calculated with htseq-count from HTseq 0.6.1p1 (Anders et al. 2015), and read counts per million (CPM) were computed with R. A gene was considered to be supported by RNA data if its CPM value was at 1 or more in at least one biological replicate. Depending on the library, only 0.03% to 4.04% of reads were mapped onto the genome, so we decided not to perform differential gene expression analysis with these data. For *Pl. viticola*, Illumina paired-end reads from a previous genome draft (and used for sequence correction, see Supplementary note) were aligned against the new assembly with BWA (Li & Durbin 2009). Heterozygous sites were then detected with VARSCAN 2.3 (Koboldt et al. 2012) (--min-coverage 20 --min-reads2 10 --min-avg-qual 15 --min-var-freq 0.2 --p-value 0.01). Sites with a read coverage more than 1.5 times the mean genome coverage were discarded. Using BEDtools 2.25 (Quinlan & Hall 2010), we determined the proportion of heterozygous sites, gene density, repeat coverage and GC content for genomic windows of 100 kb, and visualized the results with Circos. This analysis was not carried out with *Pl. muralis*, because the genome assembly was too fragmented.

The genome architecture of the two species was characterized by assessing local gene density (Saunders et al. 2014). For each gene, the 5’ and 3’ intergenic distances were calculated, and the gene count for two-dimensional distance bins was plotted as a heatmap in R. Genes located at the scaffold extremities were not taken into account for this analysis. The distance between genes and their closest repeat sequence (not allowing for overlaps) was calculated with BEDtools. We investigated the possible relationship between intergenic distance and the orientation of adjacent gene pairs, by classifying adjacent gene pairs as oriented head-to-head (HH, adjacent pairs oriented ← →), tail-to-tail (TT, → ←) or tail-to-head (TH, → → or ← ←). The observed proportions of gene pairs in the different orientations were compared with the proportions expected by chance (0.25 for HH and TT, 0.50 for TH) using a chi-square test performed in R.

### Comparative genomics with other oomycetes

Gene synteny between the genomes of *Pl. viticola* and *Pl. muralis, Pl. halstedii, Ph. infestans* and *Py. ultimum* was assessed with SynMap (merging of syntenic blocks with Quota Align Merge, syntenic depth 1:1) from CoGe, and visualized with Circos. Orthology relationships for the genes of *Pl. viticola, Pl. muralis* and 19 other oomycetes from the orders Peronosporales, Pythiales, Albuginales, and Saprolegniales and including five other obligate biotrophic species (Supplementary table 1) were determined with OrthoFinder 1.1.2 (Emms & Kelly 2015). The results were visualized with the R package UpSetR (Conway et al. 2017) for various species groups. GO enrichment analyses were carried out for genes absent from all biotrophs as described above, using the GO annotation for nine species available from the UniProt database (Supplementary table 1). The Kyoto Encyclopedia of Genes and Genomes (KEGG) annotation of the *Ph. infestans* proteome, available through the KEGG API, was also used to predict the function of orthologs missing from all biotrophs and the biological pathways they are involved into.

The amino-acid sequences of single-copy genes detected in all oomycetes in the orthology analysis were aligned with MAFFT 7. Alignments were cleaned with GBlocks (Castresana 2000) with default parameters and concatenated, discarding protein alignments extending over less than 50 positions. The concatenated alignment was used to infer a phylogeny for the species considered with PhyML 3 (Guindon et al. 2010), available from http://www.atgc-montpellier.fr/phyml-sms, with automatic selection of the substitution model with SMS (Lefort et al. 2017) and 100 bootstraps.

### Detection of positive selection

To detect genes with a high substitution rate in *Pl. viticola* and *Pl. muralis*, we computed pairwise dN/dS values between pairs of orthologs from the two species. Pairs of orthologs were identified by the reciprocal best hits (RBH) method with BLASTP+ (-evalue 1e^-6^ - qcov_hsp_perc 50, min. identity 40%) and using the Smith-Waterman algorithm, as suggested by Moreno-Hagelsieb & Latimer (2008). The nucleotide sequences of RBH pairs were aligned with MUSCLE 3.8.31 (Edgar 2004), using protein alignments as guides, and poorly aligned regions were removed with GBlocks, trimming only entire codons (-t=c -b3=6 -b4=9). Alignments of less than 120 bp in length were discarded, and pairwise dN/dS values were calculated with codeml (runmode=-2) in PAML 4.8 (Yang 2007). Pairs of orthologs with dS values greater than 3 and dN/dS values of 99 were discarded (final dataset: 9,053 pairs of orthologs). Mean pairwise dN/dS values for different gene categories (e.g. non-secreted and secreted genes, or core and specific genes) were compared in *t* tests, assuming unequal variance, in R.

The pairwise dN/dS approach makes it possible to analyze large numbers of pairs of orthologs and to include genes specific to *Pl. viticola* and *Pl. muralis*, but has some drawbacks. First, focusing on gene pairs with a pairwise dN/dS > 1 as a signature of positive selection is not a very sensitive approach, because pairwise dN/dS values provide an average estimate of selective pressure over the whole gene. Indeed, whereas most sites in most proteins are probably subject to strong evolutionary constraints, some proteins may have specific regions or sites under positive selection that remain undetected in the pairwise approach because their overall ratio is below 1. In addition, a high pairwise dN/dS value may indicate that the protein evolved rapidly in one or both species. We obtained a more precise assessment of rate changes in the various evolutionary branches and between sites, by carrying out a branch-site test (Zhang et al. 2005) to detect positive selection affecting a subset of sites in a particular lineage. This analysis was performed with data from six species: *Pl. viticola, Pl. muralis, Pl. halstedii, Ph. infestans, Ph. nicotianae* and *Ph. capsici*. Orthology relationships for the genes of the six species were determined with OrthoFinder, retaining only single-copy orthologs (resulting in 4,042 ortholog groups). For each ortholog group, nucleotide sequences were aligned and cleaned as described for the pairwise dN/dS computation, resulting in a final data set of 4,017 ortholog groups. Positive selection was tested separately for the *Pl. viticola* and *Pl. muralis* branches. The likelihoods of the null (no selection) and alternative (positive selection) models were calculated with codeml (model=2, NSsites=2). For the alternative model, three runs with different starting values for omega (0.5, 1 and 1.5) were performed, and the run with the highest likelihood was retained. The two models were compared in a likelihood-ratio test (LRT) in R, with the test statistic following a chi-square distribution with 1 degree of freedom. *P*-values were corrected for multiple testing by the FDR procedure in the R package qvalue (Dabney et al. 2015), and we retained genes with a FDR < 10%. GO enrichment analyses for genes with a significant positive selection signal were performed as described above.

## RESULTS AND DISCUSSION

### A new high-quality *Pl. viticola* genome assembly

The PacBio sequencing of *Pl. viticola* isolate INRA-PV221 produced 2.6 million reads (mean size: 8,072 bp, total data: 21.2 Gb). The genome size of *Pl. viticola* has been estimated at about 115 Mb (Voglmayr & Greilhuber 1998), so the amount of data obtained corresponded to a mean sequencing depth of 185x. The use of long reads at this coverage made it possible to obtain a high-quality assembly (Table 1), with only 358 scaffolds and a total size of 92.94 Mb. This assembly is more complete (95.7% of expected BUSCO genes, Alveolata-Stramenopiles dataset, Supplementary table 2) than previous drafts generated with short reads only (Dussert et al. 2016; Brilli et al. 2018) or with a hybrid approach in which long reads were used for scaffolding (Yin et al. 2017). The assembly was also more continuous, resolving many of the repetitive regions (Supplementary figure 1). Heterozygosity levels were high in the sequenced *Pl. viticola* isolate, with a mean of 7,952 heterozygous sites per Mb. The assembly of highly heterozygous diploid species is often problematic. However, the percentage of duplicated genes in our assembly was lower than that for previous drafts, with only 1.7% of BUSCO genes duplicated, indicating that our procedure for eliminating alternative haplotypes was efficient. The higher heterozygosity of *Pl. viticola* than of the homothallic *Pl. halstedii* (120 heterozygous sites per Mb, Sharma et al. 2015) was probably due to its heterothallism, which strongly favors outcrossing. Repeat sequences accounted for 37.34% of the genome assembly for *Pl. viticola*. Local repeat coverage varied considerably, from 0 to 100% in our 100 kb window analysis (Figure 1). Most (78.5%) of the identified repeat sequences were LTR retrotransposons and their non-autonomous derivatives (Supplementary table 3). In total, 15,960 protein-coding genes were predicted for *Pl. viticola*, 75.6% of which were supported by RNA-seq data. The completeness of the annotation was high, with 97.4% of the expected BUSCO proteins (Table 1). Almost half the genes (46.7%) consisted of a single exon (Table 1). As observed for repeat coverage, gene density in *Pl. viticola* varied considerably over the genome, from 0 to 53 genes per 100 kb window (Figure 1).

**Table 1.** Assembly and annotation statistics for the genomes of *Pl. viticola* and *Pl. muralis*.

|  | *Pl. viticola* | *Pl. muralis* |
| --- | --- | --- |
| *Assembly* |  |  |
| Assembly size (Mb) | 92.94 | 59.54 |
| Number of scaffolds | 358 | 1,437 |
| N50 (size in kb / count) | 706.5 / 38 | 184.5 / 101 |
| Max. scaffold size (Mb) | 2.85 | 1.03 |
| % GC | 44.85 | 44.29 |
| % of repeat elements | 37.34 | 10.43 |
| BUSCO genome [S / D / F] (%) | 95.7 [93.6 / 1.7 / 0.4] | 95.8 [94.9 / 0.0 / 0.9] |
| *Gene annotation* |  |  |
| Number of genes | 15,960 | 12,194 |
| BUSCO proteome [S / D / F] (%) | 97.4 [94.0 / 3.0 / 0.4] | 93.6 [ 93.2 / 0.0 / 0.4] |
| Median coding sequence length [range] (bp) | 957 [105 - 40,497] | 999 [99 - 41,409] |
| Median exon length [range] (bp) | 237 [1 - 14,776] | 207 [1 - 14,941] |
| Median exon number [range] (bp) | 2 [1 - 33] | 2 [1 - 35] |
| Median intron length [range] (bp) | 67 [4 - 3288] | 66 [4 - 3991] |
| Number of single exon genes | 7,454 | 4,602 |

BUSCO analyses have been carried out with the Alveolata-Stramenopiles data set. S: single copy, D: duplicated, F: fragmented genes.

**Figure 1.**
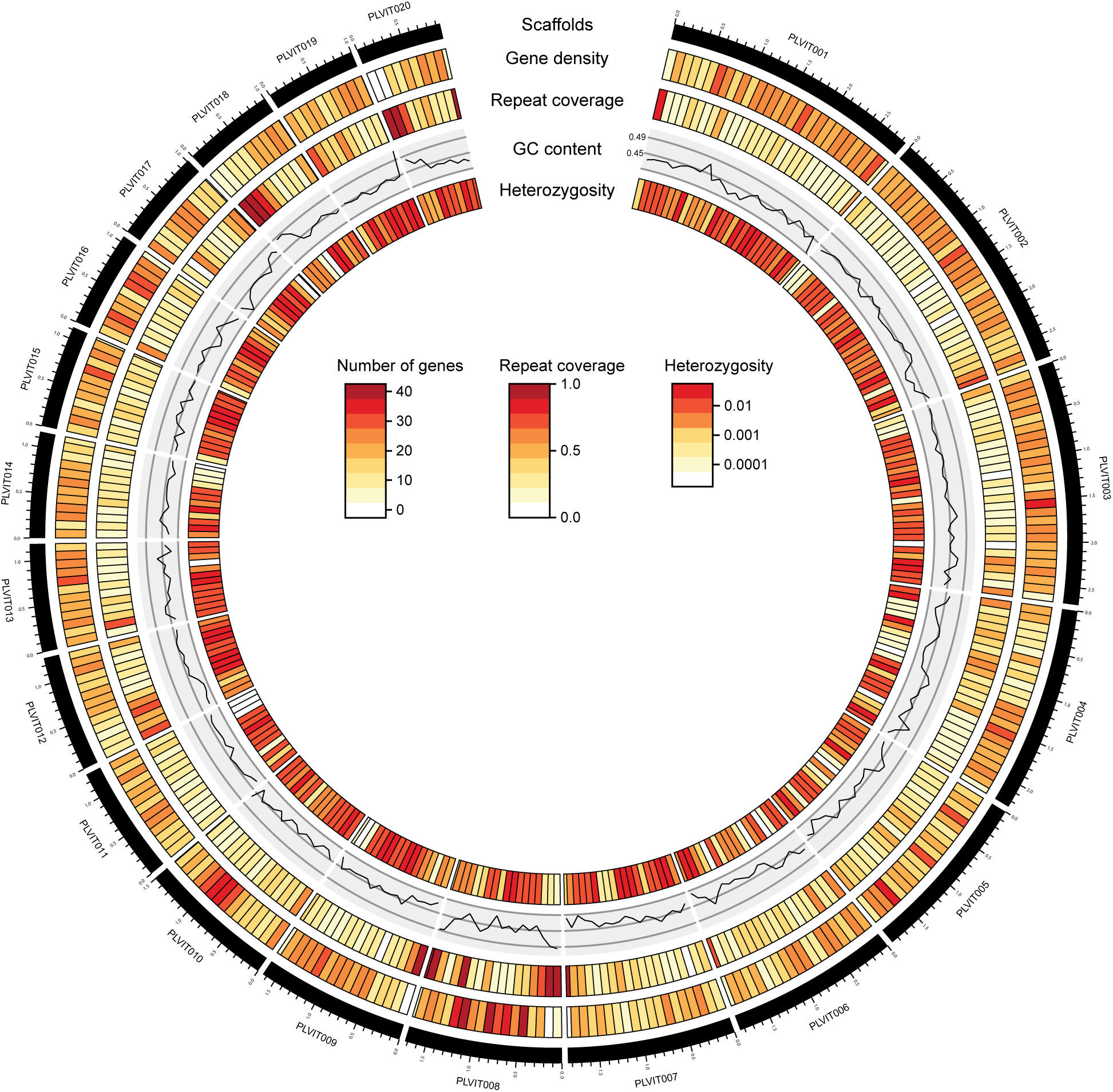
Genomic architecture of *Plasmopara viticola*. Gene density (number of genes per window), repeat coverage (proportion of the window covered by repeated sequences), GC content and the percentage of heterozygous sites are represented for genomic windows of 100 kb for the 20 largest scaffolds of the assembly. Tick marks on scaffolds represent 100 kb. The percentage of heterozygous sites is shown on a log scale.

The assembly of the *Pl. muralis* genome with short reads resulted in 1,437 scaffolds, for a total size of 59.54 Mb (Table 1). Estimated genome sizes in *Plasmopara* species range from 90 to 160 Mb (Voglmayr & Greilhuber 1998), so the smaller assembly size suggested that regions of the genome might be missing. However, the percentage of genes expected to be conserved was high in the assembly (95.8% of BUSCO genes), indicating a high level of completeness. Only 10.43% of the assembly consisted of repeat elements, suggesting that most of the repeated regions were not assembled, probably due to the exclusive use of paired-end reads with a short insert size. As in *Pl. viticola*, most (62.4%) of the identified repeat sequences were retrotransposons of the LTR order and their derivatives (Supplementary table 3). We annotated 12,194 protein-coding genes in the genome of *Pl. muralis*. The completeness of the annotation was again high, with 93.6% of expected BUSCO proteins (Table 1). As observed for *Pl. viticola*, a large proportion of genes (37.7%) in *Pl. muralis* were composed of single exons.

### Gene losses linked to biotrophic lifestyle in oomycetes

The *Pl. viticola* genome displayed a high degree of gene synteny with the genomes of *Pl. muralis, Ph. infestans* and *Py. ultimum*. Depending on the species used for comparison, the mean size of the syntenic blocks in *Pl. viticola* ranged from 130 to 172 kb, with syntenic regions covering a total of 37.7 to 43.5% of the genome (Supplementary table 4). We observed a very low level of synteny between *Pl. viticola* and *Pl. halstedii*, which was surprising since given the closeness of the phylogenetic relationship between these species: the mean syntenic block size was 38 kb, and syntenic regions accounted for only 10.2% of the genome. Moreover, syntenic blocks were not in the same order in *Pl. viticola* and *Pl. halstedii*, whereas their organization was similar in *Pl. viticola* and the other species (Supplementary figure 2). This suggested that either the genome of *Pl. halstedii* had undergone important genomic rearrangements, or alternatively, could indicate the existence of missassemblies in the *Pl. halstedii* reference genome.

The orthology analysis detected 18,115 ortholog groups, 2,474 of which were common to all oomycete species considered (Figure 2A), including 163 single-copy orthologs. The phylogeny based on these single-copy orthologs (144 single-copy orthologs after filtering, for a total of 41,428 amino-acid positions) suggested that downy mildews belonging to the Peronosporales (*Plasmopara* genus, *Pe. tabacina* and *Hy. arabidopsidis*) were paraphyletic and, consequently, that obligate biotrophy had evolved independently at least twice within this order (Figure 2B, Supplementary figure 3). These results confirmed other recent phylogenetic analyses (McCarthy & Fitzpatrick 2017; Sharma et al. 2015; Bourret et al. 2018) contradicting earlier evidence suggesting that downy mildews are monophyletic (Göker et al. 2007). However, the number of oomycete species for which genomic sequence data are available remains small, and this result should be reexamined when more species, notably from the genus *Phytophthora*, have been sequenced.

**Figure 2.**
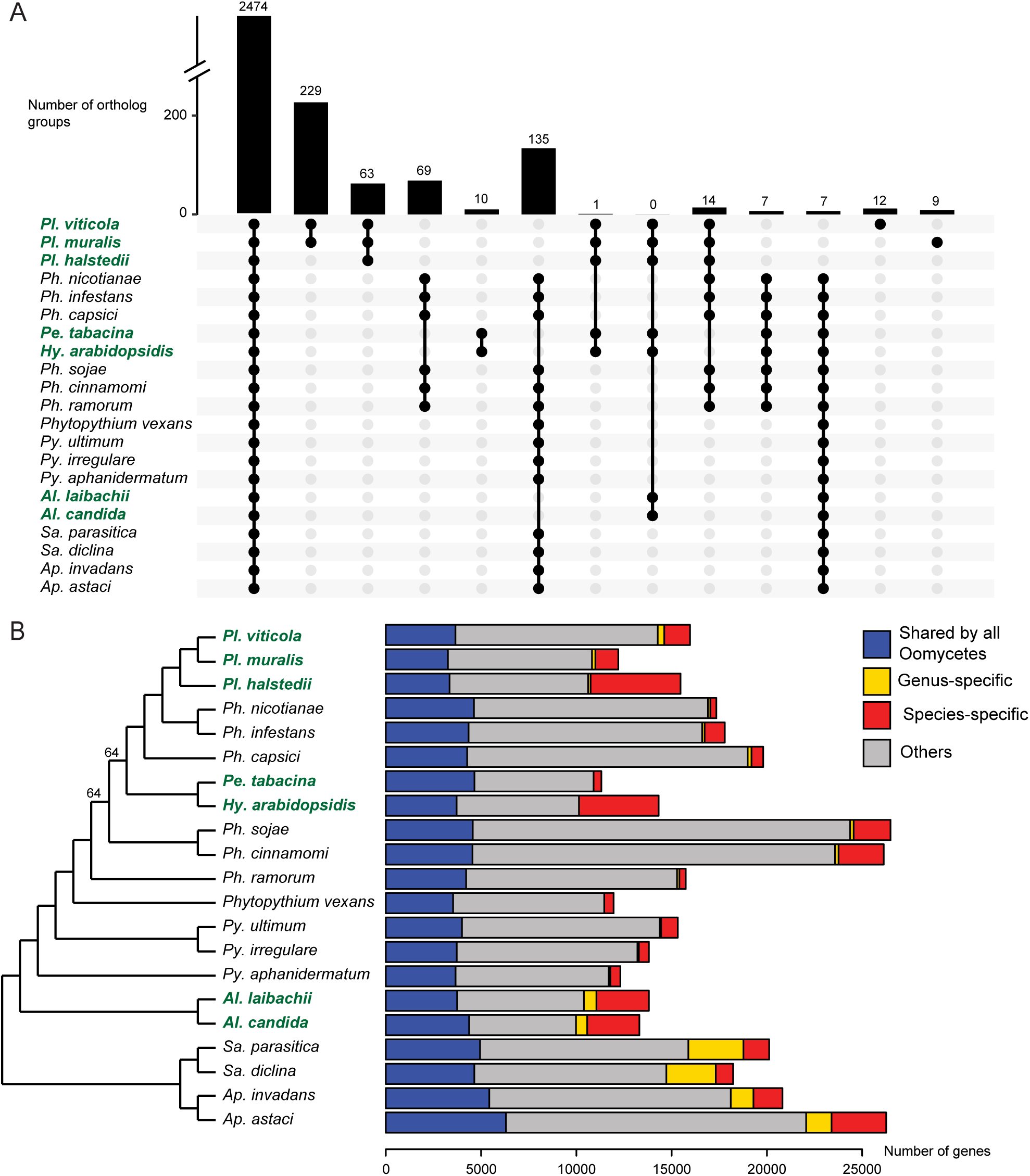
Comparative analysis of 21 oomycete proteomes. A: UpSet plot of the orthology analysis. Bars represent the number of ortholog groups common to different species denoted by black dots. B: Phylogenetic relationships and number of genes common to or specific to oomycete species. The maximum likelihood species tree (left) was built with 144 single-copy orthologs. All nodes had a bootstrap support of 100% unless otherwise indicated. The species tree with branch lengths is available in Supplementary figure 3. The bars on the right represent the numbers of genes in different categories. For both figures, biotrophic species names are shown in bold green typeface.

Previous comparative studies led to the discovery of genes lost following the adaptation to biotrophy in particular oomycete species (Baxter et al. 2010; Kemen et al. 2011). We searched for gene families common to or lost in all oomycete biotrophic species. Biotrophic species had no specific ortholog group in common, and only one group of orthologs (containing proteins of unknown function) was common to the biotrophic species of the Peronosporales order (*Plasmopara* sp., *Pe. tabacina* and *Hy. arabidopsidis*), consistent with the notion that the two lineages of biotrophic species evolved independently. We identified 135 groups of orthologs absent exclusively in biotrophic species (Figure 2A), corresponding to gene losses common to all biotrophs. While some of these gene losses might be random, they also could be due to convergent adaptive processes. Some of the GO terms for which enrichment was detected in these ortholog groups (Supplementary table 5) were totally absent from biotrophic oomycete proteomes. This was the case for proteins with riboflavin (vitamin B2) transport activity. Riboflavin is the precursor of the FAD and FMN cofactors, found in flavoproteins, and a significant depletion of GO terms characterizing flavoproteins (“FAD binding”, “FMN binding“, “D-amino-acid oxidase activity” and “succinate dehydrogenase activity“) was also observed in biotrophs. Overall, biotrophic species had smaller numbers of flavoproteins, with only 17 on average, versus 57 in other species (Supplementary table 6). Cobalamin (vitamin B12, another cofactor) binding proteins, such as methylmalonyl-CoA mutase and the cobalamin-dependent methionine synthase, were also totally absent from biotrophic species. Furthermore, one of the groups of orthologs absent from biotrophic species contained proteins involved in cobalamin metabolism. A possible explanation for the absence of these genes is that biotrophic species might not be able to obtain these cofactors or their precursors easily from the host, and they have therefore lost genes encoding proteins requiring these cofactors.

Only 15 of the *Ph. infestans* orthologs missing in biotrophic species were KEGG-annotated (Supplementary table 7), and most of their functions were found in the GO enrichment analysis. Among them were two proteins (PITG_13128 and PITG_21426) from the sterol biosynthesis pathway. These genes are expressed in *Ph. infestans*, but their actual function is unknown since they do not seem to be involved in sterol modification (Dahlin et al. 2017). The protein from the pyrimidine metabolism pathway (PITG_02105) may be involved in the pyrimidine salvage pathway during the necrotrophic phase (García-Bayona et al. 2014), which might explain why it has been lost in biotrophic oomycetes.

The GO terms depleted in biotrophic species also included “proteolysis“, which encompassed two ortholog groups including proteins with PAN/Apple domains, several of which had a secretion peptide. These genes may have been lost to facilitate the evasion of host recognition, because PAN/Apple domains, which bind polysaccharides or proteins, can induce plant defenses (Larroque et al. 2012). The proteins annotated with the terms DNA binding/DNA recombination/DNA integration probably constituted two large transposable element families, surprisingly absent from all biotrophic oomycetes. Finally, the missing groups of orthologs included a large family of transmembrane amino-acid transporters, two ortholog groups of ubiquitin-protein ligases, one ortholog group of patatin-like phospholipases (a function also found in the *Ph. infestans* KEGG annotation) and one group of secreted peroxiredoxins. These gene families may be involved in specific metabolic pathways that have been lost in biotrophic oomycetes, but the reasons for this loss remain unclear because the precise functions of these proteins are unknown.

### Repertoire of pathogenicity-related proteins in *Pl. viticola* and *Pl. muralis*

Plant pathogens secrete proteins to facilitate successful host colonization and growth. These proteins may remain in the apoplastic region or are delivered to the cytoplasm of plant cells. The proteome of *Pl. viticola* contained 1,592 putatively secreted proteins, 1,244 of which were supported by RNA-seq data. This corresponded to about 2.5 times as many secreted proteins as were found in *Pl. halstedii* (631), but was similar to the numbers found in *Phytophthora* species (see Sharma et al. 2015 for review). The number of secreted proteins in the *Pl. muralis* proteome (720) was closer to that for *Pl. halstedii*. Differences in secretome size may be due at least partly to differences in assembly completeness. Indeed, a comparison of our previous draft with the long-read assembly with QUAST revealed that 4,477 genes, including 655 encoding secreted proteins, were missing from the INRA-PV221 short-read assembly. Thus, the PacBio assembly recovered a larger number of genes, including a considerable number of genes encoding secreted proteins. The secretomes of the two species included a large proportion of candidate pathogenicity-related genes (Table 2) putatively involved in plant-pathogen interactions. Furthermore, GO enrichment analysis (Supplementary tables 8 and 9) showed that these secretomes were enriched in functions linked to plant cell wall modifications (degradation and modification of cellulose and pectin), proteolysis, protease inhibition and reactive oxygen species metabolism, which are involved in plant defenses.

**Table 2.** Putative pathogenicity-related proteins found in *Pl. viticola* and *Pl. muralis*.

|  | *Pl. viticola* | *Pl. muralis* |
| --- | --- | --- |
| Aspartic peptidases | 37 (2) | 19 (2) |
| Cutinases | 2 (1) | 2 (2) |
| Cysteine proteases | 54 (11) | 40 (8) |
| Cysteine-rich proteins | 22 (16) | 21 (14) |
| Elicitins | 19 (14) | 20 (14) |
| Glycosyl hydrolases | 195 (118) | 115 (53) |
| NPP1-like proteins | 10 (10) | 7 (5) |
| Pectate lyases | 25 (19) | 5 (3) |
| Pectin esterases | 21 (15) | 14 (5) |
| Protease inhibitors | 20 (13) | 15 (8) |
| Serine proteases | 161 (79) | 97 (27) |
| RxLRs | 540 | 143 |
| with WY motif | 317 | 66 |
| WY motif, no RxLR domain | 26 | 12 |

Number for proteins with a secretion peptide are in brackets except for RxLRs, which all have a secretion peptide.

RxLR proteins, a class of cytoplasmic effectors (i.e. translocated into the plant cell), accounted for a large proportion of the proteins in the secretomes. The secreted proteins included 540 proteins from *Pl. viticola* and 132 proteins from *Pl. muralis* (33.9% and 18.3% of the secreted proteins, respectively) identified as candidate RxLR effector genes by at least one method (Supplementary table 10). Phyre2 analysis revealed that a large proportion of the candidates identified by BLAST also displayed high-confidence structural homology to known RxLR effectors. The number of putative RxLRs with RNA-seq support in *Pl. viticola* (484) was also high. There were 343 secreted proteins (including 317 RxLRs) with at least one WY motif in *Pl. viticola*, and 78 (66 RxLRs) in *Pl. muralis*. The number of WY domains in a given protein ranged from 1 to 17 (median: 3). The sequences of secreted proteins with WY motifs not annotated as RxLR contained a dEER domain (Supplementary figure 4), indicating that they might also act as RxLR-like effectors.

Genes specific to a lineage are often involved in species-specific adaptive processes (Khalturin et al. 2009). We therefore searched for genes restricted to *Plasmopara* species infecting hosts from the Vitaceae. Orthology analysis identified 229 ortholog groups (comprising 691 *Pl. viticola* genes and 483 *Pl. muralis* genes, for a total of 1,174 genes) common only to *Pl. viticola* and *Pl. muralis* (Figure 2A). Secreted proteins accounted for a large proportion (32.7%) of these proteins, probably due to their role in pathogen-host interactions. Most of these secreted proteins (61.7%, distributed in 39 ortholog groups) were annotated as RxLR, suggesting that adaptation to hosts from the Vitaceae probably involved diversification of this effector family. Specific groups also included 17 secreted serine proteases (in one group), two glycosyl hydrolases and one necrosis-inducing NPP1-like protein. Another 1,347 genes were specific to *Pl. viticola*, including 151 secreted proteins, 28 of which were annotated as RxLRs. *Pl. viticola*-specific proteins also included two secreted cysteine proteases, three glycosyl hydrolases, three pectin esterases and one serine protease. There were 1,194 proteins (40 secreted) specific to *Pl. muralis*, including four RxLRs. Other interesting proteins included two glycosyl hydrolases (one of which was secreted) and one serine protease. Most of the genes specific to *Pl. viticola* or *Pl. muralis* were not functionally annotated, with only 21.9% and 11.6%, respectively, of these genes having at least one mapped GO term.

### A two-speed genome architecture

*Pl. viticola* and *Pl. muralis* displayed the two-speed genome architecture (Figure 3, Supplementary figure 5) observed in several filamentous plant pathogens (Dong et al. 2015). Core genes common to all oomycetes were preferentially located in gene-dense regions (Figure 3B), whereas genes encoding secreted proteins in *Pl. viticola* and *Pl. muralis* had larger flanking intergenic distances than other genes, indicating that they were preferentially located in gene-sparse regions (Figure 3C). This was also the case for genes specific to *Pl. viticola* and *Pl. muralis* (Figure 3D). Gene-sparse regions coincided with repeat-rich regions in *Pl. viticola*, as the distance between genes and their closest repeat sequence was shorter in these regions (Figure 3E). It was not clear whether this was also the case in *Pl. muralis* (Supplementary figure 5E), probably because, as previously discussed, most of the repeat regions were not present in the assembly.

**Figure 3.**
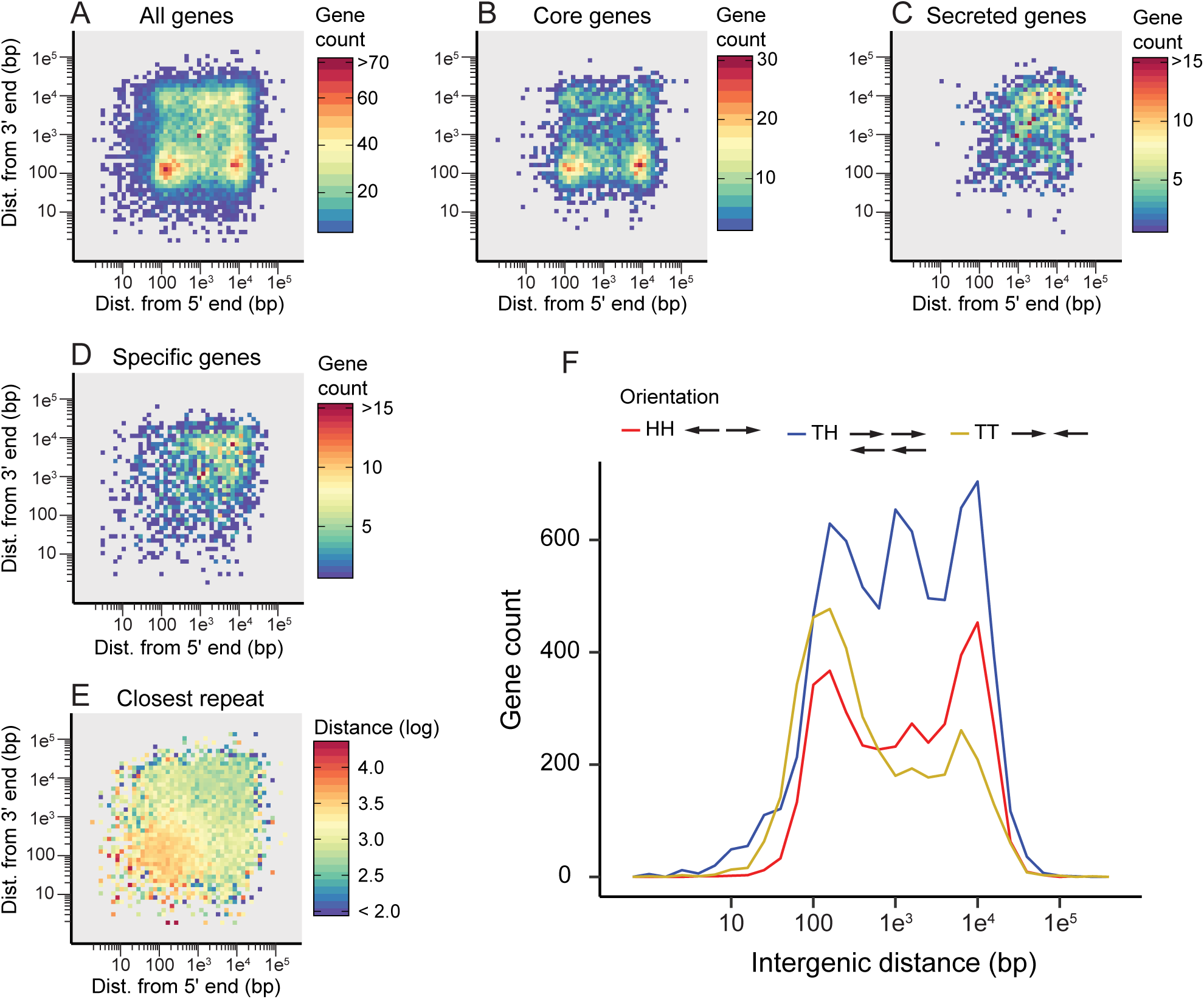
Local gene density and organization in *Plasmopara viticola*. Local gene density when considering (A) all genes, (B) core genes found in ortholog groups common to all oomycetes, (C) genes encoding secreted proteins and (D) genes found only in *Pl. viticola* or in both *Pl. viticola* and *Pl. muralis*. The number of genes for two-dimensional (5’ and 3’) intergenic distance bins is represented as a heatmap. (E) Mean distance between genes their closest repeat element, represented as a heatmap for two-dimensional intergenic distance bins. (F) Orientation and intergenic distance of adjacent gene pairs. The gene count for different intergenic distances has been plotted for adjacent gene pairs with a head-to-head (HH), tail-to-head (TH) and tail-to-tail (TT) orientation. Distances are shown on a log scale for all figures

The distribution of adjacent gene-pair orientations was unbalanced for both *Pl. viticola* and *Pl. muralis* (Figure 3F, Supplementary figure 5F). Two major groups were identified when considering all genes or only core genes with short 3’ intergenic distance: genes with short (between 80 and 300 bp) or large (between 5 and 20 kb, approximately) 5’ intergenic distances (Figure 3A and 3B). This pattern was explained by an excess of gene pairs in a head-to-head (HH) orientation when considering intergenic distances > 5 kb (32 and 37% of HH gene pairs for *Pl. viti*cola and *Pl. muralis* respectively, p < 1e^-15^), and an excess of gene pairs in a tail-to-tail (TT) orientation for intergenic distances < 500 bp (34 and 38% of TT gene pairs, p < 1e^-15^). Thus, gene pairs with a TT orientation, which had a shared 3’ intergenic region, were, on average, closer to each other than gene pairs with a HH orientation, which had a shared 5’ intergenic region. Neighboring genes generally have correlated expression levels in eukaryotic genomes (Michalak 2008), but tail-to-tail gene pairs are more frequently negatively co-regulated in *Arabidopsis thaliana* (Williams & Bowles 2004) and *Saccharomyces cerevisiae* (Wang et al. 2014) than adjacent genes in head-to-head and tail-to-head orientations. Unfortunately, we had too little RNA-seq data (not enough mapped reads, see the Materials & Methods section) to determine whether this was the case in *Pl. viticola*. In *Ph. infestans*, about 10% of adjacent genes have (positively or negatively) correlated expression levels (Roy et al. 2013), but the specific effect of different gene-pair orientations on gene expression has not been investigated in oomycetes.

### Rates of evolution of *Pl. viticola* and *Pl. muralis* genes

Pairwise dN/dS values for *Pl. viticola* and *Pl. muralis* orthologs (Figure 4A) were significantly higher (*p* < 10^-15^) for genes encoding secreted proteins (mean: 0.44) than for other genes (mean: 0.27). For RxLRs, the mean pairwise dN/dS was 0.70 (*p* < 10^-15^), indicating an even faster rate of evolution. This pattern was also observed when comparing core genes (mean: 0.20) and genes specific to species infecting hosts from the Vitaceae (mean: 0.79, *p* < 10^-15^), possibly due, at least in part, to relaxed selective constraints on lineage-specific genes (Cai & Petrov 2010). Overall, 92 pairs of orthologs had pairwise dN/dS values above 1 (Supplementary table 11), 64 of which were found expressed in our RNA-seq data in *Pl. viticola*. They included 37 genes encoding proteins secreted in at least one of the species, including 19 RxLRs, two serine proteases, one Kazal-like protease inhibitor, one cysteine-rich protein, one glycosyl hydrolase, and one mannose-binding lectin identified in a previous study (Mestre et al. 2012). All these genes encoding for secreted proteins were expressed in *Pl. viticola* except for PVIT_0017527.T1. Two of the candidate RxLRs had best BLASTP hits in the nr database with previously characterized *Pl. viticola* genes: PVIT_0013263.T1 displayed 99.6% amino-acid identity with RxLR16, which triggers a hypersensitive response in *Nicotiana benthamiana* (Xiang et al. 2017), and PVIT_0010026.T1 displayed 91.6% identity with RxLR28, which suppresses plant defenses in the same species (Xiang et al. 2016). Genes with dN/dS values above 1 were preferentially located in gene-sparse regions of the genome (Figure 4B), consistent with the notion that genes in these regions tend to evolve faster and are involved in adaptive processes.

**Figure 4.**
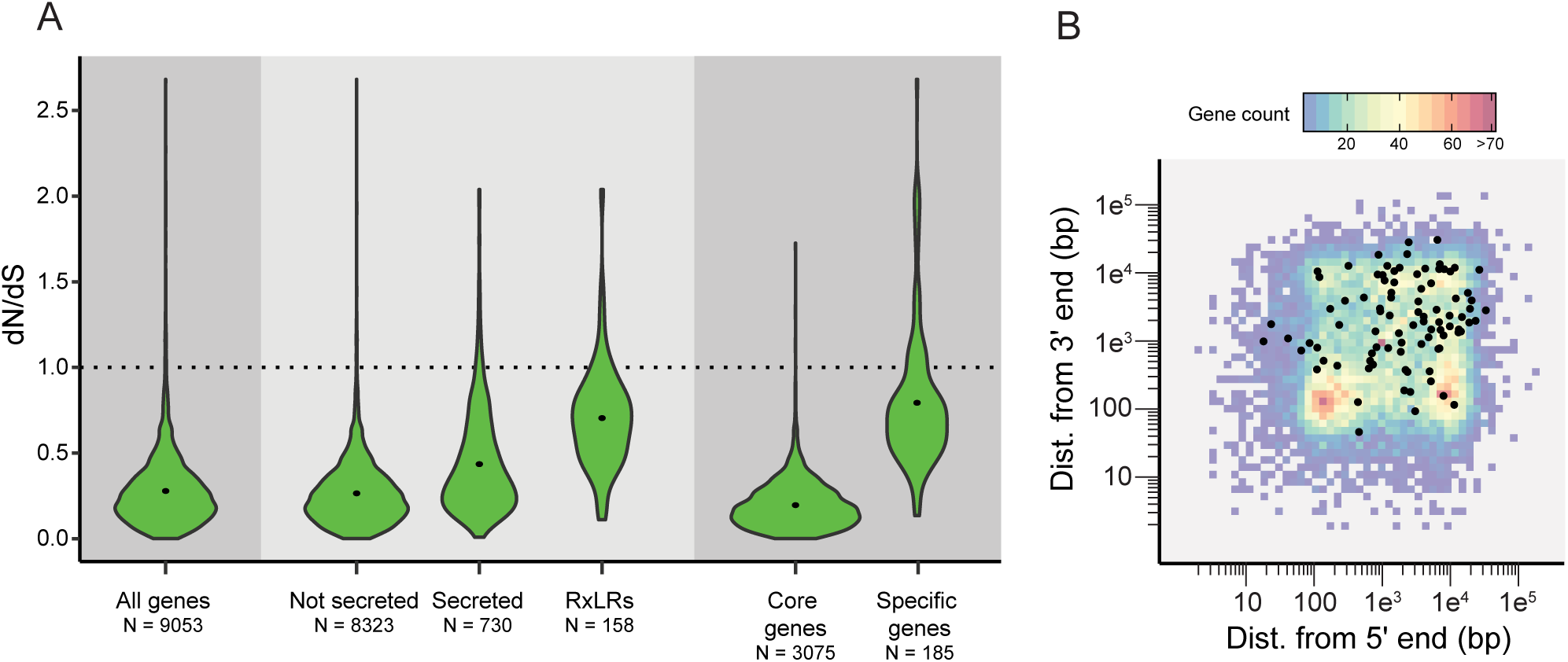
Evolution rates for ortholog pairs. (A) Pairwise dN/dS values for *Plasmopara viticola* and *Plasmopara muralis* orthologs, for different categories of genes. Core genes are genes found in ortholog groups common to all oomycetes and specific genes are found in ortholog groups common only to *Pl. viticola* and *Pl. muralis*. N: number of ortholog pairs. (B) Local gene density in *Pl. viticola* for all genes (heatmap) and for genes with a dN/dS above 1 (black dots).

### Genes under positive selection in *Pl. viticola* and *Pl. muralis*

The branch-site test was used to detect positive selection in *Pl. viticola* and *Pl. muralis*. This test had a relatively low power given that data were available for only six species, but it was nevertheless more sensitive than the pairwise dN/dS approach. In total, 270 single-copy orthologs (6.72% of the 4,017 gene families tested) were found to be under positive selection in at least one of the branches tested in branch-site tests (10% FDR threshold). The overall proportion of genes encoding secreted proteins with positive selection signals was low (Table 3). For the *Pl. viticola* branch, 131 significant genes were identified (Supplementary table 12), all of which were found expressed in the RNA-seq data except for PVIT_0009280.T1. Only three of the genes were encoding secreted proteins (2.54% of the tested secreted proteins). These proteins included a cysteine protease and two proteins of unknown function. The candidate genes also included a gene encoding a Kazal-like protease inhibitor, although this protein lacked a secretion signal. For the *Pl. muralis* branch, 157 genes were found to be under selection (Supplementary table 13), including nine encoding secreted proteins (5.17% of the secreted proteins tested): an NADPH-hemoprotein reductase, a ubiquitin-specific peptidase, a protein with an Lsm domain (found in proteins involved in mRNA maturation), a glycosyl hydrolase, a protein with a heavy metal-associated domain, three proteins with EGF-like or filamin domains, and one with no identified domain. The only secreted proteins with a positive selection signal in the two branches were the cysteine protease (no secretion peptide detected in *Pl. muralis*) and one of the proteins of unknown function.

**Table 3.** Function of genes with a signal of positive selection in the branch-site tests.

|  | *Pl. viticola* branch | *Pl. muralis* branch | Both branches |
| --- | --- | --- | --- |
| Secreted proteins | 3 | 9 | 2 |
| Transmembrane transporters | 9 | 5 | 0 |
| RNA modifications & processing* | 9 | 11 | 2 |
| Other functions | 101 | 101 | 14 |
| Unknown function** | 9 | 31 | 0 |

\*including splicing factors, mRNA capping proteins, proteins with Lsm domain, RNA methyltransferases, chromodomain proteins, DEAD/DEAH box RNA helicases, RNA polymerase subunits and proteins involved in nonsense mediated mRNA decay.
\*\*proteins with no mapped GO term and not belonging to any other categories in this table.
The precise function of secreted proteins and transmembrane transporters are detailed in the main text. The functions of all proteins are available in Supplementary tables 12 and 13.

Several genes encoding transporters also presented significant positive selection signals, and some had functions potentially involved in plant-pathogen interactions: for *Pl. viticola*, a major facilitator superfamily transporter with 1,3-beta-glucan synthase activity, a multidrug transporter, a copper ion transporter and a folate-biopterin transporter, and for *Pl. muralis,* an amino-acid transporter and a phospholipid-translocating ATPase. These transporters may play an important role in interactions between *Pl. viticola* and *Pl. muralis* and their hosts, notably for the acquisition of specific nutrients. For example, a sucrose transporter plays a major role not only in carbon uptake, but also in the avoidance of plant defenses in the biotrophic corn smut fungus *Ustilago maydis* (Wahl et al. 2010). These transporters may also be involved in resistance to plant defense compounds or fungicides (Del Sorbo et al. 2000).

Enrichment in functional terms was found only in the GO enrichment analysis for the *Pl. viticola* branch. The terms for which enrichment was detected were “snoRNA binding” (*q*=0.014) and “nuclear localization binding” (*q*=0.074). Small nucleolar RNAs (snoRNAs) are involved principally in ribosomal RNA modifications, but also in RNA interference (Scott & Ono 2011), and two of the three proteins with the “nuclear localization binding” term were transporters of ribosomal proteins. In addition, both tested branches included multiple candidate genes with functions linked to RNA modification or processing (Table 3). Genes under selection in the *Pl. viticola* branch included two orthologs of *Ph. infestans* genes encoding chromodomain proteins potentially involved in gene silencing (PITG_07902 and PITG_15837, Vetukuri et al. 2011). The other genes were linked to RNA metabolism, mRNA maturation and processing, or rRNA and tRNA modification. The genes with a selection signal in both branches included a snoRNA binding protein and a DEAD/DEAH RNA helicase. Small RNAs play a major role in plant-pathogen interactions, being involved in the regulation of pathogen virulence by the host (host-induced gene silencing) and the manipulation of host immunity by the pathogen (reviewed in Weiberg & Jin 2015), potentially accounting for the positive selection signal for some components of the RNA machinery. A recent study by Brilli et al. (2018) demonstrated the presence of RNA interference machinery genes in *Pl. viticola* and the existence of a bidirectional exchange of small RNAs between *Pl. viticola* and its host *V. vinifera*, confirming the importance of small RNAs in this pathosystem.

Our list of candidate genes could guide the discovery of new effectors and the prioritization of functional validation analyses. This is particularly important for obligate biotrophic species, for which functional studies are difficult and currently restricted to heterologous expression systems. Genome editing is currently being developed in the hemi-biotrophic *Phytophthora* genus, the degree of success differing between species (Hoogen & Govers 2018), and may be even more challenging in biotrophic downy mildews.

## CONCLUSIONS

This study presents a genome assembly of *Pl. viticola*, the grapevine downy mildew pathogen, generated exclusively with using PacBio long reads at high coverage. This sequencing strategy yielded a better assembly of repetitive regions and, consequently, improved the detection of genes encoding secreted proteins potentially involved in pathogenicity and preferentially located in these regions. We also assembled the genome of *Pl. muralis* with short reads, the first step in the comparative genomics study of multiple *Plasmopara* species infecting hosts from the Vitaceae (Rouxel et al. 2014, 2013), which will shed light on the genetic mechanisms underlying host specialization in this species complex. The different approaches used highlighted the importance of cytoplasmic RxLR effectors for the adaptation of *Pl. viticola* and *Pl. muralis* to their host, as already shown for other pathogens in the order Peronosporales (Bozkurt et al. 2012). Our genome-wide approach detected positive selection signals for other candidate genes, including several genes encoding transporters and genes linked to the RNA machinery.

The availability of a high-quality reference genome for *Pl. viticola* also paves the way for robust population genetic studies based on genome-wide diversity (i.e. by resequencing individuals), such as population genome scans to detect genes under selection and genome-wide association studies to link genotype and phenotype variations. This work will improve our understanding of the genetic basis of the rapid adaptation of *Pl. viticola* to abiotic (e.g. fungicides) and biotic (e.g. partially resistant cultivars) factors.

## Supporting information

## ACKNOWLEDGMENTS

We thank an anonymous reviewer for their suggestions which improved this manuscript. We thank Céline Bénétreau, Jérôme Jolivet and Sylvie Richart-Cervera for their help with the production of *Pl. viticola* material, Erika Sallet and Ludovic Legrand for helping us with the gene and functional annotations and David Rengel for his assistance with the analysis of RNA-seq data. We also thank Sophien Kamoun for his comments. We thank the Genotoul bioinformatics platform Toulouse Midi-Pyrenees (Bioinfo Genotoul) for providing help and computing resources. This study was funded by the European Commission [INNOVINE, FP7-KBBE-311775] and the French National Research Agency [GANDALF project, ANR-12-ADAP-0009; EFFECTOORES project, ANR-13-ADAP-0003; Investments for the future program in the Cluster of Excellence COTE, ANR-10-LABX-45]. This work was performed in collaboration with the GeT core facility, Toulouse, France (http://get.genotoul.fr), and was supported by the France Génomique National infrastructure, funded as part of the Investments for the future program managed by the French National Research Agency [ANR-10-INBS-09] and by the GET-PACBIO program [“Programme opérationnel FEDER-FSE MIDI-PYRENEES ET GARONNE 2014-2020”].

