## Supplementary material for "A high-quality grapevine downy mildew genome assembly reveals rapidly evolving and lineage-specific putative host adaptation genes"

**This document includes:**

Supplementary note

Supplementary figure 1 Comparison between the long-read and short-read assemblies of the *Pl. viticola* INRA-PV221 isolate.

Supplementary figure 2 Synteny between *Pl. viticola* and *Pl. muralis*, *Pl. halstedii*, *Ph. infestans* and *Py. ultimum*.

Supplementary figure 3 Phylogenetic relationships between oomycete species.

Supplementary figure 4 Alignment of *Pl. viticola* and *Pl. muralis* proteins with a WY motif but no RxLR signal.

Supplementary figure 5 Local gene density and organization in *Pl. muralis.*

Supplementary table 1 Oomycete species and data origin.

Supplementary table 2 Assembly statistics for published *Pl. viticola* genome sequences.

Supplementary table 3 Type of repeat sequences in the *Pl. viticola* and *Pl. muralis* genomes.

Supplementary table 4 Statistics of the synteny analysis between the genomes of *Pl. viticola* and four other oomycete species.

Supplementary table 5 GO enrichment analysis for ortholog groups missing in oomycete biotrophic species.

Supplementary table 6 Number of flavoproteins in biotrophic and non-biotrophic oomycetes.

Supplementary table 7 KEGG annotation of *Ph. infestans* orthologs missing in biotrophic species.

Supplementary table 8 GO enrichment analysis for the secretome of *Pl. viticola.*

Supplementary table 9 GO enrichment analysis for the secretome of *Pl. muralis.*

Supplementary table 10 Results of the detection of candidate RxLR effector genes in *Pl. viticola* and *Pl. muralis.*

Supplementary table 11 *Pl. viticola* and *Pl. muralis* ortholog pairs with a dN/dS above 1.

Supplementary table 12 *Pl. viticola* genes with a significant signal in the branch-site test.

Supplementary table 13 *Pl. muralis* genes with a significant signal in the branch-site test.

### SUPPLEMENTARY NOTE

#### Genome assembly for *Pl. viticola*

The PBcR wgs8.3rc2 assembly pipeline (Berlin *et al.* 2015) was used to perform read correction and assemble the corrected reads with the following parameters:
i) only raw reads longer than 5 kb were considered
ii) 50x of corrected reads were collected (assembleCoverage=50)
iii) the unitigger bogart was tuned to handle heterozygosity (utgErrorRate=0.05 and utgMergeErrorRate=0.05).
Corrected reads were also assembled using FALCON (Chin *et al.* 2016) with the following parameters:
fc_ovlp_filter: --max_diff 240 --max_cov 150 --min_cov 30 --bestn 50 --min_len 5000
fc_ovlp_to_graph: --min_len 5000

As indicate in the main text of the paper, while the PBcR assembly was around twice the expected size of the genome due to high heterozygosity, it was more comprehensive than the FALCON assembly, so we removed redundancy in the PBcR assembly using an ad-hoc procedure: the contigs representing the alternative haplotype, defined as contigs fully included in larger contigs (with structural variations allowed), were removed. In a first step, an all vs all comparison was performed (megablast, min score=1000, minimum identity percentage=90), then syntenic HSPs were chained (custom script). A contig was removed when it was the shorter one of the pair and the chain spanned more than 80% of its length.
As some contigs were bacterial contaminants (mainly *Pseudomonas extremorientalis* and a *Stenotrophomonas*), FALCON was used to assemble bacterial genomes using parameters tuned for low coverage assembly (fc_ovlp_filter: --max_diff 50 --max_cov 50 --min_cov 1 --bestn 10 --min_len 5000, fc_ovlp_to_graph: --min_len 5000). The assembled bacterial genomes were used to remove contaminant-related contigs in the PBcR assembly (NCBI-TBLASTX, -e-value $10e^{-20}$, GC content > 56%).
Finally the PBcR contigs were scaffolded (19 gaps spanned) using a FALCON assembly (fc_ovlp_filter: --max_diff 10000 --max_cov 300 --min_cov 3 --bestn 1000 --min_len 5000).
Illumina paired-end reads (SRA accession numbers: SRX1970160, SRX1970161, SRX1970162) and 3 kb mate-pair reads (SRX1970164) used for a previous draft genome of the same isolate (Dussert *et al.* 2016) were mapped on the assembly sequence with BWA (Li & Durbin 2009). After running Pilon (parameters: --diploid --vcf --changes --fix bases) a first time, quality of the genome sequence measured with BUSCO v2 (Simão *et al.* 2015) with the Alveolata-Stramenopiles dataset was low (85% of expected genes). After observation of the assembly sequence and of the read mapping in a genome browser, we concluded that intra-individual variations for indel polymorphism were not properly phased (i.e. indels close to each other and belonging to the two different haplotypes were often in the consensus assembly sequence) and were causing frame shifts in coding sequences. This was not corrected by Pilon, since the software only considers one site at a time. Heterozygous nucleotide sites with an error in the assembly sequence (for example, sites

that were A/T in the mapping data but C or G in the assembly) were also not corrected. To resolve these issues, the VCF output of Pilon was parsed using custom scripts to correct heterozygous nucleotide sites with an error and find pairs of neighboring heterozygous indels (distance < 10 bp) and correct only one of them. This procedure was performed three times, and for the final correction round, proposed corrections were checked on a genome browser and accepted or not.

**Genome assembly for *Pl. muralis***

Since *Pl. muralis* sporangia and sporiangiophores were collected in non-sterile conditions outside the lab, contaminants were expected. All Illumina reads from *Pl. muralis* were first assembled with SPAdes 3.9.0 (Bankevich *et al.* 2012) (k-mer sizes: 21, 33, 55, 77). Contigs were then aligned against the NCBI nt database with BLASTN+, and reads were re-mapped on the contigs using BWA, keeping only properly paired reads. Reads mapping to contigs identified as contaminants were discarded, and a new assembly was carried out. This procedure was performed until no known contaminants were detected (three times). Since the large majority of contaminant contigs had a very low read coverage, we also discarded contigs with a coverage inferior to 20x and no blast hits with oomycete sequences. It is very likely that some sequences belonging to *Pl. muralis* were discarded with this approach, but it was deemed preferable to reduce contaminants to a minimum. Finally, we only kept contigs with a size superior to 1 kb. The median read coverage for the final assembly was 120x.

**Annotation**

**Gene annotation.** For *Pl. viticola*, after aligning RNA-seq reads with STAR 2.4 (Dobin *et al.* 2013) (--alignIntronMax 20000 --outFilterMismatchNmax 8, keeping only properly paired reads and discarding secondary alignments), genes were first predicted on the softmasked genome (de novo repeats and Repbase db) with BRAKER 1.8 (Hoff *et al.* 2016) (--alternatives-from-evidence=false; GeneMark-ES Suite 4.21, AUGUSTUS 3.1). Aligned RNA-seq reads were also assembled into transcripts using Trinity release 20140717 (Grabherr *et al.* 2011) (genome-guided assembly, --jaccard-clip), and the gene predictor SNAP (Korf 2004) was trained with the gene models from BRAKER. SNAP was then run on the softmasked genome in MAKER 2.31 (Holt & Yandell 2011), with the assembled transcripts and the proteomes of 9 other oomycetes (Supplementary table 1) as evidence. Observation of the results from BRAKER and SNAP in a genome browser indicated that gene models from BRAKER were of much higher quality. We chose to keep all BRAKER gene models, and only SNAP gene models not overlapping with BRAKER ones. A final MAKER run was then carried out to predict UTRs sequences, but this information was not used for any analyses, since observation of the annotation in a genome browser indicated that the quality of the predicted UTRs was low, with a high prevalence of too long sequences.

For *Pl. muralis*, for which no RNA data was available, we first made an initial MAKER run with AUGUSTUS trained with the *Pl. viticola* gene annotation as gene predictor and *Pl. viticola* proteins as evidence. AUGUSTUS and SNAP were then trained with this preliminary annotation. As previously, SNAP gene models were of lower quality, so we only kept SNAP models not overlapping AUGUSTUS models. Only scaffolds with a size greater than 2 kb were annotated.

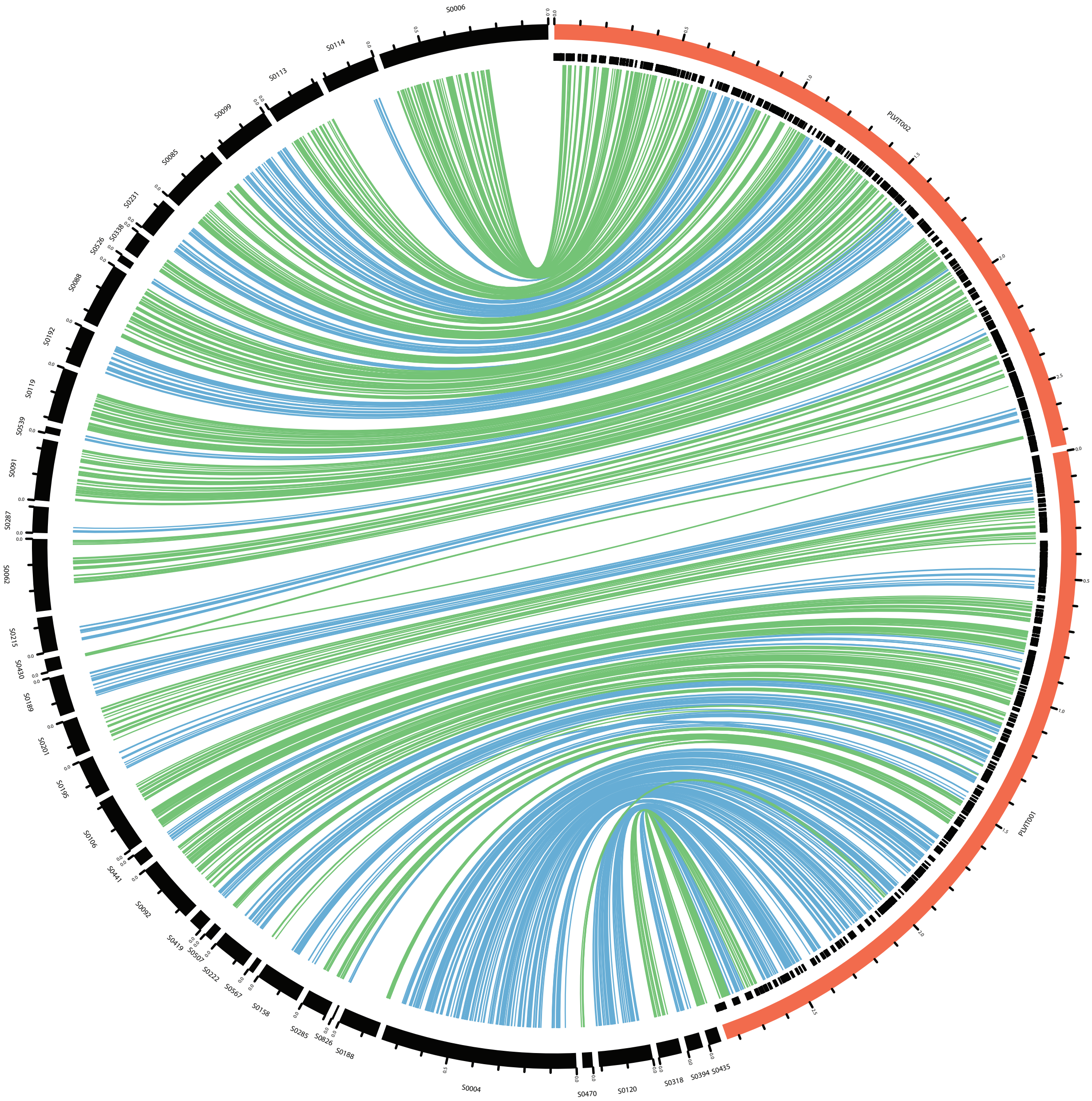

**Supplementary figure 1 Comparison between the long-read and short-read assemblies of the *Pl. viticola* INRA-PV221 isolate.** Only the results for the two biggest scaffolds of the long-read assembly are presented. Scaffolds of the long-read assembly are in red, those of the short-read assembly are in black, with tick marks each 100 kb. Repeat regions of the long-read assembly are represented as black boxes under the scaffolds. Links between scaffolds represents correspondance between genes. Links have different colors for alternating short-read scaffolds for clarity.

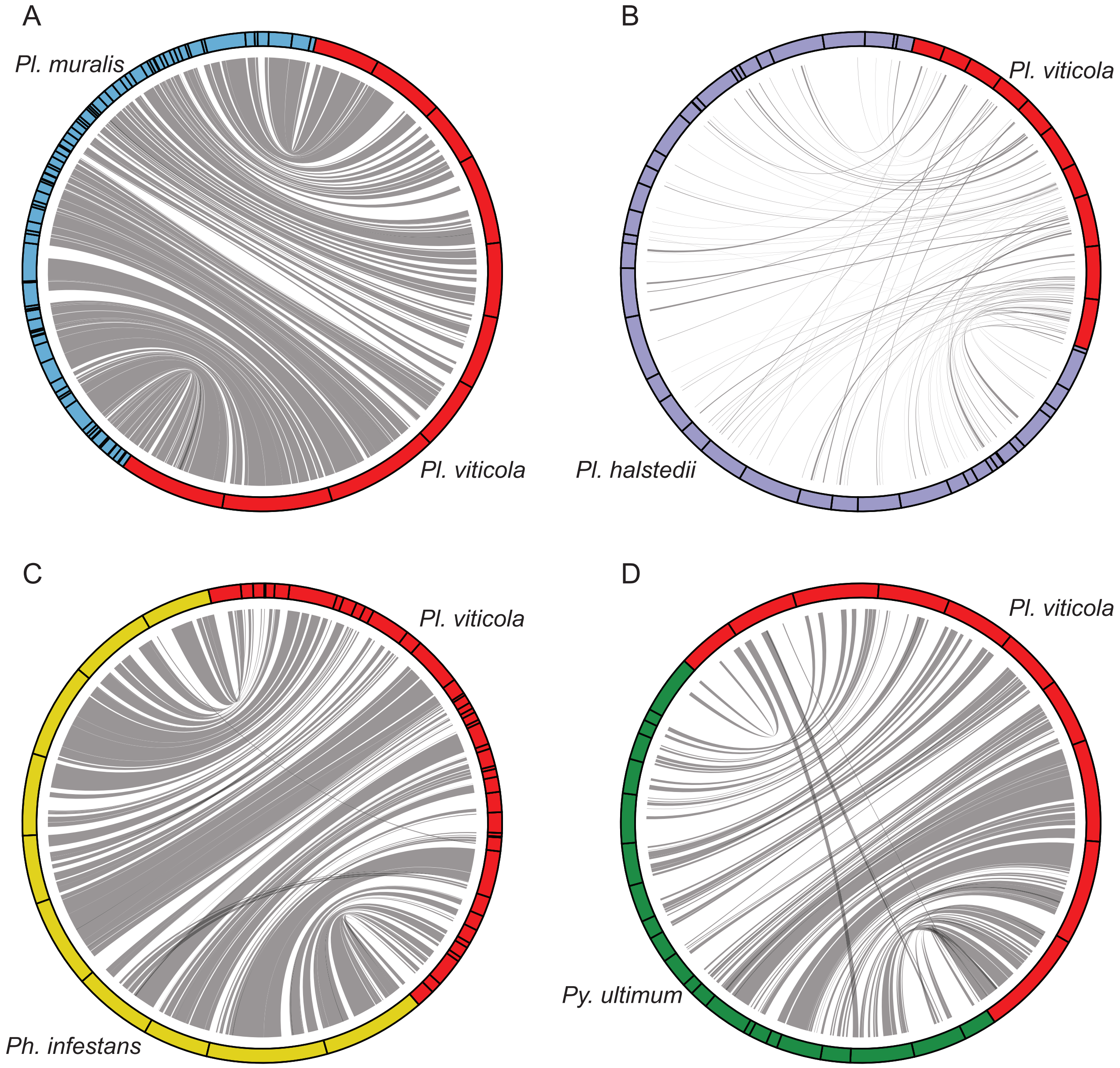

**Supplementary figure 2 Synteny between *Pl. viticola* and (A) *Pl. muralis*, (B) *Pl. halstedii*, (C) *Ph. infestans* and (D) *Py. ultimum*.** Syntenic regions are showed only for the ten largest *Pl. viticola* scaffolds, except for the comparison with *Ph. infestans*, where data is shown for the ten largest *Ph. infestans* scaffolds. Grey ribbons represent syntenic regions between genomes.

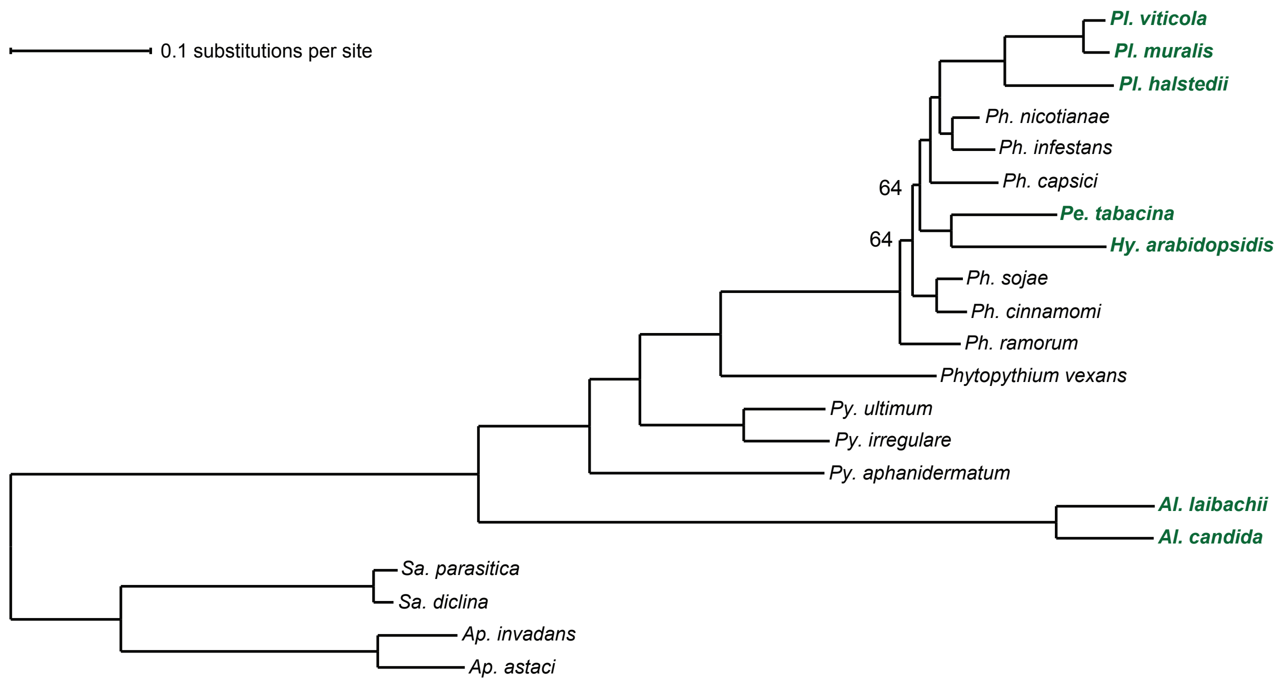

**Supplementary figure 3 Phylogenetic relationships between Oomycete species.** The maximum likelihood species tree was build using 144 single-copy orthologs. All nodes had a bootstrap support of 100% unless otherwise indicated. Biotrophic species names are shown in bold green typeface.

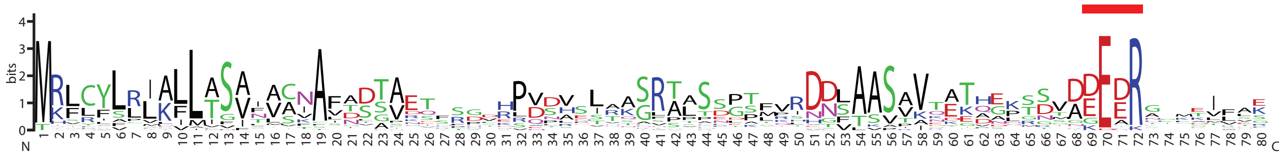

**Supplementary figure 4 Alignment of *Pl. viticola* and *Pl. muralis* proteins with a WY motif but no RXLR signal.** Proteins (38 sequences) were aligned with MAFFT and positions with more than 85% missing data (gaps) were discarded. The dEER motif common to all proteins is at positions 69-71 and is indicated by a red box.

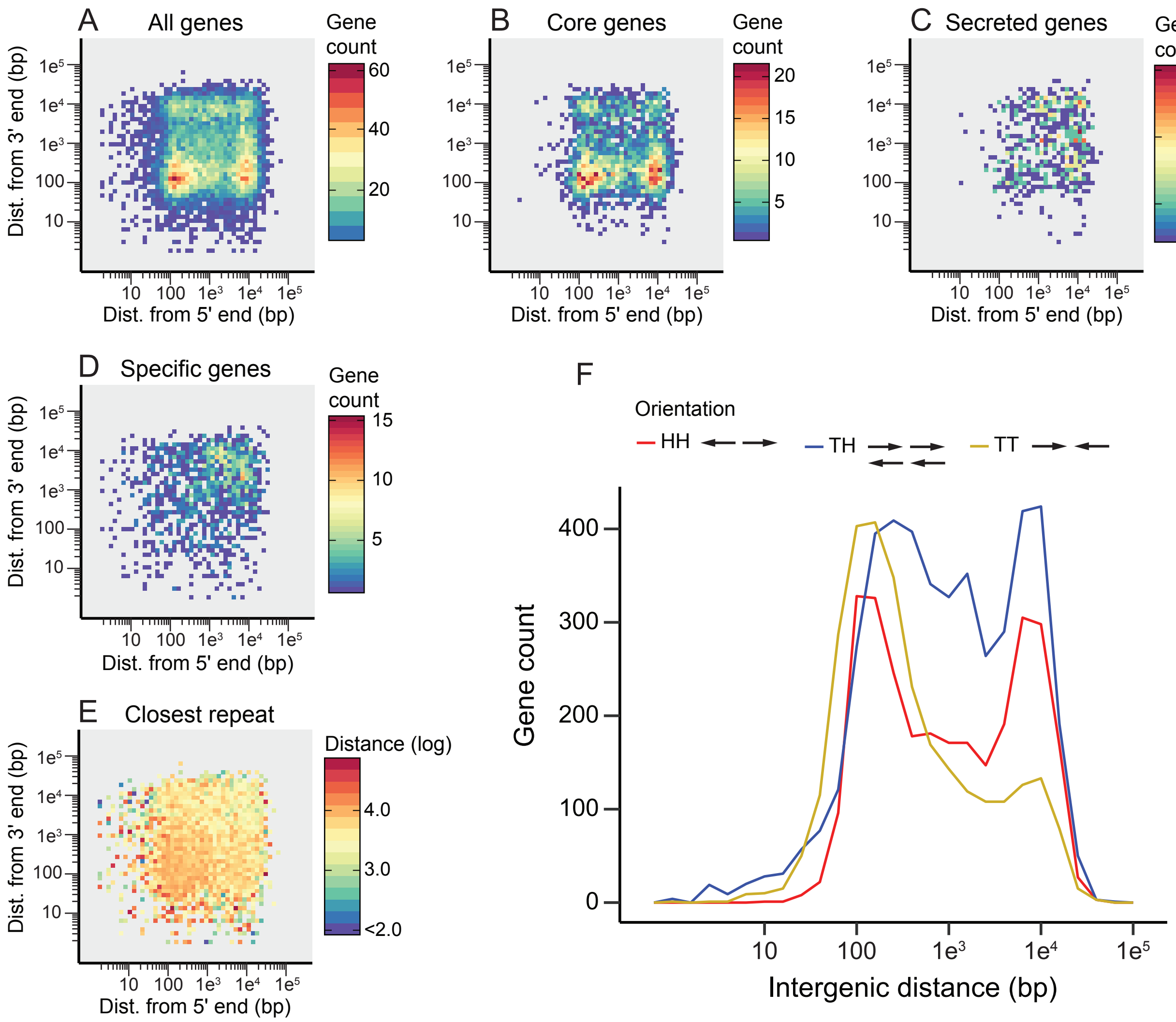

**Supplementary figure 5 Local gene density and organization in *Pl. muralis*.** Local gene density when considering (A) all genes, (B) core genes found in ortholog groups common to all Oomycetes, (C) genes encoding secreted proteins and (D) genes only found in *Pl. muralis* or in both *Pl. viticola* and *Pl. muralis*. The number of genes for two-dimensional (5' and 3') intergenic distance bins is represented as a heatmap. (E) Mean distance between genes and their closest repeat element, represented as a heatmap for two-dimensional intergenic distance bins. (F) Orientation and intergenic distance of adjacent gene pairs. The gene count for different intergenic distances has been plotted for adjacent gene pairs with a head-to-head (HH), tail-to-head (TH) and tail-to-tail (TT) orientation. Distances are on a log scale for all figures.

Supplementary table 1 Oomycete species and data origin

| Species | Code | Source | Assembly version | Reference | Used for | | | |
|---|---|---|---|---|---|---|---|---|
| | | | | | Annotation | Orthology | GO enrichment for biotrophy | Branch-site tests |
| *Albugo candida* | ALCA | Ensembl | ASM107853v1 | McMullan *et al.* 2015 | X | X | | |
| *Albugo laibachii* | ALLA | Ensembl | ENA1 | Kemen *et al.* 2011 | X | X | | |
| *Aphanomyces astaci* | APAS | Ensembl | Apha_asta_APO3_V1 | - | | X | X | |
| *Aphanomyces invadans* | APIN | Ensembl | Apha_inva_NJM9701_V1 | - | | X | X | |
| *Hyaloperonospora arabidopsidis* | HYAR | Ensembl | HyaAraEmoy2_2.0 | Baxter *et al.* 2010 | X | X | | |
| *Peronospora tabacina* | PETA | Corresponding author | 968-J2_v1 | Derevnina *et al.* 2015 | | X | | |
| *Phytophthora capsici* | PHCA | JGI | LT1534 v11.0 | Lamour *et al.* 2012 | X | X | | X |
| *Phytophthora cinnamomi* | PHCI | JGI | v1.0 | - | | X | | |
| *Phytophthora infestans* | PHIN | Ensembl | ASM14294v1 (T30-4) | Haas *et al.* 2009 | X | X | X | X |
| *Phytophthora nicotianae* | PHNI | Ensembl | ASM148298v1 | Liu *et al.* 2016 | | X | X | X |
| *Phytophthora ramorum* | PHRA | JGI | v1.1 | Tyler *et al.* 2006 | | X | X | |
| *Phytophthora sojae* | PHSO | Ensembl | P_sojae_V3_0 | Tyler *et al.* 2006 | | X | X | |
| *Plasmopara halstedii* | PLHA | Senckenberg Data & Metadata Repository* | - | Sharma *et al.* 2015 | X | X | | X |
| *Plasmopara muralis* | PLMU | | | This study | | X | | X |
| *Plasmopara viticola* | PLVI | | | This study | | X | | X |
| *Pythium aphanidermatum* | PYAP | Ensembl | pag1_scaffolds_v1 | Adhikari *et al.* 2013 | X | X | | |
| *Pythium irregulare* | PYIR | Ensembl | pir_scaffolds_v1 | Adhikari *et al.* 2013 | | X | | |
| *Pythium ultimum* | PYUL | Ensembl | pug | Lévesque *et al.* 2010 | X | X | X | |
| *Phytopythium vexans* | PYVE | Ensembl | pve_scaffolds_v1 | Adhikari *et al.* 2013 | | X | | |
| *Saprolegnia diclina* | SADI | Ensembl | Sap_diclina_VS20_V1 | - | | X | X | |
| *Saprolegnia parasitica* | SAPA | Ensembl | ASM15154v2 | Jiang *et al.* 2013 | X | X | X | |

Annotation: the proteomes of the species have been used as evidence for gene annotation. Orthology: proteomes used for the orthology analysis.
GO enrichment for biotrophy: the functional annotation of these species has been used for the GO enrichment analysis for ortholog groups absent in biotrophic species.
*http://dataportal-senckenberg.de/database/metacat/rsharma.26.4/bikf

Supplementary table 2 Assembly statistics for published *Pl. viticola* genome sequences

| | JL-7-2 (Yin *et al.* 2017) | PvitFEM01 (Brilli *et al.* 2018) | INRA-PV221 (Dussert *et al.* 2016) | INRA-PV221 (this study) |
|---|---|---|---|---|
| Assembly size (Mb) | 101.3 | 83.0 | 74.8 | 93.0 |
| Total gap length | 16.9 Mb | 3.1 Mb | 2.1 Mb | 34.3 kb |
| Number of scaffolds (minimum size) | 2,165 (500 pb) | 57,336 (300 bp) | 1,883 (1 kb) | 358 (20 kb) |
| Max. scaffold size (kb) | 806 | 132 | 763 | 2,850 |
| N50 (size in kb / count) | 172.3 / 172 | 4.1 / 3,733 | 180.6 / 130 | 706.5 / 38 |
| % of repeat elements | 25.60 | NA | NA | 37.34 |
| BUSCO genome [S / D / F] (%) | 66.6 [61.5 / 4.7 / 0.4] | 61.1 [58.5 / 0.0 / 2.6] | 92.8 [86.8 / 4.7 / 1.3] | 95.7 [93.6 / 1.7 / 0.4] |
| Number of genes | 17,014 | 38,298 | NA | 15,960 |
| BUSCO proteome [S / D / F] (%) | 65.8 [62.4 / 2.1 /1.3] | 56.4 [53.0 / 0.4 / 3.0] | NA | 97.4 [94.0 / 3.0 / 0.4] |

NA: data not available

BUSCO analyses have been carried out with the Alveolata-Stramenopiles data set. S: single copy, D: duplicated, F: fragmented genes

Supplementary table 3 Type of repeat sequences in the *Pl. viticola* and *Pl. muralis* genomes

| | *Pl. viticola* | *Pl. muralis* |
|---|---|---|
| *Class I* | | |
| DIRS-like | 1.69 | 4.36 |
| LINES | 6.62 | 10.17 |
| LTR | 42.93 | 61.61 |
| LTR (non-autonomous derivatives) | 35.53 | 0.77 |
| Penelope-like | 0.61 | 0.70 |
| Others | 0.00 | 4.90 |
| *Class II* | | |
| Crypton | 0.00 | 4.36 |
| Maverick | 6.74 | 3.94 |
| TIR | 3.34 | 6.77 |
| Others | 0.99 | 0.82 |
| *No category* | 1.55 | 1.60 |

Repeat sequences are classified following Wicker *et al.* (2007), and numbers are percentages of repeat of a given type.

Supplementary table 4 Statistics of the synteny analysis between the genomes of *Pl. viticola* and four other oomycete species

| | vs *Pl. muralis* | vs *Pl. halstedii* | vs *Ph. infestans* | vs *Py. ultimum* |
|---|---|---|---|---|
| Mean number of genes by syntenic block [range] | 23 [5 – 130] | 7 [5 – 21] | 26 [5 – 242] | 16 [5 – 141] |
| Mean syntenic block size in *Pl. viticola* [range] (kb) | 132 [5 – 716] | 38 [6 – 128] | 172 [5 – 1,533] | 130 [9 – 973] |
| Mean syntenic block size in compared species [range] (kb) | 118 [5 – 686] | 29 [6 – 153] | 414 [5 – 2,007] | 80 [3 – 618] |
| Total size of syntenic regions in *Pl. viticola* (Mb) | 46.49 | 9.46 | 40.43 | 35.06 |
| Total size of syntenic regions in compared species (Mb) | 41.49 | 7.40 | 97.33 | 21.73 |
| Total number of genes in syntenic blocks | 7,900 | 1,184 | 6,136 | 4,379 |

Supplementary table 5 GO enrichment analysis for ortholog groups missing in oomycete biotrophic species

| GO ID | GO term | Ontology | Annotated | Significant | Expected | *P* -values | Empirical *q* -values |
|---|---|---|---|---|---|---|---|
| GO:0006310 | DNA recombination | BP | 749 | 117 | 11.71 | 1.02E-81 | 0.00 |
| GO:0015074 | DNA integration | BP | 778 | 118 | 12.16 | 6.26E-81 | 0.00 |
| GO:0032217 | riboflavin transporter activity | MF | 28 | 28 | 0.39 | 7.56E-53 | 0.00 |
| GO:0015171 | amino acid transmembrane transporter activity | MF | 403 | 54 | 5.63 | 7.53E-36 | 0.00 |
| GO:0031419 | cobalamin binding | MF | 22 | 19 | 0.31 | 6.96E-33 | 0.00 |
| GO:0030798 | trans-aconitate 2-methyltransferase activity | MF | 18 | 16 | 0.25 | 2.72E-28 | 0.00 |
| GO:0005216 | ion channel activity | MF | 1057 | 63 | 14.76 | 9.95E-22 | 0.00 |
| GO:0003884 | D-amino-acid oxidase activity | MF | 24 | 14 | 0.34 | 1.67E-20 | 0.00 |
| GO:0046416 | D-amino acid metabolic process | BP | 24 | 14 | 0.38 | 7.45E-20 | 0.00 |
| GO:0036361 | racemase activity, acting on amino acids and derivatives | MF | 17 | 12 | 0.24 | 2.96E-19 | 0.00 |
| GO:0003677 | DNA binding | MF | 4254 | 135 | 59.38 | 4.71E-19 | 0.00 |
| GO:0006508 | proteolysis | BP | 1381 | 70 | 21.58 | 4.95E-18 | 0.00 |
| GO:0004367 | glycerol-3-phosphate dehydrogenase [NAD+] activity | MF | 27 | 13 | 0.38 | 1.18E-17 | 0.00 |
| GO:0004494 | methylmalonyl-CoA mutase activity | MF | 9 | 9 | 0.13 | 1.94E-17 | 0.00 |
| GO:0005089 | Rho guanyl-nucleotide exchange factor activity | MF | 28 | 13 | 0.39 | 2.17E-17 | 0.00 |
| GO:0046168 | glycerol-3-phosphate catabolic process | BP | 26 | 13 | 0.41 | 2.48E-17 | 0.00 |
| GO:0035023 | regulation of Rho protein signal transduction | BP | 28 | 13 | 0.44 | 8.68E-17 | 0.00 |
| GO:0008705 | methionine synthase activity | MF | 11 | 9 | 0.15 | 1.04E-15 | 0.00 |
| GO:0004114 | 3',5'-cyclic-nucleotide phosphodiesterase activity | MF | 127 | 20 | 1.77 | 1.60E-15 | 0.00 |
| GO:0003865 | 3-oxo-5-alpha-steroid 4-dehydrogenase activity | MF | 9 | 8 | 0.13 | 1.24E-14 | 0.00 |
| GO:0031151 | histone methyltransferase activity (H3-K79 specific) | MF | 130 | 19 | 1.81 | 3.33E-14 | 0.00 |
| GO:0043531 | ADP binding | MF | 11 | 8 | 0.15 | 2.22E-13 | 0.00 |
| GO:0000104 | succinate dehydrogenase activity | MF | 43 | 12 | 0.6 | 5.26E-13 | 0.00 |
| GO:0019236 | response to pheromone | BP | 22 | 9 | 0.34 | 2.16E-11 | 0.00 |
| GO:0005509 | calcium ion binding | MF | 2173 | 72 | 30.33 | 2.29E-11 | 0.00 |
| GO:0009235 | cobalamin metabolic process | BP | 29 | 9 | 0.45 | 3.95E-10 | 0.00 |
| GO:0016628 | oxidoreductase activity, acting on the CH-CH group of donors, NAD or NADP as acceptor | MF | 53 | 10 | 0.74 | 3.04E-09 | 0.00 |
| GO:0016559 | peroxisome fission | BP | 40 | 9 | 0.63 | 9.25E-09 | 0.00 |
| GO:0008202 | steroid metabolic process | BP | 58 | 10 | 0.91 | 2.13E-08 | 0.00 |
| GO:0007154 | cell communication | BP | 2015 | 76 | 31.49 | 2.21E-08 | 0.00 |
| GO:0019888 | protein phosphatase regulator activity | MF | 83 | 11 | 1.16 | 2.39E-08 | 0.00 |
| GO:0071949 | FAD binding | MF | 154 | 14 | 2.15 | 4.20E-08 | 0.00 |
| GO:0051920 | peroxiredoxin activity | MF | 63 | 9 | 0.88 | 2.34E-07 | 0.00 |
| GO:0004842 | ubiquitin-protein transferase activity | MF | 794 | 29 | 11.08 | 3.70E-06 | 0.00 |
| GO:0007186 | G-protein coupled receptor signaling pathway | BP | 93 | 9 | 1.45 | 1.56E-05 | 2.77E-04 |
| GO:0005388 | calcium-transporting ATPase activity | MF | 82 | 8 | 1.14 | 2.01E-05 | 2.77E-04 |
| GO:0042558 | pteridine-containing compound metabolic process | BP | 127 | 10 | 1.98 | 3.27E-05 | 6.74E-04 |
| GO:0016849 | phosphorus-oxygen lyase activity | MF | 154 | 10 | 2.15 | 6.75E-05 | 1.71E-03 |
| GO:0051726 | regulation of cell cycle | BP | 447 | 19 | 6.99 | 9.46E-05 | 2.05E-03 |
| GO:0009190 | cyclic nucleotide biosynthetic process | BP | 155 | 10 | 2.42 | 1.74E-04 | 2.75E-03 |
| GO:0009116 | nucleoside metabolic process | BP | 372 | 16 | 5.81 | 2.90E-04 | 4.63E-03 |
| GO:0016042 | lipid catabolic process | BP | 245 | 12 | 3.83 | 5.24E-04 | 8.08E-03 |
| GO:0015297 | antiporter activity | MF | 454 | 16 | 6.34 | 7.73E-04 | 1.10E-02 |
| GO:0015103 | inorganic anion transmembrane transporter activity | MF | 337 | 13 | 4.7 | 1.04E-03 | 1.37E-02 |
| GO:0007165 | signal transduction | BP | 1968 | 67 | 30.76 | 1.09E-03 | 1.41E-02 |
| GO:0008138 | protein tyrosine/serine/threonine phosphatase activity | MF | 229 | 10 | 3.2 | 1.56E-03 | 1.82E-02 |
| GO:0003777 | microtubule motor activity | MF | 926 | 25 | 12.93 | 1.58E-03 | 1.82E-02 |
| GO:0004420 | hydroxymethylglutaryl-CoA reductase (NADPH) activity | MF | 17 | 3 | 0.24 | 1.59E-03 | 1.82E-02 |
| GO:0010181 | FMN binding | MF | 363 | 13 | 5.07 | 2.02E-03 | 2.60E-02 |
| GO:0016504 | peptidase activator activity | MF | 6 | 2 | 0.08 | 2.81E-03 | 3.54E-02 |
| GO:0050982 | detection of mechanical stimulus | BP | 7 | 2 | 0.11 | 4.86E-03 | 6.64E-02 |
| GO:0051287 | NAD binding | MF | 410 | 13 | 5.72 | 5.59E-03 | 8.00E-02 |

Ontology: BP, biological process; MF, molecular function. Annotated: total number of proteins mapped with the GO term. Significant: number of proteins absent from biotrophic species mapped with the GO term. Expected: expected number of proteins mapped with the GO term if randomly distributed. *P* -values: computed using the Fisher exact test. Empirical *q* -values: computed after 200 permutations.

Supplementary table 6 Number of flavoproteins in biotrophic and non-biotrophic oomycetes

| Species | FAD binding | FMN binding | Total flavoproteins |
|---|---|---|---|
| Non-biotrophs | | | |
| *Aphanomyces astaci* | 23 | 25 | 48 |
| *Aphanomyces invadans* | 17 | 25 | 42 |
| *Phytophthora infestans* | 23 | 43 | 66 |
| *Phytophthora nicotianae* | 21 | 44 | 65 |
| *Phytophthora ramorum* | 17 | 47 | 64 |
| *Phytophthora sojae* | 16 | 90 | 106 |
| *Pythium ultimum* | 12 | 46 | 58 |
| *Saprolegnia diclina* | 11 | 20 | 31 |
| *Saprolegnia parasitica* | 14 | 23 | 37 |
| Mean | 17.11 | 40.33 | 57.44 |
| Biotrophs | | | |
| *Albugo candida* | 3 | 8 | 11 |
| *Albugo laibachii* | 3 | 10 | 13 |
| *Hyaloperonospora arabidopsidis* | 6 | 15 | 21 |
| *Plasmopara halstedii* | 5 | 11 | 16 |
| *Plasmopara muralis* | 3 | 16 | 19 |
| *Plasmopara viticola* | 4 | 17 | 21 |
| Mean | 4.00 | 12.83 | 16.83 |

Supplementary table 7 KEGG annotation of *Ph. infestans* orthologs missing in biotrophic species

| Gene | Definition | Pathways |
|---|---|---|
| PITG_00496 | 3',5'-cyclic-nucleotide phosphodiesterase | Purine metabolism |
| PITG_01804 | 5-methyltetrahydrofolate--homocysteine methyltransferase (methionine synthase) | Cysteine and methionine metabolism; Selenocompound metabolism; One carbon pool by folate; Metabolic pathways; Biosynthesis of secondary metabolites; Biosynthesis of amino acids |
| PITG_01808 | methylmalonyl-CoA mutase | Valine, leucine and isoleucine degradation; Glyoxylate and dicarboxylate metabolism; Propanoate metabolism; Metabolic pathways; Carbon metabolism |
| PITG_02105 | uracil phosphoribosyltransferase | Pyrimidine metabolism; Metabolic pathways |
| PITG_05696 | glycerol-3-phosphate dehydrogenase (NAD+) | Glycerophospholipid metabolism; Biosynthesis of secondary metabolites |
| PITG_05701 | glycerol-3-phosphate dehydrogenase (NAD+) | Glycerophospholipid metabolism; Biosynthesis of secondary metabolites |
| PITG_06147 | nicotinamide mononucleotide adenylyltransferase | Nicotinate and nicotinamide metabolism; Metabolic pathways |
| PITG_08135 | D-aspartate oxidase | Alanine, aspartate and glutamate metabolism; Peroxisome |
| PITG_10312 | patatin-like phospholipase domain-containing protein 2 | Glycerolipid metabolism; Metabolic pathways |
| PITG_13128 | 7-dehydrocholesterol reductase | Steroid biosynthesis; Metabolic pathways; Biosynthesis of secondary metabolites |
| PITG_18374 | lysophosphatidylcholine acyltransferase / lyso-PAF acetyltransferase | Glycerophospholipid metabolism; Ether lipid metabolism; Metabolic pathways |
| PITG_18748 | proteasome activator subunit 4 | Proteasome |
| PITG_18972 | sarcosine oxidase / L-pipecolate oxidase | Glycine, serine and threonine metabolism; Lysine degradation; Metabolic pathways; Peroxisome |
| PITG_21269 | proteasome activator subunit 4 | Proteasome |
| PITG_21426 | Delta7-sterol 5-desaturase | Steroid biosynthesis; Metabolic pathways; Biosynthesis of secondary metabolites; Biosynthesis of antibiotics |

Supplementary table 8 GO enrichment analysis for the secretome of *Pl. viticola*

| GO ID | GO term | Ontology | Annotated | Significant | Expected | *P* -values | Empirical *q* -values |
|---|---|---|---|---|---|---|---|
| GO:0004252 | serine-type endopeptidase activity | MF | 130 | 70 | 7.48 | 6.10E-53 | 0.00 |
| GO:0000272 | polysaccharide catabolic process | BP | 101 | 69 | 5.89 | 6.78E-33 | 0.00 |
| GO:0004553 | hydrolase activity, hydrolyzing O-glycosyl compounds | MF | 176 | 99 | 10.13 | 1.19E-32 | 0.00 |
| GO:0008810 | cellulase activity | MF | 41 | 30 | 2.36 | 5.17E-29 | 0.00 |
| GO:0006508 | proteolysis | BP | 723 | 125 | 42.18 | 4.66E-27 | 0.00 |
| GO:0030245 | cellulose catabolic process | BP | 28 | 23 | 1.63 | 1.80E-24 | 0.00 |
| GO:0005975 | carbohydrate metabolic process | BP | 358 | 126 | 20.89 | 1.84E-20 | 0.00 |
| GO:0030570 | pectate lyase activity | MF | 25 | 19 | 1.44 | 2.63E-19 | 0.00 |
| GO:0030246 | carbohydrate binding | MF | 92 | 44 | 5.30 | 1.01E-18 | 0.00 |
| GO:0030248 | cellulose binding | MF | 17 | 14 | 0.98 | 2.17E-15 | 0.00 |
| GO:0030599 | pectinesterase activity | MF | 21 | 15 | 1.21 | 8.27E-15 | 0.00 |
| GO:0042545 | cell wall modification | BP | 21 | 15 | 1.23 | 9.65E-15 | 0.00 |
| GO:0004568 | chitinase activity | MF | 16 | 12 | 0.92 | 1.74E-12 | 0.00 |
| GO:0006032 | chitin catabolic process | BP | 16 | 12 | 0.93 | 1.98E-12 | 0.00 |
| GO:0009405 | pathogenesis | BP | 38 | 15 | 2.22 | 1.09E-09 | 0.00 |
| GO:0016798 | hydrolase activity, acting on glycosyl bonds | MF | 199 | 110 | 11.46 | 2.20E-09 | 0.00 |
| GO:0006952 | defense response | BP | 53 | 17 | 3.09 | 3.56E-09 | 0.00 |
| GO:0016998 | cell wall macromolecule catabolic process | BP | 13 | 9 | 0.76 | 4.20E-09 | 0.00 |
| GO:0009251 | glucan catabolic process | BP | 47 | 32 | 2.74 | 2.59E-07 | 0.00 |
| GO:0004867 | serine-type endopeptidase inhibitor activity | MF | 8 | 6 | 0.46 | 8.99E-07 | 0.00 |
| GO:1900004 | negative regulation of serine-type endopeptidase activity | BP | 8 | 6 | 0.47 | 9.66E-07 | 0.00 |
| GO:0008422 | beta-glucosidase activity | MF | 23 | 9 | 1.32 | 2.57E-06 | 2.25E-04 |
| GO:0003993 | acid phosphatase activity | MF | 25 | 9 | 1.44 | 5.79E-06 | 4.13E-04 |
| GO:0008236 | serine-type peptidase activity | MF | 161 | 79 | 9.27 | 1.56E-05 | 4.13E-04 |
| GO:0004650 | polygalacturonase activity | MF | 5 | 4 | 0.29 | 5.19E-05 | 5.94E-04 |
| GO:0016755 | transferase activity, transferring amino-acyl groups | MF | 20 | 7 | 1.15 | 8.10E-05 | 7.62E-04 |
| GO:0071555 | cell wall organization | BP | 27 | 19 | 1.58 | 1.38E-04 | 1.83E-03 |
| GO:0008378 | galactosyltransferase activity | MF | 6 | 4 | 0.35 | 1.49E-04 | 1.95E-03 |
| GO:0004869 | cysteine-type endopeptidase inhibitor activity | MF | 7 | 4 | 0.40 | 3.31E-04 | 3.87E-03 |
| GO:0010951 | negative regulation of endopeptidase activity | BP | 15 | 10 | 0.88 | 3.31E-04 | 3.87E-03 |
| GO:0016722 | oxidoreductase activity, oxidizing metal ions | MF | 12 | 5 | 0.69 | 3.51E-04 | 3.87E-03 |
| GO:0003756 | protein disulfide isomerase activity | MF | 12 | 5 | 0.69 | 3.51E-04 | 3.87E-03 |
| GO:0010383 | cell wall polysaccharide metabolic process | BP | 9 | 6 | 0.53 | 7.41E-04 | 6.75E-03 |
| GO:0004601 | peroxidase activity | MF | 28 | 7 | 1.61 | 8.27E-04 | 7.43E-03 |
| GO:0008233 | peptidase activity | MF | 551 | 119 | 31.72 | 9.01E-04 | 7.92E-03 |
| GO:0045330 | aspartyl esterase activity | MF | 9 | 4 | 0.52 | 1.09E-03 | 1.00E-02 |
| GO:0045490 | pectin catabolic process | BP | 9 | 4 | 0.53 | 1.14E-03 | 1.00E-02 |
| GO:0034976 | response to endoplasmic reticulum stress | BP | 31 | 7 | 1.81 | 1.70E-03 | 2.02E-02 |
| GO:0008061 | chitin binding | MF | 5 | 3 | 0.29 | 1.74E-03 | 2.02E-02 |
| GO:0042124 | 1,3-beta-glucanosyltransferase activity | MF | 5 | 3 | 0.29 | 1.74E-03 | 2.02E-02 |
| GO:0034411 | cell wall (1->3)-beta-D-glucan biosynthetic process | BP | 5 | 3 | 0.29 | 1.81E-03 | 2.05E-02 |
| GO:0004197 | cysteine-type endopeptidase activity | MF | 32 | 7 | 1.84 | 1.92E-03 | 2.10E-02 |
| GO:0006493 | protein O-linked glycosylation | BP | 18 | 5 | 1.05 | 3.00E-03 | 3.16E-02 |
| GO:0005507 | copper ion binding | MF | 20 | 5 | 1.15 | 4.68E-03 | 4.85E-02 |
| GO:0072593 | reactive oxygen species metabolic process | BP | 13 | 4 | 0.76 | 5.36E-03 | 5.81E-02 |
| GO:0098869 | cellular oxidant detoxification | BP | 38 | 7 | 2.22 | 5.72E-03 | 6.07E-02 |
| GO:0044275 | cellular carbohydrate catabolic process | BP | 50 | 28 | 2.92 | 6.19E-03 | 6.29E-02 |
| GO:0003824 | catalytic activity | MF | 5353 | 446 | 308.20 | 8.79E-03 | 9.54E-02 |
| GO:0004181 | metallocarboxypeptidase activity | MF | 15 | 4 | 0.86 | 8.92E-03 | 9.54E-02 |

Ontology: BP, biological process; MF, molecular function. Annotated: total number of proteins mapped with the GO term. Significant: number of secreted proteins mapped with the GO term. Expected: expected number of proteins mapped with the GO term if randomly distributed. *P* -values: computed using the Fisher exact test. Empirical *q* -values: computed after 200 permutations.

Supplementary table 9 GO enrichment analysis for the secretome of *Pl. muralis*

| GO ID | GO term | Ontology | Annotated | Significant | Expected | *P* -values | Empirical *q* -values |
|---|---|---|---|---|---|---|---|
| GO:0004553 | hydrolase activity, hydrolyzing O-glycosyl compounds | MF | 96 | 38 | 3.97 | 2.19E-22 | 0.00 |
| GO:0005975 | carbohydrate metabolic process | BP | 223 | 62 | 10.78 | 7.09E-18 | 0.00 |
| GO:0004252 | serine-type endopeptidase activity | MF | 74 | 24 | 3.06 | 5.75E-16 | 0.00 |
| GO:0009405 | pathogenesis | BP | 21 | 14 | 1.01 | 2.23E-14 | 0.00 |
| GO:0006952 | defense response | BP | 27 | 15 | 1.3 | 1.22E-13 | 0.00 |
| GO:0006508 | proteolysis | BP | 437 | 60 | 21.12 | 1.74E-12 | 0.00 |
| GO:0030245 | cellulose catabolic process | BP | 16 | 9 | 0.77 | 1.05E-08 | 0.00 |
| GO:0030248 | cellulose binding | MF | 9 | 6 | 0.37 | 3.55E-07 | 0.00 |
| GO:0004866 | endopeptidase inhibitor activity | MF | 7 | 5 | 0.29 | 2.27E-06 | 0.00 |
| GO:0030246 | carbohydrate binding | MF | 42 | 15 | 1.74 | 4.07E-06 | 0.00 |
| GO:0010951 | negative regulation of endopeptidase activity | BP | 7 | 5 | 0.34 | 4.90E-06 | 0.00 |
| GO:0016722 | oxidoreductase activity, oxidizing metal ions | MF | 9 | 5 | 0.37 | 1.27E-05 | 0.00 |
| GO:0004197 | cysteine-type endopeptidase activity | MF | 22 | 7 | 0.91 | 1.89E-05 | 0.00 |
| GO:0016755 | transferase activity, transferring amino-acyl groups | MF | 22 | 7 | 0.91 | 1.89E-05 | 0.00 |
| GO:0000272 | polysaccharide catabolic process | BP | 46 | 20 | 2.22 | 3.03E-05 | 0.00 |
| GO:0008233 | peptidase activity | MF | 336 | 55 | 13.88 | 3.14E-05 | 0.00 |
| GO:0008422 | beta-glucosidase activity | MF | 17 | 6 | 0.7 | 3.95E-05 | 2.90E-04 |
| GO:0030599 | pectinesterase activity | MF | 14 | 5 | 0.58 | 1.70E-04 | 1.56E-03 |
| GO:0042124 | 1,3-beta-glucanosyltransferase activity | MF | 8 | 4 | 0.33 | 1.75E-04 | 1.56E-03 |
| GO:0003993 | acid phosphatase activity | MF | 23 | 6 | 0.95 | 2.61E-04 | 3.00E-03 |
| GO:0034411 | cell wall (1->3)-beta-D-glucan biosynthetic process | BP | 8 | 4 | 0.39 | 3.19E-04 | 3.00E-03 |
| GO:0005507 | copper ion binding | MF | 16 | 5 | 0.66 | 3.47E-04 | 3.00E-03 |
| GO:0042545 | cell wall modification | BP | 14 | 5 | 0.68 | 3.53E-04 | 3.00E-03 |
| GO:0009251 | glucan catabolic process | BP | 31 | 14 | 1.5 | 4.31E-04 | 3.70E-03 |
| GO:0044036 | cell wall macromolecule metabolic process | BP | 17 | 8 | 0.82 | 5.21E-04 | 4.73E-03 |
| GO:0004601 | peroxidase activity | MF | 18 | 5 | 0.74 | 6.36E-04 | 5.99E-03 |
| GO:0030570 | pectate lyase activity | MF | 5 | 3 | 0.21 | 6.55E-04 | 5.99E-03 |
| GO:0016263 | glycoprotein-N-acetylgalactosamine 3-beta-galactosyltransferase activity | MF | 5 | 3 | 0.21 | 6.55E-04 | 5.99E-03 |
| GO:0016267 | O-glycan processing, core 1 | BP | 5 | 3 | 0.24 | 1.04E-03 | 1.04E-02 |
| GO:0004568 | chitinase activity | MF | 6 | 3 | 0.25 | 1.27E-03 | 1.28E-02 |
| GO:0006955 | immune response | BP | 6 | 3 | 0.29 | 2.00E-03 | 2.05E-02 |
| GO:0006032 | chitin catabolic process | BP | 6 | 3 | 0.29 | 2.00E-03 | 2.05E-02 |
| GO:0003756 | protein disulfide isomerase activity | MF | 15 | 4 | 0.62 | 2.71E-03 | 2.94E-02 |
| GO:0010181 | FMN binding | MF | 16 | 4 | 0.66 | 3.49E-03 | 3.68E-02 |
| GO:0034976 | response to endoplasmic reticulum stress | BP | 33 | 6 | 1.59 | 4.40E-03 | 4.54E-02 |
| GO:0070011 | peptidase activity, acting on L-amino acid peptides | MF | 289 | 46 | 11.94 | 4.78E-03 | 4.96E-02 |
| GO:0098869 | cellular oxidant detoxification | BP | 24 | 5 | 1.16 | 5.04E-03 | 5.08E-02 |
| GO:0003824 | catalytic activity | MF | 3342 | 209 | 138.06 | 5.86E-03 | 5.68E-02 |
| GO:0045330 | aspartyl esterase activity | MF | 10 | 3 | 0.41 | 6.74E-03 | 6.56E-02 |
| GO:0072593 | reactive oxygen species metabolic process | BP | 9 | 3 | 0.43 | 7.53E-03 | 7.49E-02 |
| GO:0016798 | hydrolase activity, acting on glycosyl bonds | MF | 108 | 41 | 4.46 | 7.74E-03 | 7.49E-02 |
| GO:0010393 | galacturonan metabolic process | BP | 10 | 3 | 0.48 | 1.04E-02 | 9.40E-02 |
| GO:0045490 | pectin catabolic process | BP | 10 | 3 | 0.48 | 1.04E-02 | 9.40E-02 |
| GO:0045488 | pectin metabolic process | BP | 10 | 3 | 0.48 | 1.04E-02 | 9.40E-02 |

Ontology: BP, biological process; MF, molecular function. Annotated: total number of proteins mapped with the GO term. Significant: number of secreted proteins mapped with the GO term. Expected: expected number of proteins mapped with the GO term if randomly distributed. *P* -values: computed using the Fisher exact test. Empirical *q* -values: computed after 200 permutations.

Supplementary table 10 Results of the detection of candidate RxLR effector genes in *Pl. viticola* and *Pl. muralis*

| | *Pl. viticola* | *Pl. muralis* |
|---|---|---|
| Regex only | 58 (3) | 21 (0) |
| BLASTP only | 357 (312) | 80 (62) |
| Regex + BLASTP | 52 (25) | 15 (5) |
| Iterative BLASTP | 73 (1) | 16 (0) |
| Total | 540 (341) | 132 (67) |

Regex: search for regular expression using the Galaxy workflow of Cock *et al.* (2013).

BLASTP: BLASTP search at low stringency using an oomycete RxLR database.

Iterative BLASTP: BLASTP search at high stringency using the candidates detected by the Regex method as query.

Number of candidate effector genes showing structural homology to known RxLR effectors as detected by Phyre2 are shown in brackets.

Supplementary table 11 *Pl. viticola* and *Pl. muralis* ortholog pairs with a dN/dS above 1

| | | | | | Secreted | | RxLR | | GO terms | | InterPro domains | |
|---|---|---|---|---|---|---|---|---|---|---|---|---|
| PVIT_gene | PMUR_gene | dN/dS | dN | dS | *Pl. viticola* | *Pl. muralis* | *Pl. viticola* | *Pl. muralis* | *Pl. viticola* | *Pl. muralis* | *Pl. viticola* | *Pl. muralis* |
| PVIT_0000327 | PMUR_0010585 | 2.68 | 0.16 | 0.06 | no | no | no | no | | | | |
| **PVIT_0016683** | PMUR_0012182 | 2.27 | 0.08 | 0.04 | no | no | no | no | protein kinase activity; ATP binding; protein phosphorylation | protein kinase activity; ATP binding; protein phosphorylation | | |
| **PVIT_0003422** | PMUR_0003338 | 2.07 | 0.28 | 0.14 | no | no | no | no | | | | |
| **PVIT_0017530** | PMUR_0009546 | 2.04 | 0.37 | 0.18 | yes | no | yes | no | | | | |
| **PVIT_0012238** | PMUR_0003038 | 2.01 | 0.15 | 0.07 | yes | no | yes | no | | | | |
| PVIT_0017143 | PMUR_0012479 | 2.00 | 0.17 | 0.08 | no | no | no | no | | | | |
| PVIT_0014100 | PMUR_0004666 | 1.96 | 0.22 | 0.11 | no | no | no | no | | | | |
| **PVIT_0002472** | PMUR_0012195 | 1.92 | 0.18 | 0.09 | yes | yes | no | no | carbohydrate binding | | Jacalin-like lectin domain | Jacalin-like lectin domain |
| PVIT_0006091 | PMUR_0001598 | 1.92 | 0.22 | 0.12 | no | no | no | no | protein kinase activity; ATP binding; protein phosphorylation; kinase activity; phosphorylation | | | |
| **PVIT_0017049** | PMUR_0006507 | 1.91 | 0.14 | 0.07 | no | no | no | no | | | | |
| PVIT_0013491 | PMUR_0010387 | 1.84 | 0.30 | 0.16 | no | no | no | no | | | | |
| PVIT_0005573 | PMUR_0000810 | 1.76 | 0.21 | 0.12 | no | no | no | no | calcium ion binding; protein binding | | IQ motif, EF-hand binding site | |
| **PVIT_0010507** | PMUR_0008942 | 1.75 | 0.27 | 0.16 | yes | yes | no | no | | | | |
| **PVIT_0008293** | PMUR_0013073 | 1.73 | 0.21 | 0.12 | yes | yes | no | no | serine-type endopeptidase activity; proteolysis; peptidase activity; membrane; integral component of membrane | serine-type endopeptidase activity; proteolysis | Serine proteases, trypsin domain; Peptidase S1A, chymotrypsin family; Peptidase S1, PA clan | Serine proteases, trypsin domain; Peptidase S1A, chymotrypsin family; Peptidase S1, PA clan |
| PVIT_0008070 | PMUR_0008781 | 1.63 | 0.26 | 0.16 | no | no | no | no | | | | |
| **PVIT_0020870** | PMUR_0012887 | 1.55 | 0.17 | 0.11 | no | no | no | no | | nucleic acid binding | | |
| PVIT_0010475 | PMUR_0008920 | 1.51 | 0.18 | 0.12 | no | no | no | no | | | | |
| **PVIT_0006861** | PMUR_0002287 | 1.50 | 0.04 | 0.02 | no | yes | no | no | | | | |
| **PVIT_0017528** | PMUR_0013113 | 1.48 | 0.36 | 0.25 | yes | yes | yes | yes | | | | |
| **PVIT_0018321** | PMUR_0012509 | 1.46 | 0.16 | 0.11 | no | no | no | no | nucleic acid binding; DNA binding | nucleic acid binding; DNA binding | | |
| **PVIT_0012276** | PMUR_0003087 | 1.45 | 0.26 | 0.18 | yes | yes | no | yes | | | | |
| PVIT_0003436 | PMUR_0012534 | 1.44 | 0.09 | 0.06 | no | no | no | no | kinase activity; phosphorylation | | | |
| **PVIT_0013507** | PMUR_0010401 | 1.43 | 0.15 | 0.11 | yes | yes | no | no | | | | |
| **PVIT_0001143** | PMUR_0012041 | 1.42 | 0.19 | 0.13 | no | no | no | no | nucleotide binding; nuclear exosome (RNase complex); nucleic acid binding; catalytic activity; exonuclease activity; intracellular; nucleobase-containing compound metabolic process; RNA processing; 3'-5' exonuclease activity; cellular metabolic process; nucleic acid phosphodiester bond hydrolysis | nucleotide binding; nuclear exosome (RNase complex); nucleic acid binding; catalytic activity; exonuclease activity; intracellular; nucleobase-containing compound metabolic process; RNA processing; 3'-5' exonuclease activity; cellular metabolic process; nucleic acid phosphodiester bond hydrolysis | | |
| PVIT_0017527 | PMUR_0012778 | 1.34 | 0.26 | 0.19 | yes | yes | no | yes | | | | |
| **PVIT_0003150** | PMUR_0004214 | 1.33 | 0.15 | 0.11 | no | no | no | no | protein phosphorylation; motile cilium | protein phosphorylation | | |
| **PVIT_0011823** | PMUR_0004836 | 1.32 | 0.16 | 0.12 | no | yes | no | no | serine-type endopeptidase activity; proteolysis; peptidase activity | serine-type endopeptidase activity; proteolysis | Serine proteases, trypsin domain; Peptidase S1, PA clan | Serine proteases, trypsin domain; Peptidase S1, PA clan |
| **PVIT_0018318** | PMUR_0008586 | 1.31 | 0.04 | 0.03 | yes | yes | yes | yes | | | | |
| **PVIT_0013263** | PMUR_0012073 | 1.30 | 0.10 | 0.08 | yes | no | yes | no | | | | |
| **PVIT_0013728** | PMUR_0005019 | 1.29 | 0.04 | 0.03 | no | no | no | no | membrane | membrane | | |
| PVIT_0016143 | PMUR_0008646 | 1.28 | 0.15 | 0.12 | no | no | no | no | | | | |
| PVIT_0006445 | PMUR_0001691 | 1.26 | 0.44 | 0.35 | no | no | no | no | | | | |
| **PVIT_0006221** | PMUR_0002524 | 1.26 | 0.04 | 0.03 | no | no | no | no | integral component of membrane | membrane | | |
| PVIT_0012094 | PMUR_0008697 | 1.24 | 0.52 | 0.42 | no | no | no | no | | | | |
| **PVIT_0003960** | PMUR_0000032 | 1.24 | 0.07 | 0.06 | no | no | no | no | | | | |
| **PVIT_0010797** | PMUR_0006150 | 1.23 | 0.16 | 0.13 | no | yes | no | no | | | | |
| **PVIT_0010792** | PMUR_0006148 | 1.22 | 0.19 | 0.16 | yes | yes | no | no | | | | |
| **PVIT_0001002** | PMUR_0011244 | 1.21 | 0.07 | 0.06 | no | no | no | no | membrane; integral component of membrane | | | |
| **PVIT_0001929** | PMUR_0004106 | 1.20 | 0.13 | 0.11 | yes | no | no | no | | | | |
| **PVIT_0010451** | PMUR_0008115 | 1.20 | 0.30 | 0.25 | no | no | no | no | | | | |
| PVIT_0018773 | PMUR_0008030 | 1.20 | 0.20 | 0.16 | no | no | no | no | | | | |
| PVIT_0027098 | PMUR_0013178 | 1.20 | 0.02 | 0.02 | no | no | no | no | | | | |
| PVIT_0000387 | PMUR_0005950 | 1.19 | 0.56 | 0.47 | no | no | no | no | nucleic acid binding | | | |
| **PVIT_0021601** | PMUR_0012745 | 1.19 | 0.16 | 0.14 | yes | yes | no | no | | | | |
| **PVIT_0015335** | PMUR_0012557 | 1.18 | 0.31 | 0.26 | yes | yes | yes | no | | | | |
| PVIT_0013390 | PMUR_0009670 | 1.18 | 0.24 | 0.20 | no | no | no | no | | | | |
| **PVIT_0016508** | PMUR_0006959 | 1.17 | 0.17 | 0.15 | yes | yes | yes | yes | | | | |
| **PVIT_0018732** | PMUR_0003335 | 1.17 | 0.21 | 0.18 | no | no | no | no | | | | |

| | | | | | | | | | | | | |
|---|---|---|---|---|---|---|---|---|---|---|---|---|
| **PVIT_0001362** | PMUR_0003678 | 1.17 | 0.14 | 0.12 | no | no | no | no | | | | |
| PVIT_0003301 | PMUR_0012893 | 1.16 | 0.30 | 0.26 | no | no | no | no | | | | |
| PVIT_0009061 | PMUR_0010883 | 1.16 | 0.38 | 0.32 | no | no | no | no | nucleic acid binding; nucleus; DNA integration | nucleus | | Chromo/chromo shadow domain; Chromo domain; Chromo-like domain superfamily |
| **PVIT_0005694** | PMUR_0011772 | 1.16 | 0.23 | 0.20 | no | no | no | no | | | | |
| **PVIT_0019930** | PMUR_0012761 | 1.16 | 0.25 | 0.22 | no | no | no | no | extracellular region; proteolysis; hydrolase activity | | | |
| **PVIT_0024609** | PMUR_0010001 | 1.15 | 0.27 | 0.24 | no | no | no | no | | | | |
| PVIT_0006028 | PMUR_0011692 | 1.15 | 0.22 | 0.19 | no | no | no | no | | | | |
| PVIT_0003473 | PMUR_0009677 | 1.14 | 0.21 | 0.19 | no | no | no | no | | nucleotide binding; ATP binding; hydrolase activity | | P-loop containing nucleoside triphosphate hydrolase |
| **PVIT_0002228** | PMUR_0008345 | 1.13 | 0.06 | 0.05 | no | no | no | no | tRNA modification; tRNA (guanine-N7-)-methyltransferase activity; RNA (guanine-N7)-methylation; tRNA methyltransferase complex | tRNA modification; tRNA (guanine-N7-)-methyltransferase activity; RNA (guanine-N7)-methylation; tRNA methyltransferase complex | tRNA (guanine-N-7) methyltransferase, Trmb type | tRNA (guanine-N-7) methyltransferase, Trmb type |
| **PVIT_0001383** | PMUR_0003688 | 1.13 | 0.14 | 0.13 | yes | no | yes | no | | | | |
| **PVIT_0010026** | PMUR_0012725 | 1.13 | 0.21 | 0.19 | yes | yes | yes | yes | | | | |
| **PVIT_0007026** | PMUR_0012532 | 1.13 | 0.08 | 0.07 | yes | no | no | no | serine-type endopeptidase inhibitor activity; protein binding; proteolysis; peptidase activity; negative regulation of serine-type endopeptidase activity | protein binding; proteolysis; peptidase activity | Kazal domain | Kazal domain |
| **PVIT_0005735** | PMUR_0008108 | 1.13 | 0.17 | 0.15 | yes | no | no | no | | | | |
| **PVIT_0001577** | PMUR_0003648 | 1.12 | 0.25 | 0.22 | no | no | no | no | | | | |
| **PVIT_0001521** | PMUR_0003838 | 1.12 | 0.06 | 0.06 | no | no | no | no | | | | |
| PVIT_0020910 | PMUR_0011234 | 1.10 | 0.17 | 0.16 | no | no | no | no | | | | |
| PVIT_0022528 | PMUR_0007267 | 1.10 | 0.21 | 0.19 | no | no | no | no | | | | |
| **PVIT_0018709** | PMUR_0011974 | 1.09 | 0.19 | 0.18 | yes | yes | yes | yes | | | | |
| PVIT_0012353 | PMUR_0004501 | 1.09 | 0.38 | 0.35 | no | no | no | no | | | | |
| **PVIT_0003385** | PMUR_0003939 | 1.08 | 0.14 | 0.13 | no | yes | no | no | membrane; integral component of membrane | membrane; integral component of membrane | | |
| **PVIT_0006586** | PMUR_0009062 | 1.08 | 0.34 | 0.32 | yes | yes | no | no | | | CAP domain | CAP domain |
| PVIT_0016675 | PMUR_0009646 | 1.08 | 0.11 | 0.10 | no | no | no | no | | | | |
| PVIT_0011285 | PMUR_0012009 | 1.06 | 0.13 | 0.12 | no | no | no | no | | | | |
| **PVIT_0005562** | PMUR_0000823 | 1.06 | 0.17 | 0.16 | no | no | no | no | | | | |
| **PVIT_0018948** | PMUR_0011315 | 1.06 | 0.18 | 0.17 | no | no | no | no | nucleic acid binding; DNA integration | | | |
| **PVIT_0017047** | PMUR_0009702 | 1.06 | 0.25 | 0.24 | no | no | no | no | | | | |
| **PVIT_0017162** | PMUR_0006259 | 1.05 | 0.06 | 0.06 | no | no | no | no | protein phosphorylation | protein phosphorylation | | |
| PVIT_0025603 | PMUR_0012836 | 1.05 | 0.30 | 0.28 | no | no | no | no | | | | |
| **PVIT_0001524** | PMUR_0003841 | 1.05 | 0.08 | 0.07 | no | no | no | no | membrane; integral component of membrane | | | |
| **PVIT_0022516** | PMUR_0011087 | 1.05 | 0.11 | 0.11 | yes | yes | yes | yes | | | | |
| **PVIT_0020918** | PMUR_0006134 | 1.05 | 0.12 | 0.11 | yes | no | no | no | | | | |
| **PVIT_0018905** | PMUR_0009659 | 1.04 | 0.14 | 0.13 | yes | no | yes | no | | | | |
| **PVIT_0005339** | PMUR_0001275 | 1.04 | 0.12 | 0.12 | no | no | no | no | microtubule cytoskeleton organization; microtubule; microtubule binding | | | |
| **PVIT_0015820** | PMUR_0006188 | 1.04 | 0.14 | 0.14 | no | yes | no | no | | | | |
| **PVIT_0015154** | PMUR_0004184 | 1.03 | 0.24 | 0.23 | yes | no | yes | no | | | | |
| **PVIT_0001387** | PMUR_0003693 | 1.02 | 0.12 | 0.12 | yes | yes | yes | yes | | | | |
| **PVIT_0017599** | PMUR_0013161 | 1.02 | 0.10 | 0.10 | yes | yes | no | no | polysaccharide catabolic process; hydrolase activity, hydrolyzing O-glycosyl compounds; extracellular region; carbohydrate metabolic process; metabolic process; cellulase activity; hydrolase activity; hydrolase activity, acting on glycosyl bonds; cellulose binding | polysaccharide catabolic process; hydrolase activity, hydrolyzing O-glycosyl compounds; carbohydrate metabolic process; metabolic process; cellulase activity; hydrolase activity; hydrolase activity, acting on glycosyl bonds | Glycoside hydrolase family 12; Glycoside hydrolase family 11/12; Concanavalin A-like lectin/glucanase domain superfamily | Phosphatidylinositol-specific phospholipase C, X domain; Glycoside hydrolase family 12; Glycoside hydrolase family 11/12; Concanavalin A-like lectin/glucanase domain superfamily |
| PVIT_0015339 | PMUR_0008381 | 1.01 | 0.07 | 0.07 | no | no | no | no | | | | |
| **PVIT_0002656** | PMUR_0013156 | 1.01 | 0.13 | 0.13 | no | no | no | no | | | | |
| **PVIT_0011304** | PMUR_0007264 | 1.01 | 0.13 | 0.13 | yes | no | yes | no | | | | |
| **PVIT_0006307** | PMUR_0009093 | 1.01 | 0.11 | 0.11 | yes | no | yes | no | | | | |
| **PVIT_0003402** | PMUR_0003979 | 1.00 | 0.31 | 0.31 | yes | yes | yes | yes | | | | |
| **PVIT_0024814** | PMUR_0012891 | 1.00 | 0.19 | 0.19 | yes | no | no | no | protein phosphorylation; membrane; integral component of membrane | | HNH nuclease | |
| **PVIT_0012948** | PMUR_0001040 | 1.00 | 0.09 | 0.09 | no | no | no | no | | | | |

*Pl. viticola* genes found expressed in the RNA-seq data are in bold.

Supplementary table 12 *Pl. viticola* genes with a significant signal in the branch-site test

| Tested gene | Orthogroup | Align. size | Delta LRT | *P*-value | Q-value | Secreted | GO terms | InterPro domains |
|---|---|---|---|---|---|---|---|---|
| **PVIT_0006791** | Phyca11_562426, PITG_00022T0, KUF92812, Phal03539, PMUR_0004720, PVIT_0006791 | 969 | 149.74 | 1.98E-34 | 7.95E-31 | no | drug transmembrane transport; drug transmembrane transporter activity; antiporter activity; membrane; transmembrane transporter activity; transmembrane transport | Multi antimicrobial extrusion protein |
| **PVIT_0007111** | Phyca11_7933, PITG_17333T0, KUF77828, Phal11456, PMUR_0011460, PVIT_0007111 | 1593 | 118.00 | 1.73E-27 | 3.48E-24 | no | protein kinase activity; protein serine/threonine kinase activity; ATP binding; protein phosphorylation | Serine/threonine-protein kinase, active site; Protein kinase domain; Protein kinase-like domain superfamily |
| **PVIT_0005972** | Phyca11_123187, PITG_11886T0, KUF89757, Phal04063, PMUR_0009113, PVIT_0005972 | 969 | 94.54 | 2.40E-22 | 3.21E-19 | no | protein binding; cell; glycerol ether metabolic process; protein disulfide oxidoreductase activity; cell redox homeostasis; oxidation-reduction process | Ubiquitin domain; PUB domain; Ubiquilin |
| **PVIT_0020511** | Phyca11_107403, PITG_10896T0, KUF79858, Phal09861, PMUR_0008422, PVIT_0020511 | 1446 | 90.60 | 1.76E-21 | 1.77E-18 | no | kinase activity; phosphorylation | |
| **PVIT_0001212** | Phyca11_545532, PITG_02671T0, KUF87724, Phal06919, PMUR_0001253, PVIT_0001212 | 756 | 79.58 | 4.64E-19 | 3.72E-16 | no | cAMP-dependent protein kinase complex; cAMP-dependent protein kinase regulator activity; cAMP binding; regulation of protein kinase activity | Cyclic nucleotide-binding, conserved site; Cyclic nucleotide-binding domain; RmlC-like jelly roll fold; Cyclic nucleotide-binding-like |
| **PVIT_0019913** | Phyca11_528570, PITG_22924T0, KUF86048, Phal07896, PMUR_0008817, PVIT_0019913 | 1074 | 62.00 | 3.44E-15 | 2.31E-12 | no | membrane | |
| **PVIT_0003711** | Phyca11_505784, PITG_21524T0, KUF78485, Phal07365, PMUR_0000200, PVIT_0003711 | 1539 | 61.24 | 5.05E-15 | 2.90E-12 | no | monooxygenase activity; iron ion binding; oxidoreductase activity, acting on paired donors, with incorporation or reduction of molecular oxygen; lyase activity; heme binding; oxidation-reduction process | Cytochrome P450, conserved site; Cytochrome P450; Cytochrome P450, E-class, group I |
| **PVIT_0001295** | Phyca11_115737, PITG_02802T0, KUF79445, Phal11391, PMUR_0000958, PVIT_0001295 | 885 | 60.73 | 6.54E-15 | 3.28E-12 | no | membrane | Band 7 domain; Stomatin family |
| **PVIT_0013045** | Phyca11_7396, PITG_01416T0, KUF81499, Phal14135, PMUR_0001125, PVIT_0013045 | 1419 | 58.93 | 1.63E-14 | 7.28E-12 | no | protein binding; cell; anatomical structure morphogenesis; tissue development; single-organism cellular process; animal organ development; establishment of localization in cell | Tetratricopeptide repeat-containing domain; Tetratricopeptide-like helical domain superfamily; Tetratricopeptide repeat |
| **PVIT_0009562** | Phyca11_9865, PITG_11573T0, KUF85728, Phal01751, PMUR_0002096, PVIT_0009562 | 1038 | 58.26 | 2.30E-14 | 9.22E-12 | no | proteolysis; peptidase activity | Serine aminopeptidase, S33 |
| **PVIT_0005945** | Phyca11_19592, PITG_11966T0, KUF85566, Phal08576, PMUR_0006626, PVIT_0005945 | 1683 | 55.19 | 1.09E-13 | 3.99E-11 | no | ATP binding; cytoplasm; mitochondrion; protein refolding | Chaperonin Cpn60; Chaperonin Cpn60/TCP-1 family; GroEL-like apical domain superfamily; GroEL-like equatorial domain superfamily |
| **PVIT_0021795** | Phyca11_503134, PITG_19668T0, KUF79961, Phal09173, PMUR_0010414, PVIT_0021795 | 555 | 54.20 | 1.81E-13 | 6.07E-11 | no | dihydrolipoyllysine-residue acetyltransferase activity; mitochondrial matrix; ribosome; pyruvate metabolic process; metabolic process; transferase activity; transferase activity, transferring acyl groups; pyruvate dehydrogenase complex | Biotin/lipoyl attachment; Single hybrid motif |
| **PVIT_0011454** | Phyca11_511108, PITG_04545T0, KUF79947, Phal08124, PMUR_0011394, PVIT_0011454 | 894 | 53.55 | 2.52E-13 | 7.79E-11 | no | GTPase activity; GTP binding; ribosome biogenesis | GTP binding domain; GTP-binding protein, orthogonal bundle domain superfamily; P-loop containing nucleoside triphosphate hydrolase |
| **PVIT_0020495** | Phyca11_70774, PITG_19530T0, KUF82231, Phal13118, PMUR_0008410, PVIT_0020495 | 399 | 53.39 | 2.74E-13 | 7.86E-11 | no | nucleus; protein heterodimerization activity | Transcription factor CBF/NF-Y/archaeal histone domain; Histone-fold |

| Tested gene | Orthogroup | Align. size | Delta LRT | *P*-value | Q-value | Secreted | GO terms | InterPro domains |
|---|---|---|---|---|---|---|---|---|
| **PVIT_0011740** | Phyca11_554424, PITG_21176T0, KUF78349, Phal04006, PMUR_0004934, PVIT_0011740 | 1770 | 44.88 | 2.10E-11 | 5.61E-09 | no | protein import into nucleus, translocation; protein serine/threonine kinase activity; binding; cytoplasm; protein phosphorylation; NLS-bearing protein import into nucleus; ribosomal protein import into nucleus; intracellular protein transport; nuclear localization sequence binding; Ran GTPase binding; protein transporter activity; nuclear membrane; nuclear periphery | Importin-beta, N-terminal domain; Armadillo-like helical; Armadillo-type fold |
| **PVIT_0007405** | Phyca11_562095, PITG_02022T0, KUF94557, Phal00740, PMUR_0007974, PVIT_0007405 | 5097 | 44.13 | 3.07E-11 | 7.71E-09 | no | structural molecule activity; binding; protein binding; intracellular protein transport; integral component of membrane; vesicle-mediated transport; clathrin coat of trans-Golgi network vesicle; clathrin coat of coated pit | Clathrin, heavy chain, linker, core motif; Clathrin, heavy chain; Tetratricopeptide-like helical domain superfamily; Armadillo-type fold; Clathrin heavy chain, N-terminal; Clathrin, heavy chain/VPS, 7-fold repeat; Clathrin, heavy chain, propeller repeat |
| **PVIT_0006426** | Phyca11_506258, PITG_12916T0, KUF84984, Phal06271, PMUR_0011524, PVIT_0006426 | 1053 | 43.22 | 4.89E-11 | 1.16E-08 | yes | cysteine-type endopeptidase activity; extracellular space; lysosome; proteolysis; cysteine-type peptidase activity; proteolysis involved in cellular protein catabolic process | Cysteine peptidase, cysteine active site; Cysteine peptidase, histidine active site; Cysteine peptidase, asparagine active site; Peptidase C1A, papain C-terminal; Peptidase C1A |
| **PVIT_0011623** | Phyca11_508456, PITG_13718T0, KUF90555, Phal11080, PMUR_0011240, PVIT_0011623 | 1239 | 39.55 | 3.19E-10 | 7.13E-08 | no | copper ion transmembrane transporter activity; integral component of membrane; metal ion transport; copper ion transmembrane transport; metal ion binding | Heavy metal-associated domain, HMA; Ctr copper transporter |
| **PVIT_0011236** | Phyca11_511517, PITG_16916T0, KUF84073, Phal10179, PMUR_0012427, PVIT_0011236 | 1008 | 38.11 | 6.69E-10 | 1.41E-07 | no | protein kinase activity; protein serine/threonine kinase activity; ATP binding; protein phosphorylation | Protein kinase, ATP binding site; Protein kinase domain; Serine-threonine/tyrosine-protein kinase, catalytic domain; Protein kinase-like domain superfamily |
| **PVIT_0017234** | Phyca11_9084, PITG_04000T0, KUF77695, Phal06372, PMUR_0001948, PVIT_0017234 | 990 | 35.35 | 2.76E-09 | 5.55E-07 | no | protein binding | WD40 repeat, conserved site; WD40-repeat-containing domain; WD40/YVTN repeat-like-containing domain superfamily; WD40 repeat; G-protein beta WD-40 repeat |
| **PVIT_0017715** | Phyca11_550917, PITG_15026T0, KUF84755, Phal02249, PMUR_0012296, PVIT_0017715 | 2442 | 34.50 | 4.27E-09 | 8.17E-07 | no | protein import into nucleus, translocation; binding; cytoplasm; NLS-bearing protein import into nucleus; ribosomal protein import into nucleus; nuclear localization sequence binding; protein transporter activity; nuclear membrane; nuclear periphery | Armadillo-like helical; Armadillo-type fold |
| **PVIT_0011847** | Phyca11_38642, PITG_13615T0, KUF82271, Phal04750, PMUR_0004845, PVIT_0011847 | 726 | 31.62 | 1.87E-08 | 3.42E-06 | no | rRNA modification; rRNA (adenine-N6,N6-)-dimethyltransferase activity; cytoplasm; rRNA methyltransferase activity; rRNA methylation | Ribosomal RNA adenine methylase transferase, conserved site; Ribosomal RNA adenine methylase transferase, N-terminal; Ribosomal RNA adenine methyltransferase KsgA/Erm |
| **PVIT_0013056** | Phyca11_70439, PITG_01448T0, KUF81503, Phal06276, PMUR_0006342, PVIT_0013056 | 384 | 29.23 | 6.43E-08 | 1.12E-05 | no | calcium ion binding | EF-Hand 1, calcium-binding site; EF-hand domain; EF-hand domain pair |
| **PVIT_0017784** | Phyca11_7234, PITG_17975T0, KUF79813, Phal00359, PMUR_0007002, PVIT_0017784 | 1161 | 28.72 | 8.37E-08 | 1.40E-05 | no | protein binding; transferase activity; transferase activity, transferring glycosyl groups | GAF domain |
| **PVIT_0001644** | Phyca11_116367, PITG_00397T0, KUF78113, Phal01183, PMUR_0000545, PVIT_0001644 | 2541 | 26.73 | 2.34E-07 | 3.76E-05 | no | formation of translation preinitiation complex; RNA binding; translation initiation factor activity; protein binding; eukaryotic translation initiation factor 3 complex; translational initiation; regulation of translational initiation; eukaryotic 43S preinitiation complex; translation initiation factor binding; eukaryotic 48S preinitiation complex | Proteasome component (PCI) domain; Eukaryotic translation initiation factor 3 subunit C, N-terminal domain; ArsR-like helix-turn-helix domain; Eukaryotic translation initiation factor 3 subunit C |
| **PVIT_0015346** | Phyca11_21290, PITG_07455T0, KUF88067, Phal01518, PMUR_0008383, PVIT_0015346 | 3525 | 26.64 | 2.45E-07 | 3.79E-05 | no | nucleotide binding; magnesium ion binding; phospholipid-translocating ATPase activity; ATP binding; plasma membrane; phospholipid transport; integral component of membrane; phospholipid translocation; metal ion binding | P-type ATPase; P-type ATPase, subfamily IV; P-type ATPase, A domain superfamily; HAD superfamily; P-type ATPase, cytoplasmic domain N; P-type ATPase, phosphorylation site |
| **PVIT_0018929** | Phyca11_537183, PITG_11789T0, KUF82256, Phal01544, PMUR_0004550, PVIT_0018929 | 693 | 26.34 | 2.86E-07 | 4.10E-05 | no | proton transport; vacuolar proton-transporting V-type ATPase complex; hydrolase activity, acting on acid anhydrides, catalyzing transmembrane movement of substances; transmembrane transport | Coiled-coil domain containing protein 109, C-terminal |

| Tested gene | Orthogroup | Align. size | Delta LRT | *P* -value | Q-value | Secreted | GO terms | InterPro domains |
|---|---|---|---|---|---|---|---|---|
| **PVIT_0019167** | Phyca11_547133, PITG_08661T0, KUG02176, Phal08958, PMUR_0011983, PVIT_0019167 | 1476 | 26.38 | 2.81E-07 | 4.10E-05 | no | catalytic activity; RNA modification; transferase activity; protein kinase regulator activity; methylthiotransferase activity; tRNA methylthiolation; macromolecule modification; regulation of protein kinase activity; iron-sulfur cluster binding; 4 iron, 4 sulfur cluster binding | Methylthiotransferase, conserved site; Elp3/MiaB/NifB; Radical SAM; Methylthiotransferase, N-terminal; Methylthiotransferase; Radical SAM, alpha/beta horseshoe |
| **PVIT_0018786** | Phyca11_504262, PITG_19871T0, KUF80558, Phal01578, PMUR_0006571, PVIT_0018786 | 2388 | 26.05 | 3.33E-07 | 4.62E-05 | no | ATP binding; hydrolase activity; cell division | ATPase, AAA-type, conserved site; CDC48, N-terminal subdomain; AAA+ ATPase domain; ATPase, AAA-type, core; CDC48, domain 2; Vps4 oligomerisation, C-terminal; AAA ATPase, CDC48 family; Aspartate decarboxylase-like domain superfamily; P-loop containing nucleoside triphosphate hydrolase |
| **PVIT_0001977** | Phyca11_509876, PITG_05224T0, KUG01522, Phal04316, PMUR_0007898, PVIT_0001977 | 423 | 25.01 | 5.71E-07 | 7.64E-05 | no | microtubule motor activity; protein binding; ATP binding; kinesin complex; microtubule-based movement; microtubule binding; ATPase activity | Kinesin motor domain; SMAD/FHA domain superfamily; P-loop containing nucleoside triphosphate hydrolase |
| **PVIT_0012980** | Phyca11_534012, PITG_01330T0, KUF81212, Phal11222, PMUR_0001069, PVIT_0012980 | 1365 | 24.35 | 8.05E-07 | 1.04E-04 | no | integral component of membrane; transferase activity, transferring acyl groups | Membrane bound O-acyl transferase, MBOAT |
| **PVIT_0003670** | Phyca11_564754, PITG_15837T0, KUF80592, Phal08051, PMUR_0004283, PVIT_0003670 | 1401 | 23.47 | 1.27E-06 | 1.59E-04 | no | ATP binding; nucleus | SNF2-related, N-terminal domain; Chromo/chromo shadow domain; Helicase, C-terminal; Helicase superfamily 1/2, ATP-binding domain; Chromo domain; Chromo-like domain superfamily; P-loop containing nucleoside triphosphate hydrolase |
| **PVIT_0010101** | Phyca11_542607, PITG_02114T0, KUF76904, Phal04605, PMUR_0001777, PVIT_0010101 | 1383 | 23.14 | 1.51E-06 | 1.83E-04 | no | aminopeptidase activity; proteolysis; metallopeptidase activity; zinc ion binding | Peptidase M18; Peptidase M18, domain 2 |
| **PVIT_0018284** | Phyca11_34281, PITG_11759T0, KUF76521, Phal08738, PMUR_0009478, PVIT_0018284 | 1677 | 22.68 | 1.92E-06 | 2.27E-04 | no | protein binding | Ankyrin repeat-containing domain; Ankyrin repeat |
| **PVIT_0000878** | Phyca11_525004, PITG_02268T0, KUF88600, Phal00471, PMUR_0004458, PVIT_0000878 | 825 | 22.34 | 2.28E-06 | 2.62E-04 | no | integral component of membrane | |
| **PVIT_0016179** | Phyca11_561150, PITG_15735T0, KUF73665, Phal06406, PMUR_0002604, PVIT_0016179 | 1698 | 21.45 | 3.63E-06 | 3.94E-04 | no | protein kinase activity; protein serine/threonine kinase activity; protein binding; ATP binding; nucleus; cytoplasm; protein phosphorylation; intracellular signal transduction | Protein kinase, ATP binding site; Protein kinase domain; AGC-kinase, C-terminal; Pleckstrin homology domain; Ankyrin repeat-containing domain; Protein kinase-like domain superfamily; PH-like domain superfamily; Ankyrin repeat |
| **PVIT_0019500** | Phyca11_533155, PITG_08734T0, KUF75663, Phal04137, PMUR_0002790, PVIT_0019500 | 2757 | 21.46 | 3.61E-06 | 3.94E-04 | no | ion channel activity; voltage-gated chloride channel activity; chloride transport; membrane; integral component of membrane; ion transmembrane transport; transmembrane transport; regulation of anion transmembrane transport | CBS domain; Chloride channel, voltage gated; Chloride channel, core |
| **PVIT_0019185** | Phyca11_113258, PITG_08676T0, KUG02150, Phal11229, PMUR_0010308, PVIT_0019185 | 933 | 20.80 | 5.11E-06 | 5.40E-04 | no | proteasome complex; protein binding; nucleus; cytosol; glycolytic process; N-terminal protein myristoylation; protein targeting to vacuole; fatty acid beta-oxidation; water transport; hyperosmotic response; Golgi organization; response to temperature stimulus; photomorphogenesis; photorespiration; kinase activity; proteasome-mediated ubiquitin-dependent protein catabolic process; proteasome assembly; response to cadmium ion; response to misfolded protein | JAB1/MPN/MOV34 metalloenzyme domain; Rpn11/EIF3F, C-terminal |
| **PVIT_0006624** | Phyca11_571282, PITG_11102T0, KUF76508, Phal02941, PMUR_0003642, PVIT_0006624 | 1812 | 20.68 | 5.43E-06 | 5.59E-04 | no | protein kinase activity; protein serine/threonine kinase activity; ATP binding; protein phosphorylation | Serine/threonine-protein kinase, active site; Protein kinase, ATP binding site; Cyclic nucleotide-binding, conserved site; Cyclic nucleotide-binding domain; Protein kinase domain; AGC-kinase, C-terminal; Protein kinase-like domain superfamily; RmlC-like jelly roll fold; Cyclic nucleotide-binding-like |

| Tested gene | Orthogroup | Align. size | Delta LRT | *P* -value | Q-value | Secreted | GO terms | InterPro domains |
|---|---|---|---|---|---|---|---|---|
| **PVIT_0009795** | Phyca11_534373, PITG_07252T0, KUF76130, Phal00703, PMUR_0009200, PVIT_0009795 | 327 | 20.63 | 5.57E-06 | 5.59E-04 | no | synaptonemal complex; single-stranded DNA binding; ATP binding; chiasma; mismatch repair; ATPase activity; MutLalpha complex | Histidine kinase/HSP90-like ATPase; MutL, C-terminal, dimerisation |
| **PVIT_0010806** | Phyca11_19812, PITG_07902T0, KUF78648, Phal01953, PMUR_0006158, PVIT_0010806 | 1365 | 20.42 | 6.22E-06 | 6.10E-04 | no | nucleus | Chromo/chromo shadow domain; Chromo domain; Chromo-like domain superfamily |
| **PVIT_0001083** | Phyca11_569294, PITG_02550T0, KUF79783, Phal02833, PMUR_0005250, PVIT_0001083 | 4470 | 20.14 | 7.19E-06 | 6.72E-04 | no | transcription factor activity, sequence-specific DNA binding; regulation of transcription, DNA-templated | |
| **PVIT_0006127** | Phyca11_527201, PITG_05560T0, KUF76730, Phal07792, PMUR_0001567, PVIT_0006127 | 1773 | 20.16 | 7.11E-06 | 6.72E-04 | no | integral component of membrane | Transmembrane protein TqsA-like |
| **PVIT_0011106** | Phyca11_552564, PITG_16386T0, KUF83907, Phal00431, PMUR_0002397, PVIT_0011106 | 297 | 19.63 | 9.41E-06 | 8.59E-04 | no | isomerase activity | BolA protein |
| **PVIT_0022488** | Phyca11_510798, PITG_08199T0, KUG01097, Phal02854, PMUR_0006243, PVIT_0022488 | 771 | 19.37 | 1.08E-05 | 9.60E-04 | no | transcription factor activity, sequence-specific DNA binding; nucleus; regulation of transcription, DNA-templated; sequence-specific DNA binding | Heat shock factor (HSF)-type, DNA-binding; ArsR-like helix-turn-helix domain; Heat shock transcription factor family |
| **PVIT_0019045** | Phyca11_48915, PITG_23178T0, KUF89226, Phal07228, PMUR_0007287, PVIT_0019045 | 432 | 18.98 | 1.32E-05 | 1.16E-03 | no | protein binding; proteolysis; peptidase activity; membrane; integral component of membrane | Kazal domain |
| **PVIT_0010430** | Phyca11_571966, PITG_09965T0, KUF94296, Phal07840, PMUR_0008135, PVIT_0010430 | 2079 | 18.83 | 1.43E-05 | 1.22E-03 | no | intracellular protein transport; ER to Golgi vesicle-mediated transport; zinc ion binding; COPII vesicle coat | von Willebrand factor, type A; Zinc finger, Sec23/Sec24-type; Sec23/Sec24, trunk domain; Sec23/Sec24, helical domain; Gelsolin-like domain; Sec23/Sec24 beta-sandwich |
| **PVIT_0019353** | Phyca11_551426, PITG_19608T0, KUF88578, Phal00838, PMUR_0005105, PVIT_0019353 | 432 | 18.79 | 1.46E-05 | 1.22E-03 | no | cleavage involved in rRNA processing; maturation of LSU-rRNA; RNA binding; nucleolus; translation; viral nucleocapsid; cytosolic large ribosomal subunit; rRNA pseudouridine synthesis; snRNA pseudouridine synthesis; box H/ACA snoRNP complex; box H/ACA snoRNA binding | Ribosomal protein L7Ae/L30e/S12e/Gadd45; H/ACA ribonucleoprotein complex, subunit Nhp2, eukaryote; Ribosomal protein L7Ae/L8/Nhp2 family |
| **PVIT_0021040** | Phyca11_506714, PITG_13041T0, KUF76086, Phal06340, PMUR_0006775, PVIT_0021040 | 2190 | 18.45 | 1.74E-05 | 1.43E-03 | no | DNA binding; transcription elongation from RNA polymerase II promoter; histone modification; Cdc73/Paf1 complex; ligase activity | Plus-3 domain |
| **PVIT_0011277** | Phyca11_130345, PITG_19647T0, KUF84328, Phal07232, PMUR_0011200, PVIT_0011277 | 1359 | 17.71 | 2.58E-05 | 2.07E-03 | no | nucleotide binding; protein kinase activity; ATP binding; protein phosphorylation; membrane; integral component of membrane; kinase activity; phosphorylation | Meiosis-specific nuclear structural protein 1 |
| **PVIT_0000670** | Phyca11_8263, PITG_03474T0, KUG02199, Phal03009, PMUR_0000248, PVIT_0000670 | 972 | 17.58 | 2.75E-05 | 2.13E-03 | no | integral component of membrane | Rab-GTPase-TBC domain |
| **PVIT_0018286** | Phyca11_503414, PITG_11755T0, KUF94440, Phal11255, PMUR_0010817, PVIT_0018286 | 1257 | 17.59 | 2.74E-05 | 2.13E-03 | yes | | |
| **PVIT_0009363** | Phyca11_503496, PITG_03788T0, KUF85578, Phal01185, PMUR_0007546, PVIT_0009363 | 738 | 17.30 | 3.19E-05 | 2.42E-03 | no | | |
| **PVIT_0001841** | Phyca11_570517, PITG_00634T0, KUF76976, Phal04188, PMUR_0003890, PVIT_0001841 | 2142 | 16.68 | 4.43E-05 | 3.30E-03 | no | microtubule cytoskeleton organization; spindle pole; equatorial microtubule organizing center; gamma-tubulin complex; structural constituent of cytoskeleton; centrosome; microtubule organizing center; spindle pole body; microtubule nucleation; meiotic nuclear division; cytoplasmic microtubule organization; gamma-tubulin binding; microtubule minus-end binding; centrosome duplication; interphase microtubule nucleation by interphase microtubule organizing center; mitotic spindle assembly | Gamma-tubulin complex component protein |
| **PVIT_0004681** | Phyca11_5374, PITG_14971T0, KUF76285, Phal04784, PMUR_0011370, PVIT_0004681 | 2307 | 16.63 | 4.54E-05 | 3.32E-03 | no | protein kinase activity; ATP binding; protein phosphorylation; membrane; kinase activity; phosphorylation | Protein kinase domain; Protein kinase-like domain superfamily |

| Tested gene | Orthogroup | Align. size | Delta LRT | *P* -value | Q-value | Secreted | GO terms | InterPro domains |
|---|---|---|---|---|---|---|---|---|
| **PVIT_0001940** | Phyca11_118922, PITG_00748T0, KUF96721, Phal12777, PMUR_0004112, PVIT_0001940 | 747 | 14.95 | 1.10E-04 | 7.92E-03 | no | maturation of SSU-rRNA from tricistronic rRNA transcript (SSU-rRNA, 5.8S rRNA, LSU-rRNA); binding; snoRNA binding; 90S preribosome; small-subunit processome; t-UTP complex; positive regulation of transcription from RNA polymerase I promoter | BP28, C-terminal domain; U3 small nucleolar RNA-associated protein 10; Armadillo-like helical; Armadillo-type fold |
| **PVIT_0012672** | Phyca11_546135, PITG_13257T0, KUF80953, Phal03465, PMUR_0010352, PVIT_0012672 | 879 | 14.81 | 1.19E-04 | 8.37E-03 | no | translation initiation factor activity; translational initiation | MaoC-like domain |
| **PVIT_0016699** | Phyca11_553199, PITG_21150T0, KUF78371, Phal08237, PMUR_0010053, PVIT_0016699 | 1422 | 14.45 | 1.44E-04 | 9.98E-03 | no | transport; integral component of membrane | Major facilitator superfamily domain; Folate-biopterin transporter |
| **PVIT_0018593** | Phyca11_511666, PITG_06131T0, KUF87355, Phal04830, PMUR_0004484, PVIT_0018593 | 639 | 14.23 | 1.62E-04 | 1.10E-02 | no | integral component of membrane; posttranslational protein targeting to membrane, translocation; Sec62/Sec63 complex | Translocation protein Sec66 |
| **PVIT_0003408** | Phyca11_13754, PITG_05778T0, KUF93709, Phal00231, PMUR_0010404, PVIT_0003408 | 1722 | 13.95 | 1.87E-04 | 1.25E-02 | no | protein binding | WD40 repeat, conserved site; WD40-repeat-containing domain; WD40/YVTN repeat-like-containing domain superfamily; WD40 repeat; G-protein beta WD-40 repeat |
| **PVIT_0009348** | Phyca11_539800, PITG_03792T0, KUF85576, Phal14413, PMUR_0007554, PVIT_0009348 | 1302 | 13.69 | 2.16E-04 | 1.42E-02 | no | RNA binding; translation initiation factor activity; catalytic activity; cytoplasm; translational initiation; metabolic process; 2,3-bisphosphoglycerate-independent phosphoglycerate mutase activity; metal ion binding | Eukaryotic translation initiation factor 4E (eIF-4E), conserved site; Metalloenzyme; Translation Initiation factor eIF- 4e; 2,3-bisphosphoglycerate-independent phosphoglycerate mutase; Alkaline phosphatase-like, alpha/beta/alpha; Alkaline-phosphatase-like, core domain superfamily; Translation Initiation factor eIF- 4e-like |
| **PVIT_0007471** | Phyca11_532861, PITG_10152T0, KUF83836, Phal14326, PMUR_0001376, PVIT_0007471 | 612 | 13.48 | 2.41E-04 | 1.56E-02 | no | calcium ion binding; integral component of membrane | EF-Hand 1, calcium-binding site; EF-hand domain; EF-hand domain pair |
| **PVIT_0002178** | Phyca11_527718, PITG_05274T0, KUF97665, Phal03178, PMUR_0008321, PVIT_0002178 | 2754 | 13.41 | 2.50E-04 | 1.59E-02 | no | binding; cytoplasm; protein import into nucleus; nuclear localization sequence binding; protein transporter activity; nuclear membrane | Exportin-1/Importin-beta-like; Armadillo-type fold |
| **PVIT_0005329** | Phyca11_103899, PITG_01795T0, KUG00625, Phal07880, PMUR_0001267, PVIT_0005329 | 522 | 13.31 | 2.63E-04 | 1.65E-02 | no | RNA binding | YTH domain |
| **PVIT_0014035** | Phyca11_18813, PITG_10990T0, KUF86772, Phal13704, PMUR_0002981, PVIT_0014035 | 663 | 13.22 | 2.77E-04 | 1.71E-02 | no | dimethylallyltranstransferase activity; geranyltranstransferase activity; cytoplasm; isoprenoid biosynthetic process; integral component of membrane; farnesyl diphosphate biosynthetic process | Polyprenyl synthetase; Isoprenoid synthase domain superfamily |
| **PVIT_0013247** | Phyca11_509394, PITG_16466T0, KUF74988, Phal11912, PMUR_0009691, PVIT_0013247 | 624 | 13.17 | 2.84E-04 | 1.73E-02 | no | methyltransferase activity; transferase activity; methylation | |
| **PVIT_0002890** | Phyca11_526931, PITG_10461T0, KUF98786, Phal09730, PMUR_0004401, PVIT_0002890 | 2895 | 13.03 | 3.07E-04 | 1.84E-02 | no | protein folding in endoplasmic reticulum; ER membrane protein complex | ER membrane protein complex subunit 1, C-terminal; ER membrane protein complex subunit 1; Quinoprotein alcohol dehydrogenase-like superfamily; Pyrrolo-quinoline quinone repeat |
| **PVIT_0007430** | Phyca11_532846, PITG_02048T0, KUF89727, Phal06707, PMUR_0010425, PVIT_0007430 | 687 | 12.99 | 3.14E-04 | 1.85E-02 | no | nucleus; tricarboxylic acid cycle; DNA replication; transferase activity; transferase activity, transferring acyl groups, acyl groups converted into alkyl on transfer | DNA polymerase subunit Cdc27 |
| **PVIT_0014721** | Phyca11_511401, PITG_18434T0, KUF80971, Phal00587, PMUR_0009450, PVIT_0014721 | 573 | 12.91 | 3.27E-04 | 1.91E-02 | no | | B-block binding subunit of TFIIIC |
| **PVIT_0008239** | Phyca11_510870, PITG_06740T0, KUF88260, Phal13237, PMUR_0006086, PVIT_0008239 | 648 | 12.80 | 3.47E-04 | 1.99E-02 | no | kinase activity; phosphorylation | PH-like domain superfamily |
| **PVIT_0001411** | Phyca11_548533, PITG_02952T0, KUF66944, Phal10413, PMUR_0008440, PVIT_0001411 | 1200 | 12.65 | 3.76E-04 | 2.10E-02 | no | protein binding; kinase activity; phosphorylation | Tetratricopeptide repeat-containing domain; Tetratricopeptide-like helical domain superfamily; Tetratricopeptide repeat 1; Tetratricopeptide repeat |

| Tested gene | Orthogroup | Align. size | Delta LRT | *P*-value | Q-value | Secreted | GO terms | InterPro domains |
|---|---|---|---|---|---|---|---|---|
| **PVIT_0001830** | Phyca11_118786, PITG_00620T0, KUF76974, Phal03586, PMUR_0003899, PVIT_0001830 | 888 | 12.65 | 3.76E-04 | 2.10E-02 | no | microtubule motor activity; binding; ATP binding; microtubule-based movement; microtubule binding; hydrolase activity | Kinesin motor domain; P-loop containing nucleoside triphosphate hydrolase |
| **PVIT_0004386** | Phyca11_560447, PITG_09437T0, KUF76756, Phal05034, PMUR_0007734, PVIT_0004386 | 306 | 12.49 | 4.10E-04 | 2.26E-02 | no | deadenylation-dependent decapping of nuclear-transcribed mRNA; RNA cap binding; P-body; nucleus | LSM domain, eukaryotic/archaea-type; LSM domain superfamily |
| **PVIT_0013244** | Phyca11_511177, PITG_16468T0, KUF74967, Phal03634, PMUR_0009687, PVIT_0013244 | 1887 | 12.44 | 4.21E-04 | 2.29E-02 | no | ATP binding; phosphorylation; inositol pentakisphosphate 2-kinase activity; intracellular part; single-organism process | Inositol-pentakisphosphate 2-kinase |
| **PVIT_0002656** | Phyca11_526539, PITG_06232T0, KUF83513, Phal11618, PMUR_0013156, PVIT_0002656 | 153 | 12.40 | 4.30E-04 | 2.31E-02 | no | | |
| **PVIT_0006578** | Phyca11_571248, PITG_11063T0, KUF82691, Phal07237, PMUR_0010212, PVIT_0006578 | 936 | 12.30 | 4.54E-04 | 2.40E-02 | no | proteasome complex; protein binding; proteasome-mediated ubiquitin-dependent protein catabolic process | JAB1/MPN/MOV34 metalloenzyme domain; Rpn11/EIF3F, C-terminal |
| **PVIT_0012117** | Phyca11_573771, PITG_13918T0, KUF81328, Phal13689, PMUR_0006408, PVIT_0012117 | 2349 | 12.20 | 4.77E-04 | 2.49E-02 | no | peptide alpha-N-acetyltransferase activity; protein binding; cytoplasm; N-terminal peptidyl-methionine acetylation | Tetratricopeptide repeat-containing domain; N-terminal acetyltransferase A, auxiliary subunit; Tetratricopeptide-like helical domain superfamily; Tetratricopeptide repeat 2; Tetratricopeptide repeat |
| **PVIT_0011519** | Phyca11_561723, PITG_04645T0, KUG00480, Phal08717, PMUR_0002719, PVIT_0011519 | 261 | 12.02 | 5.27E-04 | 2.71E-02 | no | maintenance of transcriptional fidelity during DNA-templated transcription elongation from RNA polymerase II promoter; DNA-directed 5'-3' RNA polymerase activity; binding; DNA-directed RNA polymerase II, core complex; transcription-coupled nucleotide-excision repair; transcription initiation from RNA polymerase II promoter | DNA-directed RNA polymerase M, 15kDa subunit, conserved site; DNA-directed RNA polymerase, M/15kDa subunit |
| **PVIT_0002330** | Phyca11_536953, PITG_06358T0, KUG00093, Phal04000, PMUR_0006800, PVIT_0002330 | 6957 | 11.93 | 5.51E-04 | 2.80E-02 | no | 1,3-beta-D-glucan synthase complex; 1,3-beta-D-glucan synthase activity; (1->3)-beta-D-glucan biosynthetic process; membrane; integral component of membrane; transmembrane transporter activity; transmembrane transport | Major facilitator superfamily domain; 1,3-beta-glucan synthase subunit FKS1-like, domain-1; Glycosyl transferase, family 48; Major facilitator, sugar transporter-like |
| **PVIT_0017543** | Phyca11_105873, PITG_08806T0, KUF88912, Phal00085, PMUR_0009543, PVIT_0017543 | 2775 | 11.86 | 5.75E-04 | 2.89E-02 | no | calcium-dependent cysteine-type endopeptidase activity; intracellular; cytoplasm; proteolysis | Peptidase C2, calpain, catalytic domain; Peptidase C2, calpain, large subunit, domain III; Peptidase C2, calpain, domain III; Peptidase C2, calpain family |
| **PVIT_0003723** | Phyca11_6928, PITG_15920T0, KUF84589, Phal09845, PMUR_0000189, PVIT_0003723 | 693 | 11.68 | 6.33E-04 | 3.06E-02 | no | integral component of membrane | Transmembrane protein TMEM64 |
| **PVIT_0005563** | Phyca11_35790, PITG_07145T0, KUF77196, Phal04042, PMUR_0000822, PVIT_0005563 | 975 | 11.70 | 6.25E-04 | 3.06E-02 | no | calcium ion binding; peroxisome; fatty acid biosynthetic process; malonyl-CoA decarboxylase activity | Malonyl-CoA decarboxylase, C-terminal |
| **PVIT_0012806** | Phyca11_525423, PITG_19851T0, KUF82130, Phal12053, PMUR_0011389, PVIT_0012806 | 1653 | 11.69 | 6.30E-04 | 3.06E-02 | no | DNA binding | SANT/Myb domain; Myb domain; Homeobox-like domain superfamily |
| **PVIT_0023713** | Phyca11_109643, PITG_05359T0, KUG00654, Phal08368, PMUR_0010774, PVIT_0023713 | 2934 | 11.56 | 6.74E-04 | 3.22E-02 | no | binding | Telomere-associated protein Rif1, N-terminal; Armadillo-type fold |
| **PVIT_0025164** | Phyca11_16573, PITG_13434T0, KUG01795, Phal08065, PMUR_0006374, PVIT_0025164 | 1539 | 11.53 | 6.83E-04 | 3.23E-02 | yes | protein binding; integral component of membrane | Tetratricopeptide-like helical domain superfamily; Sel1-like repeat |
| **PVIT_0008790** | Phyca11_507044, PITG_06514T0, KUF88950, Phal01618, PMUR_0009358, PVIT_0008790 | 447 | 11.02 | 9.02E-04 | 4.21E-02 | no | DNA binding; DNA-directed 5'-3' RNA polymerase activity; calcium ion binding | EF-Hand 1, calcium-binding site; EF-hand domain; RNA polymerase, beta subunit, protrusion; EF-hand domain pair |
| **PVIT_0010265** | Phyca11_510199, PITG_17157T0, KUF64291, Phal14163, PMUR_0005615, PVIT_0010265 | 762 | 10.91 | 9.57E-04 | 4.42E-02 | no | nucleotide binding; nucleic acid binding | RNA recognition motif domain; Nucleotide-binding alpha-beta plait domain superfamily |

| Tested gene | Orthogroup | Align. size | Delta LRT | *P* -value | Q-value | Secreted | GO terms | InterPro domains |
|---|---|---|---|---|---|---|---|---|
| **PVIT_0015981** | Phyca11_551179, PITG_04917T0, KUF80394, Phal14097, PMUR_0007136, PVIT_0015981 | 1530 | 10.83 | 9.99E-04 | 4.56E-02 | no | protein binding; anaphase-promoting complex; regulation of mitotic metaphase/anaphase transition | Cdc23; Tetratricopeptide repeat-containing domain; Tetratricopeptide-like helical domain superfamily; Tetratricopeptide repeat 1; Tetratricopeptide repeat |
| **PVIT_0001226** | Phyca11_506190, PITG_02698T0, KUF84935, Phal08820, PMUR_0001030, PVIT_0001226 | 534 | 10.77 | 1.03E-03 | 4.61E-02 | no | binding | Armadillo-type fold |
| **PVIT_0008576** | Phyca11_525308, PITG_06889T0, KUF91944, Phal11045, PMUR_0007434, PVIT_0008576 | 1887 | 10.77 | 1.03E-03 | 4.61E-02 | no | proteolysis; exocytosis; integral component of membrane; dipeptidase activity | Peptidase C69, dipeptidase A |
| **PVIT_0000445** | Phyca11_548193, PITG_03266T0, KUF69650, Phal04494, PMUR_0004883, PVIT_0000445 | 600 | 10.52 | 1.18E-03 | 5.05E-02 | no | | |
| **PVIT_0000656** | Phyca11_534689, PITG_03458T0, KUG02205, Phal03704, PMUR_0000261, PVIT_0000656 | 840 | 10.50 | 1.19E-03 | 5.05E-02 | no | | Pleckstrin homology domain; ELMO domain; PH-like domain superfamily |
| **PVIT_0007171** | Phyca11_548255, PITG_17258T0, KUF77216, Phal14089, PMUR_0007037, PVIT_0007171 | 1164 | 10.57 | 1.15E-03 | 5.05E-02 | no | GTP binding | GTP binding domain; Uncharacterised GTP-binding protein, C-terminal; P-loop containing nucleoside triphosphate hydrolase |
| **PVIT_0007431** | Phyca11_562149, PITG_02047T0, KUF89726, Phal06708, PMUR_0010426, PVIT_0007431 | 912 | 10.48 | 1.21E-03 | 5.05E-02 | no | epithelial cilium movement; DNA helicase activity; ATP binding; nucleus; DNA repair; DNA recombination; transcription, DNA-templated; heart development; DNA duplex unwinding; ATP-dependent 5'-3' DNA helicase activity; motile cilium assembly; negative regulation of transcription, DNA-templated; digestive tract development; regulation of heart growth; axonemal dynein complex assembly | AAA+ ATPase domain; TIP49, C-terminal; RuvB-like; Nucleic acid-binding, OB-fold; P-loop containing nucleoside triphosphate hydrolase |
| **PVIT_0008480** | Phyca11_503137, PITG_06796T0, KUF86440, Phal05304, PMUR_0006526, PVIT_0008480 | 798 | 10.49 | 1.20E-03 | 5.05E-02 | no | protein kinase activity; ATP binding; protein phosphorylation | Protein kinase domain; Protein kinase-like domain superfamily |
| **PVIT_0010560** | Phyca11_572114, PITG_09827T0, KUF80614, Phal00324, PMUR_0000416, PVIT_0010560 | 705 | 10.52 | 1.18E-03 | 5.05E-02 | no | kinetochore; protein binding; kinetochore microtubule; mitotic spindle assembly checkpoint; phragmoplast; mitotic checkpoint complex; ubiquitin binding; bub1-bub3 complex | WD40-repeat-containing domain; WD40/YVTN repeat-like-containing domain superfamily; WD40 repeat |
| **PVIT_0012401** | Phyca11_18432, PITG_04027T0, KUG01718, Phal08460, PMUR_0007470, PVIT_0012401 | 1299 | 10.44 | 1.24E-03 | 5.12E-02 | no | integral component of membrane | |
| **PVIT_0004375** | Phyca11_503853, PITG_09432T0, KUF76754, Phal05064, PMUR_0007741, PVIT_0004375 | 1440 | 10.31 | 1.32E-03 | 5.32E-02 | no | rRNA modification; protein binding; snoRNA binding; box C/D snoRNP complex; small-subunit processome | Nop domain; NOP5, N-terminal; NOSIC |
| **PVIT_0015139** | Phyca11_41439, PITG_11813T0, KUF76193, Phal08159, PMUR_0004175, PVIT_0015139 | 462 | 10.30 | 1.33E-03 | 5.32E-02 | no | single-organism process | KIF-1 binding protein |
| **PVIT_0018513** | Phyca11_508797, PITG_12489T0, KUF64475, Phal08555, PMUR_0009815, PVIT_0018513 | 1797 | 10.29 | 1.34E-03 | 5.32E-02 | no | nucleus; cytosol; regulation of transcription, DNA-templated; protein methylation; methyltransferase activity; histone-arginine N-methyltransferase activity; peptidyl-arginine methylation, to symmetrical-dimethyl arginine; histone arginine methylation; peptidyl-arginine N-methylation | Protein arginine N-methyltransferase PRMT5; Protein arginine N-methyltransferase |
| **PVIT_0021050** | Phyca11_558170, PITG_13014T0, KUF76541, Phal03980, PMUR_0006766, PVIT_0021050 | 1161 | 10.31 | 1.33E-03 | 5.32E-02 | no | DNA binding; aminopeptidase activity; proteolysis; metallopeptidase activity | Peptidase M24; ArsR-like helix-turn-helix domain; Peptidase M24, methionine aminopeptidase; PA2G4 family |
| **PVIT_0003374** | Phyca11_39056, PITG_05796T0, KUF93703, Phal08266, PMUR_0003949, PVIT_0003374 | 681 | 10.24 | 1.37E-03 | 5.41E-02 | no | helicase activity; nucleolus; methyltransferase activity; S-adenosylmethionine-dependent methyltransferase activity; methylation | Ribosomal RNA processing protein 8 |
| **PVIT_0012102** | Phyca11_530140, PITG_13901T0, KUF87461, Phal10533, PMUR_0006423, PVIT_0012102 | 2223 | 10.20 | 1.41E-03 | 5.48E-02 | no | protein binding | Leucine-rich repeat |

| Tested gene | Orthogroup | Align. size | Delta LRT | $P$-value | Q-value | Secreted | GO terms | InterPro domains |
|---|---|---|---|---|---|---|---|---|
| **PVIT_0003624** | Phyca11_510275, PITG_17786T0, KUF78419, Phal14118, PMUR_0005879, PVIT_0003624 | 1311 | 10.11 | 1.48E-03 | 5.71E-02 | no | catalytic activity; argininosuccinate lyase activity; cytosol; arginine biosynthetic process via ornithine; protein tetramerization | Fumarate lyase, conserved site; Fumarate lyase, N-terminal; Fumarate lyase family; Argininosuccinate lyase; L-Aspartase-like; Fumarase/histidase, N-terminal |
| **PVIT_0000959** | Phyca11_117156, PITG_02419T0, KUF81990, Phal03620, PMUR_0003474, PVIT_0000959 | 1134 | 10.03 | 1.54E-03 | 5.90E-02 | no | ion channel activity; voltage-gated ion channel activity; voltage-gated potassium channel activity; transport; ion transport; potassium ion transport; voltage-gated potassium channel complex; membrane; integral component of membrane; ion transmembrane transport; regulation of ion transmembrane transport; transmembrane transport; potassium ion transmembrane transport | Ion transport domain; Potassium channel domain |
| **PVIT_0003069** | Phyca11_106683, PITG_10252T0, KUG00411, Phal14188, PMUR_0005817, PVIT_0003069 | 414 | 10.00 | 1.57E-03 | 5.94E-02 | no | nucleic acid binding | |
| **PVIT_0000873** | Phyca11_538407, PITG_02256T0, KUF88613, Phal01432, PMUR_0004466, PVIT_0000873 | 1194 | 9.93 | 1.63E-03 | 6.09E-02 | no | catalytic activity; L-aspartate:2-oxoglutarate aminotransferase activity; mitochondrion; cellular amino acid metabolic process; transaminase activity; biosynthetic process; pyridoxal phosphate binding; identical protein binding; L-phenylalanine:2-oxoglutarate aminotransferase activity | Aminotransferases, class-I, pyridoxal-phosphate-binding site; Aminotransferase, class I/classII; Aspartate/other aminotransferase; Pyridoxal phosphate-dependent transferase, major domain; Pyridoxal phosphate-dependent transferase |
| **PVIT_0001204** | Phyca11_545592, PITG_02636T0, KUF88357, Phal07595, PMUR_0001246, PVIT_0001204 | 636 | 9.92 | 1.64E-03 | 6.09E-02 | no | peroxisome | Pex19 protein |
| **PVIT_0016184** | Phyca11_5519, PITG_15748T0, KUF73660, Phal08511, PMUR_0002609, PVIT_0016184 | 837 | 9.85 | 1.70E-03 | 6.25E-02 | no | protein binding; isomerase activity | Tetratricopeptide repeat-containing domain; Peptidyl-prolyl cis-trans isomerase, FKBP-type; Tetratricopeptide-like helical domain superfamily; Tetratricopeptide repeat |
| **PVIT_0004577** | Phyca11_34812, PITG_09792T0, KUF80477, Phal07279, PMUR_0007242, PVIT_0004577 | 1200 | 9.73 | 1.81E-03 | 6.62E-02 | no | nucleic acid binding; ATP-dependent RNA helicase activity; ATP binding; RNA secondary structure unwinding; ion binding | Helicase, C-terminal; DEAD/DEAH box helicase domain; Helicase superfamily 1/2, ATP-binding domain; RNA helicase, DEAD-box type, Q motif; P-loop containing nucleoside triphosphate hydrolase |
| **PVIT_0006926** | Phyca11_571672, PITG_12226T0, KUF86600, Phal01085, PMUR_0008751, PVIT_0006926 | 7878 | 9.66 | 1.88E-03 | 6.80E-02 | no | protein serine/threonine kinase activity; binding; protein binding; ATP binding; nucleus; DNA repair; protein phosphorylation; integral component of membrane; kinase activity; macromolecular complex binding | Phosphatidylinositol 3/4-kinase, conserved site; Phosphatidylinositol 3-/4-kinase, catalytic domain; PIK-related kinase, FAT; FATC domain; FKBP12-rapamycin binding domain; PIK-related kinase; Domain of unknown function DUF3385, target of rapamycin protein; Serine/threonine-protein kinase TOR; Protein kinase-like domain superfamily; Armadillo-like helical; Tetratricopeptide-like helical domain superfamily; Armadillo-type fold |
| **PVIT_0008172** | Phyca11_511439, PITG_09282T0, KUG00452, Phal10033, PMUR_0003742, PVIT_0008172 | 3102 | 9.61 | 1.93E-03 | 6.93E-02 | no | nucleotide binding; ATP binding; endoplasmic reticulum; integral component of plasma membrane; integral component of membrane; cation-transporting ATPase activity; metal ion binding; cation transmembrane transport | Cation-transporting P-type ATPase, N-terminal; Cation-transporting P-type ATPase, C-terminal; P-type ATPase; P-type ATPase, A domain superfamily; HAD superfamily; P-type ATPase, transmembrane domain superfamily; P-type ATPase, cytoplasmic domain N; P-type ATPase, phosphorylation site |
| **PVIT_0015709** | Phyca11_530603, PITG_20263T0, KUF87812, Phal10051, PMUR_0003160, PVIT_0015709 | 477 | 9.50 | 2.05E-03 | 7.29E-02 | no | | |
| **PVIT_0003669** | Phyca11_505826, PITG_15847T0, KUF80597, Phal00683, PMUR_0004281, PVIT_0003669 | 324 | 9.44 | 2.12E-03 | 7.41E-02 | no | integral component of membrane | |
| **PVIT_0006721** | Phyca11_504657, PITG_11215T0, KUF68290, Phal10588, PMUR_0009239, PVIT_0006721 | 1239 | 9.45 | 2.12E-03 | 7.41E-02 | no | DNA binding; transcription factor activity, sequence-specific DNA binding; protein binding; nucleus; regulation of transcription, DNA-templated; membrane; integral component of membrane; sequence-specific DNA binding | DnaJ domain, conserved site; DnaJ domain; Tetratricopeptide repeat-containing domain; Tetratricopeptide-like helical domain superfamily; Tetratricopeptide repeat 1; Tetratricopeptide repeat 2; Tetratricopeptide repeat |

| Tested gene | Orthogroup | Align. size | Delta LRT | *P*-value | Q-value | Secreted | GO terms | InterPro domains |
|---|---|---|---|---|---|---|---|---|
| **PVIT_0014447** | Phyca11_127143, PITG_16078T0, KUF92384, Phal13077, PMUR_0005325, PVIT_0014447 | 3390 | 9.42 | 2.14E-03 | 7.43E-02 | no | protein binding; ligase activity | Tetratricopeptide-like helical domain superfamily; Pentatricopeptide repeat |
| **PVIT_0019916** | Phyca11_507990, PITG_10675T0, KUF86043, Phal09097, PMUR_0008814, PVIT_0019916 | 681 | 9.39 | 2.18E-03 | 7.47E-02 | no | protein kinase activity; protein serine/threonine kinase activity; ATP binding; protein phosphorylation; integral component of membrane | Serine/threonine-protein kinase, active site; Protein kinase, ATP binding site; Protein kinase domain; Protein kinase-like domain superfamily |
| PVIT_0009280 | Phyca11_532120, PITG_03746T0, KUF64635, Phal01178, PMUR_0003371, PVIT_0009280 | 702 | 9.34 | 2.25E-03 | 7.64E-02 | no | phosphatidylinositol binding | Phox homologous domain |
| **PVIT_0004430** | Phyca11_5246, PITG_09508T0, KUF87566, Phal04971, PMUR_0007503, PVIT_0004430 | 903 | 9.29 | 2.31E-03 | 7.72E-02 | no | thiol-dependent ubiquitin-specific protease activity; protein binding; intracellular; nucleus; cytoplasm; ubiquitin-dependent protein catabolic process; shoot system morphogenesis; protein deubiquitination; leaf development | Peptidase C12, ubiquitin carboxyl-terminal hydrolase; Ubiquitinyl hydrolase, UCH37 type |
| **PVIT_0022467** | Phyca11_531502, PITG_12871T0, KUF85714, Phal07421, PMUR_0011340, PVIT_0022467 | 1659 | 9.30 | 2.29E-03 | 7.72E-02 | no | maturation of SSU-rRNA from tricistronic rRNA transcript (SSU-rRNA, 5.8S rRNA, LSU-rRNA); protein binding; nucleolus; snoRNA binding; 90S preribosome; RENT complex; small-subunit processome; rDNA heterochromatin; t-UTP complex; positive regulation of transcription from RNA polymerase I promoter | WD40 repeat, conserved site; WD40-repeat-containing domain; Quinoprotein alcohol dehydrogenase-like superfamily; WD40/YVTN repeat-like-containing domain superfamily; WD40 repeat |
| **PVIT_0006958** | Phyca11_509288, PITG_12191T0, KUG02054, Phal05048, PMUR_0005127, PVIT_0006958 | 1557 | 9.27 | 2.33E-03 | 7.75E-02 | no | | |
| **PVIT_0006664** | Phyca11_504624, PITG_11171T0, KUG00077, Phal11472, PMUR_0003607, PVIT_0006664 | 1305 | 9.14 | 2.51E-03 | 8.25E-02 | no | endosome; endocytosis; vesicle organization; extrinsic component of membrane; phosphatidylinositol binding | Phox homologous domain; AH/BAR domain superfamily |
| **PVIT_0019187** | Phyca11_112787, PITG_08708T0, KUG02168, Phal02063, PMUR_0010310, PVIT_0019187 | 1752 | 9.08 | 2.58E-03 | 8.42E-02 | no | nucleotide binding; nucleic acid binding; RNA binding; transferase activity, transferring glycosyl groups | RNA recognition motif domain; RNA recognition motif domain, eukaryote; Nucleotide-binding alpha-beta plait domain superfamily |
| **PVIT_0018337** | Phyca11_528327, PITG_21219T0, KUF84842, Phal07762, PMUR_0008572, PVIT_0018337 | 1206 | 9.02 | 2.67E-03 | 8.59E-02 | no | nucleotide binding; nucleic acid binding; DNA binding; nucleus; transcription, DNA-templated; regulation of transcription, DNA-templated | RNA recognition motif domain; UPF3 domain; Nucleotide-binding alpha-beta plait domain superfamily |
| **PVIT_0022458** | Phyca11_502634, PITG_12837T0, KUF82068, Phal07425, PMUR_0011413, PVIT_0022458 | 762 | 9.02 | 2.67E-03 | 8.59E-02 | no | nitrogen compound metabolic process; hydrolase activity, acting on carbon-nitrogen (but not peptide) bonds | Carbon-nitrogen hydrolase |
| **PVIT_0021053** | Phyca11_538207, PITG_13017T0, KUF76543, Phal03983, PMUR_0006763, PVIT_0021053 | 3054 | 8.95 | 2.78E-03 | 8.86E-02 | no | nucleotide binding; nucleic acid binding; ATP-dependent RNA helicase activity; helicase activity; ATP binding; nucleus; cytoplasm; RNA processing; ATP-dependent helicase activity; transferase activity | DNA/RNA helicase, ATP-dependent, DEAH-box type, conserved site; Helicase, C-terminal; AAA+ ATPase domain; Helicase-associated domain; DEAD/DEAH box helicase domain; Domain of unknown function DUF1605; Helicase superfamily 1/2, ATP-binding domain; P-loop containing nucleoside triphosphate hydrolase |
| **PVIT_0011532** | Phyca11_526141, PITG_04651T0, KUF81026, Phal03557, PMUR_0002704, PVIT_0011532 | 3273 | 8.90 | 2.86E-03 | 9.04E-02 | no | nuclear export signal receptor activity; binding; nuclear pore; cytoplasm; protein export from nucleus; intracellular protein transport; Ran GTPase binding | Importin-beta, N-terminal domain; Armadillo-like helical; Armadillo-type fold |
| **PVIT_0020049** | Phyca11_502825, PITG_02203T0, KUF76695, Phal00507, PMUR_0003517, PVIT_0020049 | 1713 | 8.87 | 2.90E-03 | 9.11E-02 | no | metal ion binding | FYVE zinc finger; Zinc finger, FYVE-related; Zinc finger, FYVE/PHD-type; Zinc finger, RING/FYVE/PHD-type |
| **PVIT_0010005** | Phyca11_532048, PITG_19361T0, KUF78680, Phal08207, PMUR_0005191, PVIT_0010005 | 1095 | 8.80 | 3.01E-03 | 9.37E-02 | no | DNA binding; nucleus; transcription, DNA-templated; regulation of transcription, DNA-templated | SANT/Myb domain; Myb domain, plants; Myb domain; Homeobox-like domain superfamily |

| Tested gene | Orthogroup | Align. size | Delta LRT | *P* -value | Q-value | Secreted | GO terms | InterPro domains |
|---|---|---|---|---|---|---|---|---|
| **PVIT_0002851** | Phyca11_564360, PITG_10525T0, KUF97566, Phal00241, PMUR_0008019, PVIT_0002851 | 687 | 8.76 | 3.08E-03 | 9.46E-02 | no | nucleic acid binding; metal ion binding | Zinc finger C2H2-type |
| **PVIT_0013001** | Phyca11_7431, PITG_01363T0, KUF97689, Phal01606, PMUR_0001089, PVIT_0013001 | 1236 | 8.76 | 3.08E-03 | 9.46E-02 | no | | |

Orthogroup: orthology group used for the branch-site test. Align. size: size of the alignment for the computation of likelihoods. Delta LRT: test statistic value for the likelihood ratio test. *P* -values were computed considering that the test statistic followed a chi-square distribution with 1 degree of freedom. *Q* -values were computed using a FDR procedure. Genes found expressed in the RNA-seq data are in bold.

Supplementary table 13 *Pl. muralis* genes with a significant signal in the branch-site test

| Tested gene | Orthogroup | Align. size | Delta LRT | *P*-value | *Q*-value | Secreted | GO terms | InterPro domains |
|---|---|---|---|---|---|---|---|---|
| PMUR_0001567 | Phyca11_527201, PITG_05560T0, KUF76730, Phal07792, PMUR_0001567, PVIT_0006127 | 1773 | 142.22 | 8.71E-33 | 3.50E-29 | no | | Transmembrane protein TqsA-like |
| PMUR_0003310 | Phyca11_575261, PITG_16801T0, KUF76457, Phal02727, PMUR_0003310, PVIT_0005623 | 2427 | 140.05 | 2.60E-32 | 5.22E-29 | no | protein binding; nucleus; transcription, DNA-templated; regulation of transcription, DNA-templated | WD40 repeat, conserved site; TUP1-like enhancer of split; WD40-repeat-containing domain; WD40/YVTN repeat-like-containing domain superfamily; WD40 repeat; G-protein beta WD-40 repeat |
| PMUR_0003488 | Phyca11_569416, PITG_02432T0, KUF82000, Phal09734, PMUR_0003488, PVIT_0000975 | 1557 | 133.54 | 6.90E-31 | 9.24E-28 | no | endoplasmic reticulum; gamma-tubulin ring complex; integral component of membrane; transferase activity, transferring hexosyl groups; gamma-tubulin complex localization | Glycosyltransferase, ALG3; Diphthamide synthesis DPH1/DPH2; Mitotic-spindle organizing protein 1 |
| PMUR_0007058 | Phyca11_534782, PITG_17298T0, KUF77837, Phal11291, PMUR_0007058, PVIT_0007143 | 1797 | 131.60 | 1.83E-30 | 1.84E-27 | no | nucleic acid binding; RNA binding; methyltransferase activity; zinc ion binding; tRNA (cytosine-5-)-methyltransferase activity; tRNA methylation | Bacterial Fmu (Sun)/eukaryotic nucleolar NOL1/Nop2p, conserved site; SAM-dependent methyltransferase RsmB/NOP2-type; Zinc finger, CCHC-type; RNA (C5-cytosine) methyltransferase |
| PMUR_0001117 | Phyca11_506369, PITG_01402T0, KUG01891, Phal05882, PMUR_0001117, PVIT_0013032 | 960 | 127.97 | 1.14E-29 | 9.15E-27 | no | protein binding; metal ion binding | Ankyrin repeat-containing domain; Ankyrin repeat |
| PMUR_0004400 | Phyca11_73678, PITG_10460T0, KUF98791, Phal08974, PMUR_0004400, PVIT_0002889 | 4497 | 108.06 | 2.61E-25 | 1.75E-22 | no | metal ion binding | Zinc finger, CCCH-type; P-loop containing nucleoside triphosphate hydrolase |
| PMUR_0005013 | Phyca11_562024, PITG_01941T0, KUF91124, Phal05002, PMUR_0005013, PVIT_0013721 | 1401 | 104.51 | 1.57E-24 | 8.99E-22 | no | DNA binding | SANT/Myb domain; Myb-like domain; Myb domain; Homeobox-like domain superfamily; Leucine-rich repeat, cysteine-containing subtype |
| PMUR_0007681 | Phyca11_102611, PITG_15241T0, KUG01569, Phal02187, PMUR_0007681, PVIT_0009132 | 1506 | 99.92 | 1.59E-23 | 7.96E-21 | yes | | EGF-like, conserved site; EGF-like domain; EGF-like domain, extracellular |
| PMUR_0001531 | Phyca11_117367, PITG_01173T0, KUF79081, Phal08698, PMUR_0001531, PVIT_0010408 | 600 | 95.03 | 1.87E-22 | 8.36E-20 | no | transferase activity | |
| PMUR_0002275 | Phyca11_112853, PITG_07307T0, KUF73082, Phal02050, PMUR_0002275, PVIT_0006848 | 5376 | 81.50 | 1.76E-19 | 7.06E-17 | yes | protein binding | PA14 domain; Galactose-binding-like domain superfamily; Immunoglobulin-like fold; Immunoglobulin E-set; Filamin/ABP280 repeat; Filamin/ABP280 repeat-like |
| PMUR_0007021 | Phyca11_114386, PITG_18010T0, KUF85090, Phal03365, PMUR_0007021, PVIT_0017815 | 951 | 74.68 | 5.53E-18 | 2.02E-15 | no | nucleus | XAP5 protein |
| PMUR_0001949 | Phyca11_117803, PITG_04001T0, KUF77694, Phal06371, PMUR_0001949, PVIT_0017233 | 918 | 70.18 | 5.40E-17 | 1.81E-14 | no | cysteine-type endopeptidase activity; metalloendopeptidase activity; membrane; integral component of endoplasmic reticulum membrane; CAAX-box protein processing | CAAX amino terminal protease |
| PMUR_0006066 | Phyca11_5774, PITG_06715T0, KUF88877, Phal14365, PMUR_0006066, PVIT_0008218 | 891 | 68.03 | 1.61E-16 | 4.98E-14 | no | phosphoprotein phosphatase activity; protein dephosphorylation; hydrolase activity | Calcineurin-like phosphoesterase domain, ApaH type; Serine/threonine-specific protein phosphatase/bis(5-nucleosyl)-tetraphosphatase |
| PMUR_0002613 | Phyca11_5513, PITG_15779T0, KUG00870, Phal08503, PMUR_0002613, PVIT_0016188 | 1308 | 67.78 | 1.83E-16 | 5.25E-14 | no | RNA binding; calcium ion binding | EF-Hand 1, calcium-binding site; DnaJ domain; EF-hand domain; cAMP-dependent protein kinase regulatory subunit, dimerization-anchoring domain; K Homology domain, type 1; EF-hand domain pair |
| PMUR_0007729 | Phyca11_525738, PITG_09447T0, KUF76766, Phal05022, PMUR_0007729, PVIT_0004392 | 3705 | 67.24 | 2.41E-16 | 6.44E-14 | no | peptide alpha-N-acetyltransferase activity; protein binding; N-terminal peptidyl-methionine acetylation; NatB complex | WD40 repeat, conserved site; WD40-repeat-containing domain; Fyv7/TAP26; N-acetyltransferase B complex, non-catalytic subunit; Tetratricopeptide-like helical domain superfamily; WD40/YVTN repeat-like-containing domain superfamily; WD40 repeat |
| PMUR_0005277 | Phyca11_508083, PITG_02503T0, KUF82417, Phal02332, PMUR_0005277, PVIT_0001050 | 846 | 64.99 | 7.53E-16 | 1.89E-13 | no | catalytic activity; metabolic process | Phosphoadenosine phosphosulphate reductase; Rossmann-like alpha/beta/alpha sandwich fold |
| PMUR_0011578 | Phyca11_507107, PITG_06631T0, KUF85240, Phal00047, PMUR_0011578, PVIT_0008718 | 1278 | 63.85 | 1.34E-15 | 3.17E-13 | no | integral component of membrane; phosphatidylinositol binding | Phox homologous domain; Choline transporter-like |
| PMUR_0004875 | Phyca11_115006, PITG_03283T0, KUF69644, Phal01385, PMUR_0004875, PVIT_0000453 | 1197 | 56.71 | 5.05E-14 | 1.13E-11 | yes | thiol-dependent ubiquitin-specific protease activity; protein binding; proteolysis | F-box domain; Peptidase C19, ubiquitin-specific peptidase, DUSP domain |
| PMUR_0001980 | Phyca11_528810, PITG_03943T0, KUF90630, Phal13935, PMUR_0001980, PVIT_0017200 | 321 | 54.22 | 1.79E-13 | 3.78E-11 | no | | |
| PMUR_0010325 | Phyca11_573197, PITG_07841T0, KUF80120, Phal05566, PMUR_0010325, PVIT_0010839 | 1431 | 54.09 | 1.91E-13 | 3.84E-11 | no | nucleotide binding; aminoacyl-tRNA ligase activity; glycine-tRNA ligase activity; ATP binding; cytoplasm; tRNA aminoacylation for protein translation; glycyl-tRNA aminoacylation | Aminoacyl-tRNA synthetase, class II (G/ P/ S/T); Anticodon-binding; Aminoacyl-tRNA synthetase, class II; Glycyl-tRNA synthetase; Glycyl-tRNA synthetase/DNA polymerase subunit gamma-2 |

| Tested gene | Orthogroup | Align. size | Delta LRT | *P*-value | *Q*-value | Secreted | GO terms | InterPro domains |
|---|---|---|---|---|---|---|---|---|
| PMUR_0004180 | Phyca11_41303, PITG_11805T0, KUF79758, Phal02166, PMUR_0004180, PVIT_0015144 | 450 | 51.02 | 9.13E-13 | 1.75E-10 | no | nucleotide binding; nucleic acid binding | RNA recognition motif domain |
| PMUR_0007200 | Phyca11_99433, PITG_06944T0, KUF90117, Phal08541, PMUR_0007200, PVIT_0018116 | 189 | 45.06 | 1.91E-11 | 3.49E-09 | no | | |
| PMUR_0002593 | Phyca11_504150, PITG_15696T0, KUF86113, Phal09989, PMUR_0002593, PVIT_0016169 | 2031 | 43.84 | 3.56E-11 | 6.23E-09 | no | nucleic acid binding; DNA binding; ATP-dependent DNA helicase activity; ATP binding; nucleus; nucleobase-containing compound metabolic process; nucleotide-excision repair; ATP-dependent helicase activity; DNA duplex unwinding | DNA/RNA helicase, ATP-dependent, DEAH-box type, conserved site; Helicase-like, DEXD box c2 type; ATP-dependent helicase, C-terminal; DEAD2; Helical and beta-bridge domain; Helicase superfamily 1/2, ATP-binding domain, DinG/Rad3-type; RAD3/XPD family; ATP-dependent helicase Rad3/Chl1-like; P-loop containing nucleoside triphosphate hydrolase |
| PMUR_0004879 | Phyca11_507482, PITG_03275T0, KUF69635, Phal04866, PMUR_0004879, PVIT_0000449 | 936 | 41.95 | 9.36E-11 | 1.55E-08 | no | ATP binding; nucleus; microtubule-severing ATPase activity; cytoplasmic microtubule organization | ATPase, AAA-type, conserved site; ATPase, AAA-type, core; MIT; Vps4 oligomerisation, C-terminal; P-loop containing nucleoside triphosphate hydrolase |
| PMUR_0005294 | Phyca11_126376, PITG_16118T0, KUF79594, Phal13797, PMUR_0005294, PVIT_0014415 | 876 | 41.89 | 9.63E-11 | 1.55E-08 | no | metal ion binding | WIBG, Mago-binding; Alpha-ketoglutarate-dependent dioxygenase AlkB-like |
| PMUR_0009520 | Phyca11_505136, PITG_08834T0, KUF98811, Phal04384, PMUR_0009520, PVIT_0017554 | 219 | 39.60 | 3.11E-10 | 4.81E-08 | no | RNA binding; ribosome | |
| PMUR_0002542 | Phyca11_11834, PITG_18455T0, KUG01019, Phal06479, PMUR_0002542, PVIT_0014700 | 375 | 39.46 | 3.34E-10 | 4.98E-08 | no | | Complex 1 LYR protein |
| PMUR_0009897 | Phyca11_6670, PITG_17028T0, KUF83453, Phal14352, PMUR_0009897, PVIT_0008810 | 6555 | 38.48 | 5.52E-10 | 7.92E-08 | no | protein binding | cDENN domain; Tetratricopeptide-like helical domain superfamily |
| PMUR_0001897 | Phyca11_84261, PITG_00939T0, KUF78034, Phal02269, PMUR_0001897, PVIT_0007258 | 672 | 38.18 | 6.46E-10 | 8.95E-08 | no | | Diphthamide synthase domain; Endoribonuclease L-PSP/chorismate mutase-like; YjgF/YER057c/UK114 family; Rossmann-like alpha/beta/alpha sandwich fold |
| PMUR_0000837 | Phyca11_20153, PITG_01525T0, KUF77185, Phal10401, PMUR_0000837, PVIT_0005547 | 537 | 37.92 | 7.38E-10 | 9.88E-08 | no | | FAM92 protein |
| PMUR_0005372 | Phyca11_537973, PITG_14622T0, KUF79070, Phal02365, PMUR_0005372, PVIT_0016023 | 630 | 37.47 | 9.28E-10 | 1.20E-07 | no | nucleus; regulation of exit from mitosis | |
| PMUR_0001680 | Phyca11_4263, PITG_12984T0, KUF91955, Phal07113, PMUR_0001680, PVIT_0006460 | 897 | 34.54 | 4.18E-09 | 5.25E-07 | no | integral component of membrane | Tetraspanin; Tetraspanin/Peripherin |
| PMUR_0010436 | Phyca11_532851, PITG_02055T0, KUF89733, Phal09569, PMUR_0010436, PVIT_0007443 | 1092 | 33.94 | 5.67E-09 | 6.90E-07 | no | protein binding; protein folding; prefoldin complex; unfolded protein binding | Tetratricopeptide repeat-containing domain; Prefoldin alpha-like; Tetratricopeptide-like helical domain superfamily; Tetratricopeptide repeat 2; Tetratricopeptide repeat |
| PMUR_0006374 | Phyca11_16573, PITG_13434T0, KUG01795, Phal08065, PMUR_0006374, PVIT_0025164 | 1539 | 33.82 | 6.03E-09 | 7.13E-07 | no | integral component of membrane | Sel1-like repeat |
| PMUR_0009118 | Phyca11_552955, PITG_11907T0, KUF78523, Phal10251, PMUR_0009118, PVIT_0005967 | 465 | 33.41 | 7.47E-09 | 8.57E-07 | no | diphthine synthase activity; metabolic process; methyltransferase activity; peptidyl-diphthamide biosynthetic process from peptidyl-histidine; methylation | Tetrapyrrole methylase; Diphthine synthase; Tetrapyrrole methylase, subdomain 2; Tetrapyrrole methylase, subdomain 1 |
| PMUR_0006492 | Phyca11_531357, PITG_18912T0, KUF84612, Phal13873, PMUR_0006492, PVIT_0017064 | 1797 | 33.25 | 8.08E-09 | 9.02E-07 | no | nuclear-transcribed mRNA catabolic process, nonsense-mediated decay; DNA binding; helicase activity; ATP binding; cytoplasm; zinc ion binding | RNA helicase UPF1, UPF2-interacting domain; P-loop containing nucleoside triphosphate hydrolase |
| PMUR_0008455 | Phyca11_548492, PITG_02924T0, KUF77925, Phal13246, PMUR_0008455, PVIT_0001441 | 1386 | 33.03 | 9.06E-09 | 9.83E-07 | no | | Pentatricopeptide repeat |
| PMUR_0004112 | Phyca11_118922, PITG_00748T0, KUF96721, Phal12777, PMUR_0004112, PVIT_0001940 | 747 | 32.95 | 9.44E-09 | 9.97E-07 | no | maturation of SSU-rRNA from tricistronic rRNA transcript (SSU-rRNA, 5.8S rRNA, LSU-rRNA); binding; transport; integral component of membrane; snoRNA binding; 90S preribosome; small-subunit processome; t-UTP complex; positive regulation of transcription from RNA polymerase I promoter | BP28, C-terminal domain; Major facilitator superfamily domain; Folate-biopterin transporter; U3 small nucleolar RNA-associated protein 10; Armadillo-like helical; Armadillo-type fold |
| PMUR_0005271 | Phyca11_508077, PITG_02526T0, KUF76525, Phal05602, PMUR_0005271, PVIT_0001060 | 3987 | 30.96 | 2.63E-08 | 2.71E-06 | no | microtubule motor activity; ATP binding; microtubule-based movement; ATPase activity; dynein complex | Sigma-54 interaction domain, ATP-binding site 1; AAA+ ATPase domain; Dynein heavy chain domain; ATPase, dynein-related, AAA domain; Dynein heavy chain, domain-1; Dynein heavy chain, domain-2; Dynein heavy chain, AAA module D4; Dynein heavy chain, coiled coil stalk; Dynein heavy chain; P-loop containing nucleoside triphosphate hydrolase |

| Tested gene | Orthogroup | Align. size | Delta LRT | *P* -value | *Q* -value | Secreted | GO terms | InterPro domains |
|---|---|---|---|---|---|---|---|---|
| PMUR_0013122 | Phyca11_8928, PITG_02482T0, KUF89910, Phal04643, PMUR_0013122, PVIT_0001024 | 2658 | 30.71 | 3.00E-08 | 3.01E-06 | no | mRNA splicing, via spliceosome; ATP-dependent RNA helicase activity; helicase activity; ATP binding; spliceosomal complex; cytoplasm | DNA/RNA helicase, ATP-dependent, DEAH-box type, conserved site; Helicase, C-terminal; AAA+ ATPase domain; Helicase-associated domain; Domain of unknown function DUF1605; Helicase superfamily 1/2, ATP-binding domain; P-loop containing nucleoside triphosphate hydrolase |
| PMUR_0002272 | Phyca11_7912, PITG_07319T0, KUF76263, Phal14400, PMUR_0002272, PVIT_0006841 | 2721 | 30.59 | 3.18E-08 | 3.12E-06 | no | GTPase activity; ion channel activity; voltage-gated potassium channel activity; GTP binding; mitochondrial inner membrane; ion transport; potassium ion transport; voltage-gated potassium channel complex; membrane; ribosome biogenesis; transmembrane transport; large conductance calcium-activated potassium channel activity; potassium ion transmembrane transport | Ion transport domain; GTP binding domain; Potassium channel, BK, alpha subunit; GTP-binding protein, orthogonal bundle domain superfamily; P-loop containing nucleoside triphosphate hydrolase |
| PMUR_0004578 | Phyca11_552288, PITG_14171T0, KUF75707, Phal03709, PMUR_0004578, PVIT_0015222 | 1461 | 29.16 | 6.65E-08 | 6.36E-06 | no | amino acid transmembrane transport; integral component of plasma membrane; amino acid transmembrane transporter activity; L-amino acid transmembrane transporter activity; antiporter activity; membrane; L-alpha-amino acid transmembrane transport | Amino acid/polyamine transporter I |
| PMUR_0002160 | Phyca11_529849, PITG_01031T0, KUF89154, Phal07024, PMUR_0002160, PVIT_0009498 | 1803 | 27.87 | 1.29E-07 | 1.21E-05 | no | protein binding | Tetratricopeptide repeat-containing domain; Tetratricopeptide-like helical domain superfamily; Tetratricopeptide repeat 1; Tetratricopeptide repeat |
| PMUR_0002000 | Phyca11_508348, PITG_03940T0, KUF90639, Phal15119, PMUR_0002000, PVIT_0017186 | 183 | 27.65 | 1.45E-07 | 1.32E-05 | no | catalytic activity; integral component of membrane; molybdenum ion binding; pyridoxal phosphate binding | Molybdenum cofactor sulfurase, C-terminal; MOSC, N-terminal beta barrel; Pyruvate kinase-like, insert domain superfamily |
| PMUR_0000736 | Phyca11_503654, PITG_03576T0, KUF64923, Phal09687, PMUR_0000736, PVIT_0000762 | 642 | 27.44 | 1.62E-07 | 1.45E-05 | no | transferase activity | tRNA methyltransferase TRMD/TRM10-type domain; tRNA (guanine-N1-)-methyltransferase, eukaryotic |
| PMUR_0009831 | Phyca11_550767, PITG_12465T0, KUF64516, Phal03922, PMUR_0009831, PVIT_0018529 | 423 | 27.39 | 1.67E-07 | 1.45E-05 | no | adenylate kinase activity; ATP binding; nucleus; ATPase activity; nucleotide phosphorylation | P-loop containing nucleoside triphosphate hydrolase |
| PMUR_0010613 | Phyca11_504513, PITG_01923T0, KUF85018, Phal06254, PMUR_0010613, PVIT_0013712 | 1875 | 25.76 | 3.86E-07 | 3.30E-05 | no | DNA binding; nucleus; transcription, DNA-templated | PWWP domain; Transcription elongation factor S-II, central domain |
| PMUR_0012785 | Phyca11_111612, PITG_13262T0, KUF80949, Phal11076, PMUR_0012785, PVIT_0012692 | 1497 | 25.37 | 4.72E-07 | 3.95E-05 | no | cystathionine beta-synthase activity; cysteine synthase activity; cytoplasm; cysteine biosynthetic process from serine; vesicle docking involved in exocytosis; integral component of membrane; vesicle-mediated transport; cysteine biosynthetic process via cystathionine; pyridoxal phosphate binding | Cysteine synthase/cystathionine beta-synthase, pyridoxal-phosphate attachment site; CBS domain; Pyridoxal-phosphate dependent enzyme; Sec1-like protein; Cystathionine beta-synthase |
| PMUR_0007052 | Phyca11_548231, PITG_17272T0, KUF85373, Phal03255, PMUR_0007052, PVIT_0007149 | 1779 | 25.16 | 5.29E-07 | 4.33E-05 | no | RNA binding; binding | Armadillo-like helical; Armadillo-type fold; Pumilio RNA-binding repeat |
| PMUR_0001201 | Phyca11_556876, PITG_02622T0, KUF85772, Phal06972, PMUR_0001201, PVIT_0001162 | 1920 | 24.66 | 6.83E-07 | 5.49E-05 | no | aspartic-type endopeptidase activity; intracellular; nucleus; proteolysis; zinc ion binding; response to light stimulus; regulation of flower development; membrane; protein catabolic process | B-box-type zinc finger; Aspartic peptidase A1 family; Zinc finger, RING/FYVE/PHD-type; Aspartic peptidase domain superfamily |
| PMUR_0008437 | Phyca11_507649, PITG_02948T0, KUF66946, Phal10416, PMUR_0008437, PVIT_0001408 | 1029 | 23.97 | 9.79E-07 | 7.71E-05 | no | ATP binding; protein folding; response to heat; heat shock protein binding; metal ion binding; unfolded protein binding | DnaJ domain, conserved site; Heat shock protein DnaJ, cysteine-rich domain; DnaJ domain; Chaperone DnaJ, C-terminal; Chaperone DnaJ; HSP40/DnaJ peptide-binding |
| PMUR_0004280 | Phyca11_505827, PITG_15846T0, KUF80590, Phal00682, PMUR_0004280, PVIT_0003668 | 2298 | 23.41 | 1.31E-06 | 1.01E-04 | no | membrane; integral component of membrane | Calcium-dependent channel, 7TM region, putative phosphate; 10TM putative phosphate transporter, cytosolic domain |
| PMUR_0005674 | Phyca11_511802, PITG_23191T0, KUF75376, Phal10091, PMUR_0005674, PVIT_0019782 | 492 | 23.27 | 1.41E-06 | 1.07E-04 | no | mRNA splicing, via spliceosome; RNA binding; protein binding; U1 snRNP; viral nucleocapsid; catalytic step 2 spliceosome | LSM domain, eukaryotic/archaea-type; Tetratricopeptide repeat-containing domain; LSM domain superfamily; Tetratricopeptide-like helical domain superfamily; Tetratricopeptide repeat |
| PMUR_0009083 | Phyca11_120327, PITG_11108T0, KUF76033, Phal00418, PMUR_0009083, PVIT_0006615 | 324 | 22.83 | 1.77E-06 | 1.32E-04 | yes | | LSM domain, eukaryotic/archaea-type; LSM domain superfamily |
| PMUR_0003444 | Phyca11_531591, PITG_02325T0, KUG00912, Phal04997, PMUR_0003444, PVIT_0000931 | 1236 | 22.64 | 1.95E-06 | 1.43E-04 | yes | nucleotide binding; cation transport; membrane; integral component of membrane; hydrolase activity; cation-transporting ATPase activity; metal ion transport; metal ion binding; cation transmembrane transport | Heavy metal-associated domain, HMA |

| Tested gene | Orthogroup | Align. size | Delta LRT | $P$-value | $Q$-value | Secreted | GO terms | InterPro domains |
|---|---|---|---|---|---|---|---|---|
| PMUR_0002428 | Phyca11_529777, PITG_16342T0, KUG00159, Phal05532, PMUR_0002428, PVIT_0011072 | 378 | 22.59 | 2.00E-06 | 1.43E-04 | no | transcription coactivator activity; nucleus; mitochondrion; cytosol; regulation of transcription from RNA polymerase II promoter; membrane; integral component of membrane; glyoxalase III activity; methylglyoxal catabolic process to D-lactate via S-lactoyl-glutathione; lactate biosynthetic process; cellular response to hydrogen peroxide | DJ-1/PfpI |
| PMUR_0002711 | Phyca11_5748, PITG_04644T0, KUG00479, Phal03556, PMUR_0002711, PVIT_0011528 | 1584 | 21.71 | 3.18E-06 | 2.24E-04 | no | mRNA (guanine-N7-)-methyltransferase activity; protein binding; nucleus; mRNA cap binding complex; 7-methylguanosine mRNA capping; RNA (guanine-N7)-methylation | WW domain; mRNA (guanine-N(7))-methyltransferase domain; mRNA (guanine-N(7))-methyltransferase |
| PMUR_0000822 | Phyca11_35790, PITG_07145T0, KUF77196, Phal04042, PMUR_0000822, PVIT_0005563 | 975 | 21.67 | 3.25E-06 | 2.24E-04 | no | fatty acid biosynthetic process; malonyl-CoA decarboxylase activity | Malonyl-CoA decarboxylase, C-terminal |
| PMUR_0009071 | Phyca11_509092, PITG_11127T0, KUF76536, Phal01636, PMUR_0009071, PVIT_0006602 | 1539 | 21.64 | 3.29E-06 | 2.24E-04 | no | transport; retrograde vesicle-mediated transport, Golgi to ER; protein transport; COPI vesicle coat | Coatomer delta subunit; Longin-like domain superfamily |
| PMUR_0009216 | Phyca11_546925, PITG_07262T0, KUF79909, Phal12526, PMUR_0009216, PVIT_0009773 | 2088 | 21.14 | 4.26E-06 | 2.85E-04 | no | | Ribosomal protein L30, conserved site; Ribosomal protein L30, N-terminal; Ribosomal protein L30, ferredoxin-like fold domain; FMP27, C-terminal; Ribosomal protein L7, eukaryotic |
| PMUR_0006586 | Phyca11_526518, PITG_06171T0, KUF77017, Phal07554, PMUR_0006586, PVIT_0002687 | 3594 | 19.83 | 8.47E-06 | 5.57E-04 | no | spliceosomal complex assembly; mRNA binding; binding; U2 snRNP; U12-type spliceosomal complex; U2-type prespliceosome; catalytic step 2 spliceosome | Splicing factor 3B subunit 1; Armadillo-like helical; Armadillo-type fold |
| PMUR_0008667 | Phyca11_551967, PITG_15630T0, KUF94802, Phal09504, PMUR_0008667, PVIT_0019840 | 249 | 19.44 | 1.04E-05 | 6.74E-04 | no | ESCRT II complex; structural molecule activity; protein homodimerization activity; protein targeting to vacuole involved in ubiquitin-dependent protein catabolic process via the multivesicular body sorting pathway | ArsR-like helix-turn-helix domain; ESCRT-II complex, Vps25 subunit; ESCRT-II complex, Vps25 subunit, N-terminal winged helix |
| PMUR_0008085 | Phyca11_546359, PITG_13184T0, KUF81475, Phal04416, PMUR_0008085, PVIT_0016443 | 1263 | 19.20 | 1.18E-05 | 7.51E-04 | no | membrane | Polycystin cation channel, PKD1/PKD2 |
| PMUR_0002375 | Phyca11_561297, PITG_11451T0, KUF91097, Phal05449, PMUR_0002375, PVIT_0005733 | 1938 | 18.80 | 1.45E-05 | 9.11E-04 | no | | |
| PMUR_0003716 | Phyca11_530047, PITG_09305T0, KUG00458, Phal00064, PMUR_0003716, PVIT_0008141 | 585 | 18.72 | 1.51E-05 | 9.35E-04 | no | recombinase activity; four-way junction DNA binding; DNA recombinase assembly; condensed nuclear chromosome; DNA binding; double-stranded DNA binding; single-stranded DNA binding; endodeoxyribonuclease activity; ATP binding; nucleoplasm; DNA repair; mitotic recombination; reciprocal meiotic recombination; DNA-dependent ATPase activity; response to ionizing radiation; strand invasion; chromosome organization involved in meiotic cell cycle | AAA+ ATPase domain; DNA recombination and repair protein Rad51-like, C-terminal; DNA recombination and repair protein RecA-like, ATP-binding domain; DNA recombination and repair protein, RecA-like; P-loop containing nucleoside triphosphate hydrolase |
| PMUR_0003497 | Phyca11_528670, PITG_02447T0, KUF83950, Phal03756, PMUR_0003497, PVIT_0000984 | 666 | 18.69 | 1.54E-05 | 9.38E-04 | no | | NADH:ubiquinone oxidoreductase intermediate-associated protein 30; Galactose-binding-like domain superfamily |
| PMUR_0006334 | Phyca11_110703, PITG_01431T0, KUF81519, Phal14129, PMUR_0006334, PVIT_0013064 | 1644 | 17.56 | 2.78E-05 | 1.66E-03 | yes | carbohydrate metabolic process; beta-glucosidase activity; integral component of membrane; glycosyl compound metabolic process | Glycoside hydrolase family 1; Glycoside hydrolase superfamily |
| PMUR_0008841 | Phyca11_511335, PITG_15139T0, KUF92685, Phal06804, PMUR_0008841, PVIT_0015324 | 2142 | 17.38 | 3.06E-05 | 1.81E-03 | no | nucleotide binding; magnesium ion binding; phospholipid-translocating ATPase activity; ATP binding; plasma membrane; phospholipid transport; integral component of membrane; phospholipid translocation | P-type ATPase; P-type ATPase, subfamily IV; P-type ATPase, A domain superfamily; HAD superfamily; P-type ATPase, transmembrane domain superfamily; P-type ATPase, cytoplasmic domain N; P-type ATPase, phosphorylation site |
| PMUR_0004612 | Phyca11_504053, PITG_14818T0, KUF90246, Phal13482, PMUR_0004612, PVIT_0019686 | 1851 | 17.34 | 3.13E-05 | 1.82E-03 | no | GTPase activity; GTP binding | Guanylate-binding protein/Atlastin, C-terminal; Guanylate-binding protein, N-terminal; P-loop containing nucleoside triphosphate hydrolase |
| PMUR_0001580 | Phyca11_545202, PITG_05541T0, KUF76734, Phal05708, PMUR_0001580, PVIT_0006114 | 933 | 17.21 | 3.35E-05 | 1.92E-03 | no | endoplasmic reticulum membrane; protein N-linked glycosylation; oligosaccharide-lipid intermediate biosynthetic process; integral component of membrane; dolichyl pyrophosphate Man9GlcNAc2 alpha-1,3-glucosyltransferase activity | Glycosyl transferase, ALG6/ALG8 |
| PMUR_0006626 | Phyca11_19592, PITG_11966T0, KUF85566, Phal08576, PMUR_0006626, PVIT_0005945 | 1683 | 17.14 | 3.48E-05 | 1.97E-03 | no | ATP binding; cytoplasm; protein refolding | Chaperonin Cpn60; Chaperonin Cpn60/TCP-1 family; GroEL-like apical domain superfamily; GroEL-like equatorial domain superfamily |

| Tested gene | Orthogroup | Align. size | Delta LRT | *P* -value | *Q* -value | Secreted | GO terms | InterPro domains |
|---|---|---|---|---|---|---|---|---|
| PMUR_0003710 | Phyca11_510133, PITG_09345T0, KUF78834, Phal00069, PMUR_0003710, PVIT_0008133 | 588 | 16.69 | 4.41E-05 | 2.42E-03 | no | ribosomal small subunit assembly; RNA binding; mRNA binding; structural constituent of ribosome; translation; small ribosomal subunit; rRNA binding; cytosolic small ribosomal subunit | Ribosomal protein S7, conserved site; Ribosomal protein S7 domain; Ribosomal protein S5/S7; Ribosomal protein S5/S7, eukaryotic/archaeal |
| PMUR_0005362 | Phyca11_502585, PITG_14639T0, KUF79068, Phal06879, PMUR_0005362, PVIT_0016036 | 969 | 16.70 | 4.37E-05 | 2.42E-03 | no | catalytic activity; transaminase activity; 4-hydroxy-tetrahydrodipicolinate reductase; biosynthetic process; lysine biosynthetic process via diaminopimelate; pyridoxal phosphate binding; oxidation-reduction process | Dihydrodipicolinate reductase, conserved site; Dihydrodipicolinate reductase, N-terminal; Aminotransferase, class I/classII; NAD(P)-binding domain; Dihydrodipicolinate reductase, C-terminal; Pyridoxal phosphate-dependent transferase, major domain; Pyridoxal phosphate-dependent transferase domain 1; Pyridoxal phosphate-dependent transferase |
| PMUR_0004615 | Phyca11_560957, PITG_14806T0, KUF77657, Phal04782, PMUR_0004615, PVIT_0019688 | 591 | 16.54 | 4.77E-05 | 2.59E-03 | no | TRAPP complex; Golgi vesicle transport | Transport protein particle (TRAPP) component; TRAPP I complex, subunit 5; NO signalling/Golgi transport ligand-binding domain superfamily |
| PMUR_0004268 | Phyca11_505833, PITG_15831T0, KUF80602, Phal08044, PMUR_0004268, PVIT_0003655 | 534 | 16.46 | 4.98E-05 | 2.67E-03 | no | ATP binding; mitochondrion; folic acid-containing compound biosynthetic process; 5-formyltetrahydrofolate cyclo-ligase activity; tetrahydrofolate interconversion; metal ion binding | 5-formyltetrahydrofolate cyclo-ligase |
| PMUR_0007242 | Phyca11_34812, PITG_09792T0, KUF80477, Phal07279, PMUR_0007242, PVIT_0004577 | 1200 | 16.37 | 5.20E-05 | 2.75E-03 | no | nucleic acid binding; ATP-dependent RNA helicase activity; ATP binding; RNA secondary structure unwinding; ion binding | Helicase, C-terminal; DEAD/DEAH box helicase domain; Helicase superfamily 1/2, ATP-binding domain; RNA helicase, DEAD-box type, Q motif; P-loop containing nucleoside triphosphate hydrolase |
| PMUR_0006578 | Phyca11_102887, PITG_19877T0, KUF79491, Phal10225, PMUR_0006578, PVIT_0018779 | 1110 | 16.17 | 5.80E-05 | 3.03E-03 | no | GTPase activity; GTP binding | Pleckstrin homology domain; Guanylate-binding protein/Atlastin, C-terminal; Guanylate-binding protein, N-terminal; PH-like domain superfamily; P-loop containing nucleoside triphosphate hydrolase |
| PMUR_0003373 | Phyca11_539851, PITG_03744T0, KUF64622, Phal02084, PMUR_0003373, PVIT_0009278 | 585 | 15.80 | 7.04E-05 | 3.63E-03 | no | protein binding | Bromodomain, conserved site; Bromodomain; NET domain |
| PMUR_0006068 | Phyca11_127589, PITG_06718T0, KUF78798, Phal13655, PMUR_0006068, PVIT_0008220 | 453 | 15.62 | 7.75E-05 | 3.94E-03 | yes | cilium or flagellum-dependent cell motility; microtubule motor activity; axonemal dynein complex; microtubule-based movement; ATPase activity | EGF-like, conserved site; EGF-like domain; EGF-like domain, extracellular |
| PMUR_0000412 | Phyca11_121892, PITG_09835T0, KUF84368, Phal00319, PMUR_0000412, PVIT_0010555 | 1044 | 15.35 | 8.91E-05 | 4.47E-03 | no | protein binding; cytosol; COP9 signalosome; NEDD8-specific protease activity | Proteasome component (PCI) domain; ArsR-like helix-turn-helix domain |
| PMUR_0002201 | Phyca11_538823, PITG_03169T0, KUF83342, Phal12579, PMUR_0002201, PVIT_0000369 | 765 | 15.29 | 9.20E-05 | 4.51E-03 | no | metal ion binding | GPN-loop GTPase; P-loop containing nucleoside triphosphate hydrolase |
| PMUR_0005390 | Phyca11_557901, PITG_14602T0, KUF84861, Phal04182, PMUR_0005390, PVIT_0016004 | 2628 | 15.30 | 9.19E-05 | 4.51E-03 | no | | |
| PMUR_0000189 | Phyca11_6928, PITG_15920T0, KUF84589, Phal09845, PMUR_0000189, PVIT_0003723 | 693 | 15.08 | 1.03E-04 | 4.99E-03 | no | integral component of membrane | Transmembrane protein TMEM64 |
| PMUR_0011101 | Phyca11_15660, PITG_10849T0, KUF80138, Phal11409, PMUR_0011101, PVIT_0017034 | 1290 | 14.95 | 1.10E-04 | 5.27E-03 | no | | |
| PMUR_0011413 | Phyca11_502634, PITG_12837T0, KUF82068, Phal07425, PMUR_0011413, PVIT_0022458 | 762 | 14.93 | 1.12E-04 | 5.27E-03 | no | nitrogen compound metabolic process; hydrolase activity, acting on carbon-nitrogen (but not peptide) bonds | Carbon-nitrogen hydrolase; MORN motif |
| PMUR_0004491 | Phyca11_511651, PITG_12428T0, KUF91253, Phal00893, PMUR_0004491, PVIT_0018586 | 282 | 14.85 | 1.17E-04 | 5.45E-03 | no | nucleus; nucleosome assembly; CAF-1 complex | Chromatin assembly factor 1 subunit A; Armadillo-like helical |
| PMUR_0005574 | Phyca11_531963, PITG_06986T0, KUF97067, Phal10488, PMUR_0005574, PVIT_0018084 | 2154 | 14.82 | 1.18E-04 | 5.45E-03 | no | nucleic acid binding; exonuclease activity; 5'-3' exoribonuclease activity; nucleus; nucleobase-containing compound metabolic process; RNA phosphodiester bond hydrolysis, exonucleolytic | Putative 5-3 exonuclease; 5'-3' exoribonuclease type 2; 5'-3' exoribonuclease |
| PMUR_0011524 | Phyca11_506258, PITG_12916T0, KUF84984, Phal06271, PMUR_0011524, PVIT_0006426 | 1053 | 14.75 | 1.23E-04 | 5.60E-03 | no | cysteine-type endopeptidase activity; extracellular space; lysosome; proteolysis; cysteine-type peptidase activity; membrane; proteolysis involved in cellular protein catabolic process | Cysteine peptidase, cysteine active site; Cysteine peptidase, histidine active site; Cysteine peptidase, asparagine active site; Peptidase C1A, papain C-terminal; Peptidase C1A |

| Tested gene | Orthogroup | Align. size | Delta LRT | $P$-value | $Q$-value | Secreted | GO terms | InterPro domains |
|---|---|---|---|---|---|---|---|---|
| PMUR_0011039 | Phyca11_511466, PITG_21071T0, KUF97922, Phal11133, PMUR_0011039, PVIT_0009159 | 2070 | 14.69 | 1.27E-04 | 5.73E-03 | no | nucleotide binding; RNA binding; aminoacyl-tRNA ligase activity; threonine-tRNA ligase activity; ATP binding; cytoplasm; tRNA aminoacylation for protein translation; threonyl-tRNA aminoacylation; tRNA aminoacylation | Aminoacyl-tRNA synthetase, class II (G/ P/ S/T); TGS; Anticodon-binding; Aminoacyl-tRNA synthetase, class II; Threonyl/alanyl tRNA synthetase, SAD; Threonine-tRNA ligase, class IIa; Beta-grasp domain superfamily; TGS-like; Threonyl/alanyl tRNA synthetase, class II-like, putative editing domain superfamily |
| PMUR_0010795 | Phyca11_552406, PITG_20520T0, KUF74545, Phal11007, PMUR_0010795, PVIT_0008057 | 1191 | 14.03 | 1.80E-04 | 8.03E-03 | no | hydrolase activity | UbiB domain; Protein kinase-like domain superfamily |
| PMUR_0002374 | Phyca11_504193, PITG_11457T0, KUF91107, Phal05450, PMUR_0002374, PVIT_0005734 | 3966 | 13.70 | 2.15E-04 | 9.39E-03 | no | phosphoribosylformylglycinamidine synthase activity; ATP binding; cytoplasm; 'de novo' IMP biosynthetic process; glutamine metabolic process | PurM-like, C-terminal domain; PurM-like, N-terminal domain; Glutamine amidotransferase; Phosphoribosylformylglycinamidine synthase |
| PMUR_0007961 | Phyca11_542166, PITG_02035T0, KUG00299, Phal08830, PMUR_0007961, PVIT_0007421 | 951 | 13.70 | 2.14E-04 | 9.39E-03 | no | protein binding | WD40-repeat-containing domain; Six-bladed beta-propeller, TolB-like; WD40/YVTN repeat-like-containing domain superfamily; WD40 repeat |
| PMUR_0003742 | Phyca11_511439, PITG_09282T0, KUG00452, Phal10033, PMUR_0003742, PVIT_0008172 | 3102 | 13.65 | 2.20E-04 | 9.50E-03 | no | nucleotide binding; ATP binding; endoplasmic reticulum; integral component of plasma membrane; integral component of membrane; cation-transporting ATPase activity; metal ion binding; cation transmembrane transport | Cation-transporting P-type ATPase, N-terminal; Cation-transporting P-type ATPase, C-terminal; P-type ATPase; P-type ATPase, A domain superfamily; HAD superfamily; P-type ATPase, transmembrane domain superfamily; P-type ATPase, cytoplasmic domain N; P-type ATPase, phosphorylation site |
| PMUR_0005881 | Phyca11_125088, PITG_17784T0, KUF78421, Phal06976, PMUR_0005881, PVIT_0003622 | 855 | 13.52 | 2.36E-04 | 1.00E-02 | no | methyltransferase activity; methylation | SAM-dependent methyltransferase RsmB/NOP2-type; RNA (C5-cytosine) methyltransferase |
| PMUR_0005965 | Phyca11_530811, PITG_17370T0, KUF77762, Phal03380, PMUR_0005965, PVIT_0007077 | 1599 | 13.51 | 2.37E-04 | 1.00E-02 | no | vacuole inheritance; cytosol; Golgi to endosome transport; endocytosis; cytoplasmic side of endosome membrane; Rab GTPase binding; endosomal vesicle fusion; metal ion binding | FYVE zinc finger; Zinc finger, FYVE-related; Zinc finger, FYVE/PHD-type; Zinc finger, RING/FYVE/PHD-type; START-like domain superfamily |
| PMUR_0005763 | Phyca11_534052, PITG_21886T0, KUF76579, Phal04857, PMUR_0005763, PVIT_0012793 | 2190 | 12.87 | 3.34E-04 | 1.40E-02 | no | catalytic activity; histidine-tRNA ligase activity; ATP binding; cytoplasm; mitochondrion; histidyl-tRNA aminoacylation; mitochondrial translation | Anticodon-binding; Aminoacyl-tRNA synthetase, class II; Aromatic amino acid lyase; Histidine-tRNA ligase/ATP phosphoribosyltransferase regulatory subunit; Histidine-tRNA ligase; L-Aspartase-like |
| PMUR_0001138 | Phyca11_545851, PITG_01412T0, KUF77091, Phal03341, PMUR_0001138, PVIT_0013050 | 276 | 12.83 | 3.40E-04 | 1.41E-02 | no | ion transport; ion transmembrane transporter activity; membrane; integral component of membrane; ion transmembrane transport; transmembrane transport | Tricarboxylate/iron carrier |
| PMUR_0001623 | Phyca11_527238, PITG_05482T0, KUF80011, Phal09010, PMUR_0001623, PVIT_0006066 | 654 | 12.51 | 4.05E-04 | 1.66E-02 | no | | PUA-like superfamily |
| PMUR_0005464 | Phyca11_511142, PITG_21377T0, KUF82285, Phal13589, PMUR_0005464, PVIT_0011409 | 579 | 12.39 | 4.32E-04 | 1.75E-02 | no | calcium ion binding | EF-Hand 1, calcium-binding site; EF-hand domain; EF-hand domain pair |
| PMUR_0001936 | Phyca11_509604, PITG_00973T0, KUF90323, Phal05387, PMUR_0001936, PVIT_0007287 | 600 | 12.24 | 4.69E-04 | 1.88E-02 | no | | von Willebrand factor, type A; Copine |
| PMUR_0006766 | Phyca11_558170, PITG_13014T0, KUF76541, Phal03980, PMUR_0006766, PVIT_0021050 | 1161 | 12.06 | 5.14E-04 | 2.05E-02 | no | DNA binding; proteolysis; metallopeptidase activity | Peptidase M24; ArsR-like helix-turn-helix domain; PA2G4 family |
| PMUR_0011519 | Phyca11_110421, PITG_12919T0, KUF84986, Phal06266, PMUR_0011519, PVIT_0006431 | 3036 | 12.00 | 5.32E-04 | 2.10E-02 | no | | |
| PMUR_0008412 | Phyca11_543837, PITG_18633T0, KUF82234, Phal13120, PMUR_0008412, PVIT_0020500 | 921 | 11.94 | 5.49E-04 | 2.14E-02 | no | pseudouridine synthesis; RNA binding; RNA processing; tRNA modification; RNA modification; pseudouridine synthase activity; mRNA pseudouridine synthesis | PUA domain; Pseudouridine synthase II, N-terminal; tRNA pseudouridine synthase II, TruB; Pseudouridine synthase, catalytic domain superfamily |
| PMUR_0006895 | Phyca11_504209, PITG_11425T0, KUF82386, Phal13957, PMUR_0006895, PVIT_0009011 | 1974 | 11.53 | 6.83E-04 | 2.61E-02 | no | telomere maintenance via recombination; telomerase activity; protein binding; telomerase holoenzyme complex; RNA-dependent DNA biosynthetic process; nuclear matrix | WD40 repeat, conserved site; Transcriptional repressor Tup1, N-terminal; WD40-repeat-containing domain; Six-bladed beta-propeller, TolB-like; WD40 repeat; G-protein beta WD-40 repeat |
| PMUR_0007560 | Phyca11_503483, PITG_03811T0, KUF85587, Phal06083, PMUR_0007560, PVIT_0009341 | 768 | 11.54 | 6.80E-04 | 2.61E-02 | no | kinase activity; phosphorylation | HAD superfamily |
| PMUR_0001748 | Phyca11_571032, PITG_04715T0, KUF80265, Phal13768, PMUR_0001748, PVIT_0015920 | 960 | 11.34 | 7.57E-04 | 2.87E-02 | no | pantothenate kinase activity; ATP binding; coenzyme A biosynthetic process; integral component of membrane; phosphorylation | Type II pantothenate kinase |
| PMUR_0009594 | Phyca11_508289, PITG_17093T0, KUF79517, Phal06874, PMUR_0009594, PVIT_0010310 | 453 | 11.33 | 7.65E-04 | 2.87E-02 | no | ribosomal large subunit assembly; structural constituent of ribosome; translation; cytosolic large ribosomal subunit; assembly of large subunit precursor of preribosome | Ribosomal protein L24e, conserved site; Ribosomal protein L24e-related |

| Tested gene | Orthogroup | Align. size | Delta LRT | $P$-value | $Q$-value | Secreted | GO terms | InterPro domains |
|---|---|---|---|---|---|---|---|---|
| PMUR_0005817 | Phyca11_106683, PITG_10252T0, KUG00411, Phal14188, PMUR_0005817, PVIT_0003069 | 414 | 11.16 | 8.34E-04 | 3.10E-02 | no | | |
| PMUR_0005400 | Phyca11_36571, PITG_14593T0, KUF76064, Phal13378, PMUR_0005400, PVIT_0015994 | 1278 | 11.11 | 8.60E-04 | 3.17E-02 | no | binding; protein binding | Armadillo-like helical; Armadillo-type fold; Armadillo |
| PMUR_0009691 | Phyca11_509394, PITG_16466T0, KUF74988, Phal11912, PMUR_0009691, PVIT_0013247 | 624 | 11.06 | 8.84E-04 | 3.20E-02 | no | | |
| PMUR_0011370 | Phyca11_5374, PITG_14971T0, KUF76285, Phal04784, PMUR_0011370, PVIT_0004681 | 2307 | 11.06 | 8.82E-04 | 3.20E-02 | no | protein kinase activity; ATP binding; protein phosphorylation; membrane; kinase activity; phosphorylation | Protein kinase domain; Protein kinase-like domain superfamily |
| PMUR_0006889 | Phyca11_571513, PITG_11565T0, KUF75992, Phal06787, PMUR_0006889, PVIT_0009006 | 444 | 10.89 | 9.69E-04 | 3.48E-02 | no | RNA ligase (ATP) activity; ATP binding; tRNA splicing, via endonucleolytic cleavage and ligation | NEDD4-binding protein 2-like 2; P-loop containing nucleoside triphosphate hydrolase |
| PMUR_0006042 | Phyca11_5314, PITG_09682T0, KUF79157, Phal00585, PMUR_0006042, PVIT_0004519 | 606 | 10.84 | 9.92E-04 | 3.50E-02 | no | | |
| PMUR_0012189 | Phyca11_534810, PITG_00845T0, KUF77237, Phal09887, PMUR_0012189, PVIT_0007185 | 1404 | 10.84 | 9.93E-04 | 3.50E-02 | no | 3',5'-cyclic-nucleotide phosphodiesterase activity; protein binding; signal transduction; phosphoric diester hydrolase activity; metal ion binding | 3'5'-cyclic nucleotide phosphodiesterase, conserved site; 3'5'-cyclic nucleotide phosphodiesterase, catalytic domain; GAF domain; HD/PDEase domain |
| PMUR_0001089 | Phyca11_7431, PITG_01363T0, KUF97689, Phal01606, PMUR_0001089, PVIT_0013001 | 1236 | 10.82 | 1.01E-03 | 3.52E-02 | no | | |
| PMUR_0008270 | Phyca11_99088, PITG_06848T0, KUF86793, Phal09162, PMUR_0008270, PVIT_0008522 | 705 | 10.60 | 1.13E-03 | 3.88E-02 | no | RNA methylation; tRNA nucleoside ribose methylation; cytoplasmic translation; cytoplasm; methyltransferase activity; tRNA methyltransferase activity; methylation | Ribosomal RNA methyltransferase FtsJ domain; Ribosomal RNA large subunit methyltransferase E |
| PMUR_0008538 | Phyca11_11261, PITG_00531T0, KUF95640, Phal08382, PMUR_0008538, PVIT_0001747 | 1272 | 10.61 | 1.12E-03 | 3.88E-02 | no | protein-lysine N-methyltransferase activity; peptidyl-lysine monomethylation | Rubisco LSMT, substrate-binding domain |
| PMUR_0001052 | Phyca11_7451, PITG_01313T0, KUF83241, Phal03492, PMUR_0001052, PVIT_0012961 | 975 | 10.58 | 1.14E-03 | 3.89E-02 | no | peroxisomal membrane; peroxisome organization | Peroxisome membrane protein, Pex16 |
| PMUR_0004784 | Phyca11_568079, PITG_03343T0, KUF76147, Phal06948, PMUR_0004784, PVIT_0000531 | 891 | 10.50 | 1.20E-03 | 4.03E-02 | no | | |
| PMUR_0003899 | Phyca11_118786, PITG_00620T0, KUF76974, Phal03586, PMUR_0003899, PVIT_0001830 | 888 | 10.43 | 1.24E-03 | 4.16E-02 | no | nucleotide binding; microtubule motor activity; ATP binding; kinesin complex; microtubule; microtubule-based movement; microtubule binding; ATPase activity | Kinesin motor domain; P-loop containing nucleoside triphosphate hydrolase |
| PMUR_0005922 | Phyca11_4436, PITG_04425T0, KUF76438, Phal11717, PMUR_0005922, PVIT_0016636 | 588 | 10.36 | 1.29E-03 | 4.27E-02 | no | protein polyubiquitination; protein binding; ATP binding; nucleus; cytoplasm; ubiquitin protein ligase binding; proteasome-mediated ubiquitin-dependent protein catabolic process; ubiquitin protein ligase activity | Ubiquitin-conjugating enzyme, active site; Ubiquitin-conjugating enzyme E2; Ubiquitin-associated domain; UBA-like superfamily; Ubiquitin-conjugating enzyme/RWD-like |
| PMUR_0009868 | Phyca11_506559, PITG_13346T0, KUG01598, Phal06319, PMUR_0009868, PVIT_0012642 | 1173 | 10.28 | 1.35E-03 | 4.43E-02 | no | protein binding | IQ motif, EF-hand binding site |
| PMUR_0010618 | Phyca11_106461, PITG_16210T0, KUF76884, Phal13557, PMUR_0010618, PVIT_0013314 | 810 | 10.10 | 1.48E-03 | 4.84E-02 | no | catalytic activity; ATP binding; cellular protein modification process; ligase activity | Tubulin-tyrosine ligase/Tubulin polyglutamylase; Tubulin--tyrosine ligase-like protein 12; ATP-grasp fold, subdomain 1 |
| PMUR_0000724 | Phyca11_540044, PITG_03594T0, KUF64885, Phal09375, PMUR_0000724, PVIT_0000771 | 2784 | 9.95 | 1.61E-03 | 5.17E-02 | no | protein binding | WD40 repeat, conserved site; WD40-repeat-containing domain; WD40/YVTN repeat-like-containing domain superfamily; WD40 repeat |
| PMUR_0002151 | Phyca11_10446, PITG_01043T0, KUF76949, Phal04939, PMUR_0002151, PVIT_0009513 | 1452 | 9.94 | 1.62E-03 | 5.17E-02 | no | protein kinase activity; protein serine/threonine kinase activity; ATP binding; intracellular; protein phosphorylation; peptidyl-serine phosphorylation; intracellular signal transduction | Serine/threonine-protein kinase, active site; Protein kinase, ATP binding site; Protein kinase domain; Protein kinase-like domain superfamily; PH-like domain superfamily |
| PMUR_0013320 | Phyca11_108908, PITG_15417T0, KUG00710, Phal10170, PMUR_0013320, PVIT_0018693 | 720 | 9.93 | 1.62E-03 | 5.17E-02 | no | asparagine synthase (glutamine-hydrolyzing) activity; cytosol; asparagine biosynthetic process; glutamine metabolic process; protein homodimerization activity | Asparagine synthase; Glutamine amidotransferase type 2 domain; Rossmann-like alpha/beta/alpha sandwich fold |
| PMUR_0005360 | Phyca11_557962, PITG_15641T0, KUG00944, Phal07048, PMUR_0005360, PVIT_0016038 | 777 | 9.85 | 1.70E-03 | 5.37E-02 | no | | |
| PMUR_0002730 | Phyca11_83303, PITG_04609T0, KUF79936, Phal02515, PMUR_0002730, PVIT_0011507 | 225 | 9.79 | 1.75E-03 | 5.49E-02 | no | | |
| PMUR_0003509 | Phyca11_502845, PITG_19128T0, KUF77603, Phal09915, PMUR_0003509, PVIT_0020040 | 1257 | 9.74 | 1.80E-03 | 5.62E-02 | no | | DnaJ domain, conserved site; DnaJ domain |
| PMUR_0005937 | Phyca11_558581, PITG_04452T0, KUF79007, Phal04815, PMUR_0005937, PVIT_0016653 | 1212 | 9.63 | 1.92E-03 | 5.93E-02 | no | ATP binding; ATP synthesis coupled proton transport; proton-transporting ATP synthase complex, catalytic core F(1); proton-transporting ATP synthase activity, rotational mechanism; proton-transporting ATPase activity, rotational mechanism | ATPase, F1/V1/A1 complex, alpha/beta subunit, nucleotide-binding domain; P-loop containing nucleoside triphosphate hydrolase |

| Tested gene | Orthogroup | Align. size | Delta LRT | $P$-value | $Q$-value | Secreted | GO terms | InterPro domains |
|---|---|---|---|---|---|---|---|---|
| PMUR_0007380 | Phyca11_528591, PITG_18185T0, KUF87125, Phal07119, PMUR_0007380, PVIT_0019958 | 1413 | 9.46 | 2.10E-03 | 6.43E-02 | no | catalytic activity | Phospholipase D/Transphosphatidylase; Phospholipase D-like domain |
| PMUR_0004545 | Phyca11_128667, PITG_18992T0, KUF90381, Phal01553, PMUR_0004545, PVIT_0018936 | 1221 | 9.42 | 2.14E-03 | 6.52E-02 | no | integral component of membrane; transferase activity, transferring acyl groups other than amino-acyl groups | Acyltransferase 3 |
| PMUR_0003495 | Phyca11_549442, PITG_02445T0, KUF83940, Phal03758, PMUR_0003495, PVIT_0000982 | 651 | 9.40 | 2.17E-03 | 6.54E-02 | no | | |
| PMUR_0005540 | Phyca11_531975, PITG_07024T0, KUG02032, Phal09272, PMUR_0005540, PVIT_0018047 | 1521 | 9.35 | 2.23E-03 | 6.69E-02 | no | nucleotide binding; catalytic activity; IMP dehydrogenase activity; cytoplasm; purine nucleotide biosynthetic process; GMP biosynthetic process; oxidoreductase activity; metal ion binding; oxidation-reduction process | IMP dehydrogenase / GMP reductase, conserved site; CBS domain; IMP dehydrogenase/GMP reductase; Inosine-5'-monophosphate dehydrogenase; Aldolase-type TIM barrel |
| PMUR_0007952 | Phyca11_100998, PITG_14934T0, KUF76276, Phal12359, PMUR_0007952, PVIT_0004686 | 1170 | 9.26 | 2.34E-03 | 6.98E-02 | no | | |
| PMUR_0011522 | Phyca11_533911, PITG_12914T0, KUF84990, Phal06269, PMUR_0011522, PVIT_0006427 | 1797 | 9.21 | 2.41E-03 | 7.11E-02 | no | signal transduction; single-organism cellular process; phosphatidylinositol dephosphorylation | Rho GTPase-activating protein domain; Inositol polyphosphate-related phosphatase; Endonuclease/exonuclease/phosphatase; Rho GTPase activation protein; Immunoglobulin-like fold |
| PMUR_0000424 | Phyca11_121943, PITG_08640T0, KUF97780, Phal01840, PMUR_0000424, PVIT_0010567 | 927 | 9.16 | 2.47E-03 | 7.25E-02 | no | 1-phosphatidylinositol 4-kinase activity; intracellular; plasma membrane; kinase activity; phosphatidylinositol phosphorylation; metal ion binding; phosphatidylinositol-mediated signaling | Phosphatidylinositol 3/4-kinase, conserved site; Phosphatidylinositol 3-/4-kinase, catalytic domain; Pleckstrin homology domain; Phosphatidylinositol kinase; Protein kinase-like domain superfamily; Zinc finger, FYVE/PHD-type; PH-like domain superfamily; Zinc finger, RING/FYVE/PHD-type |
| PMUR_0000516 | Phyca11_534969, PITG_00361T0, KUF77127, Phal02770, PMUR_0000516, PVIT_0001618 | 840 | 9.13 | 2.52E-03 | 7.32E-02 | no | | YAP-binding/ALF4/Glomulin; Glomulin/ALF4 |
| PMUR_0005398 | Phyca11_4109, PITG_14595T0, KUF76069, Phal13376, PMUR_0005398, PVIT_0015996 | 414 | 9.12 | 2.53E-03 | 7.32E-02 | no | | |
| PMUR_0004715 | Phyca11_104632, PITG_00025T0, KUF92809, Phal04027, PMUR_0004715, PVIT_0006794 | 1446 | 9.06 | 2.61E-03 | 7.48E-02 | no | membrane; metal ion binding | Zinc finger, RING-type, conserved site; Zinc finger, RING-type; Zinc finger, RING/FYVE/PHD-type |
| PMUR_0001364 | Phyca11_526284, PITG_10160T0, KUF83843, Phal11516, PMUR_0001364, PVIT_0007458 | 1896 | 9.04 | 2.64E-03 | 7.53E-02 | yes | NADPH-hemoprotein reductase activity; endoplasmic reticulum membrane; FMN binding; integral component of membrane; oxidoreductase activity; [methionine synthase] reductase activity; oxidation-reduction process | Oxidoreductase FAD/NAD(P)-binding; Flavoprotein pyridine nucleotide cytochrome reductase; FAD-binding, type 1; Flavodoxin/nitric oxide synthase; Ferredoxin reductase-type FAD-binding domain; Flavodoxin-like; Riboflavin synthase-like beta-barrel |
| PMUR_0002175 | Phyca11_558908, PITG_03094T0, KUF81709, Phal01416, PMUR_0002175, PVIT_0000339 | 441 | 9.00 | 2.70E-03 | 7.58E-02 | no | integral component of membrane | |
| PMUR_0011785 | Phyca11_566753, PITG_01674T0, KUG00358, Phal10720, PMUR_0011785, PVIT_0005446 | 900 | 9.01 | 2.69E-03 | 7.58E-02 | no | | P-loop containing nucleoside triphosphate hydrolase |
| PMUR_0003036 | Phyca11_131317, PITG_20288T0, KUF95286, Phal10374, PMUR_0003036, PVIT_0012240 | 777 | 8.94 | 2.80E-03 | 7.80E-02 | yes | | |
| PMUR_0006956 | Phyca11_507023, PITG_04970T0, KUF86944, Phal08323, PMUR_0006956, PVIT_0016512 | 2853 | 8.84 | 2.94E-03 | 8.15E-02 | no | | |
| PMUR_0010865 | Phyca11_511575, PITG_12694T0, KUF81275, Phal00713, PMUR_0010865, PVIT_0013480 | 873 | 8.83 | 2.97E-03 | 8.16E-02 | no | 1-alkyl-2-acetylglycerophosphocholine esterase activity; protein binding; intracellular; vacuolar transport; hydrolase activity | WD40-repeat-containing domain; Gamma-secretase aspartyl protease complex, presenilin enhancer-2 subunit; WD40/YVTN repeat-like-containing domain superfamily; WD40 repeat; G-protein beta WD-40 repeat |
| PMUR_0008878 | Phyca11_506576, PITG_15467T0, KUF80863, Phal10188, PMUR_0008878, PVIT_0020102 | 720 | 8.75 | 3.10E-03 | 8.48E-02 | no | | |
| PMUR_0010656 | Phyca11_551908, PITG_15552T0, KUF81933, Phal02609, PMUR_0010656, PVIT_0021188 | 1110 | 8.73 | 3.13E-03 | 8.50E-02 | no | phosphatidylinositol phosphate kinase activity; phosphatidylinositol metabolic process; phosphatidylinositol phosphorylation | Phosphatidylinositol-4-phosphate 5-kinase, core; Phosphatidylinositol-4-phosphate 5-kinase; Phosphatidylinositol-4-phosphate 5-kinase, C-terminal; Phosphatidylinositol-4-phosphate 5-kinase, N-terminal |
| PMUR_0006542 | Phyca11_539182, PITG_06792T0, KUF86444, Phal07324, PMUR_0006542, PVIT_0008465 | 2760 | 8.63 | 3.30E-03 | 8.90E-02 | no | nucleic acid binding; DNA binding; ATP-dependent DNA helicase activity; ATP binding; nucleus; nucleobase-containing compound metabolic process; ATP-dependent helicase activity; zinc ion binding; DNA duplex unwinding | DNA/RNA helicase, ATP-dependent, DEAH-box type, conserved site; Zinc finger, RING-type; Helicase-like, DEXD box c2 type; ATP-dependent helicase, C-terminal; DEAD2; Helicase superfamily 1/2, ATP-binding domain, DinG/Rad3-type; ATP-dependent helicase Rad3/Chl1-like; Zinc finger, RING/FYVE/PHD-type; P-loop containing nucleoside triphosphate hydrolase |

| Tested gene | Orthogroup | Align. size | Delta LRT | $P$-value | $Q$-value | Secreted | GO terms | InterPro domains |
|---|---|---|---|---|---|---|---|---|
| PMUR_0009488 | Phyca11_11491, PITG_16813T0, KUF96155, Phal10207, PMUR_0009488, PVIT_0022045 | 1395 | 8.59 | 3.38E-03 | 9.04E-02 | no | protein serine/threonine kinase activity; ATP binding; protein phosphorylation | RIO kinase, conserved site; RIO kinase; Serine/threonine-protein kinase Rio1; Protein kinase-like domain superfamily |
| PMUR_0011335 | Phyca11_538054, PITG_12876T0, KUF84254, Phal07415, PMUR_0011335, PVIT_0022472 | 471 | 8.55 | 3.45E-03 | 9.18E-02 | no | nucleotide binding; nucleic acid binding | RNA recognition motif domain |
| PMUR_0005097 | Phyca11_18948, PITG_17384T0, KUF87775, Phal09280, PMUR_0005097, PVIT_0019364 | 876 | 8.53 | 3.49E-03 | 9.23E-02 | no | protein binding | Tetratricopeptide repeat-containing domain; Tetratricopeptide-like helical domain superfamily; Tetratricopeptide repeat |
| PMUR_0009825 | Phyca11_550750, PITG_12479T0, KUF64498, Phal02835, PMUR_0009825, PVIT_0018522 | 399 | 8.48 | 3.59E-03 | 9.43E-02 | no | | |
| PMUR_0004175 | Phyca11_41439, PITG_11813T0, KUF76193, Phal08159, PMUR_0004175, PVIT_0015139 | 462 | 8.44 | 3.67E-03 | 9.57E-02 | no | mitochondrion; mitochondrial transport; nervous system development; kinesin binding | KIF-1 binding protein |
| PMUR_0011035 | Phyca11_529546, PITG_21222T0, KUF97925, Phal11965, PMUR_0011035, PVIT_0009155 | 2226 | 8.39 | 3.77E-03 | 9.78E-02 | no | intracellular protein transport; retrograde vesicle-mediated transport, Golgi to ER; Golgi organization; Golgi transport complex | Conserved oligomeric Golgi complex subunit 7 |
| PMUR_0007898 | Phyca11_509876, PITG_05224T0, KUG01522, Phal04316, PMUR_0007898, PVIT_0001977 | 423 | 8.33 | 3.91E-03 | 1.00E-01 | no | microtubule motor activity; catalytic activity; 4-aminobutyrate transaminase activity; protein binding; ATP binding; kinesin complex; microtubule-based movement; microtubule binding; transaminase activity; gamma-aminobutyric acid metabolic process; ATPase activity; pyridoxal phosphate binding; phosphatidylinositol dephosphorylation | C2 domain; Inositol polyphosphate-related phosphatase; Kinesin motor domain; Endonuclease/exonuclease/phosphatase; 4-aminobutyrate aminotransferase, eukaryotic; Aminotransferase class-III; SMAD/FHA domain superfamily; Pyridoxal phosphate-dependent transferase, major domain; Pyridoxal phosphate-dependent transferase domain 1; Pyridoxal phosphate-dependent transferase; P-loop containing nucleoside triphosphate hydrolase |
| PMUR_0011824 | Phyca11_121654, PITG_15623T0, KUF94800, Phal09498, PMUR_0011824, PVIT_0009166 | 1629 | 8.34 | 3.89E-03 | 1.00E-01 | no | mRNA splicing, via spliceosome; RNA binding; binding; protein binding; U2-type catalytic step 1 spliceosome; catalytic step 2 spliceosome | MIF4G-like, type 3; Initiation factor eIF-4 gamma, MA3; Armadillo-type fold |

Orthogroup: orthology group used for the branch-site test. Align. size: size of the alignment for the computation of likelihoods. Delta LRT: test statistic value for the likelihood ratio test. $P$-values were computed considering that the test statistic followed a chi-square distribution with 1 degree of freedom. $Q$-values were computed using a FDR procedure.
